# A supernumerary designer chromosome for modular *in vivo* pathway assembly in *Saccharomyces cerevisiae*

**DOI:** 10.1101/2020.02.18.954131

**Authors:** Eline D. Postma, Sofia Dashko, Lars van Breemen, Shannara K. Taylor Parkins, Marcel van den Broek, Jean-Marc Daran, Pascale Daran-Lapujade

## Abstract

The construction of microbial cell factories for sustainable production of chemicals and pharmaceuticals requires extensive genome engineering. Using *Saccharomyces cerevisiae*, this study proposes Synthetic Chromosomes (SynChs) as orthogonal expression platforms for rewiring native cellular processes and implementing new functionalities. Capitalizing the powerful homologous recombination capability of *S. cerevisiae*, modular SynChs of 50 and 100 Kb were fully assembled *de novo* from up to 44 transcriptional-unit-sized fragments in a single transformation. These assemblies were remarkably efficient and faithful to their *in silico* design. SynChs made of non-coding DNA were stably replicated and segregated irrespective of their size without affecting the physiology of their host. These non-coding SynChs were successfully used as landing pad and as exclusive expression platform for the essential glycolytic pathway. This work pushes the limit of DNA assembly in *S. cerevisiae* and paves the way for *de novo* designer chromosomes as modular genome engineering platforms in *S. cerevisiae*.

## INTRODUCTION

Microbial cell factories have an important role to play in the development of a sustainable and environmentally friendly biobased economy, as already exemplified by the ca. 100 billion litres of bioethanol (1) and half of the world’s insulin (2) annually produced by *Saccharomyces cerevisiae*, more commonly known as baker’s yeast. However the construction of novel microbial production hosts for chemicals and pharmaceuticals is impeded by the poor economic viability of bio-based processes. Improving microbial process profitability and competition with current (petrochemical based) production methods, requires powerful microbial cell factories that produce (novel) chemicals from non-native feedstock (3) with high product yields and productivity in harsh industrial conditions. This can only be accomplished by extensive genome engineering that not only adds new functionalities to the host microbe, but also deeply rewires its native cellular and metabolic processes (4,5).

Although the construction of synthetic cell factories with designer genomes, tailor-made for the optimum production of certain chemicals, might become feasible in the future, it is currently far from reach. One reason is our still limited understanding for the absolute requirements of life, as demonstrated by the recent reconstruction of a minimal *Mycoplasma* genome in which one third of the essential genes are not functionally characterized (6). The second major hurdle in the synthesis of designer genomes is DNA synthesis. The maximum size for the chemical synthesis of ssDNA of a quality that can be used for biotechnological application is 200 bp (7,8). Despite recent developments in stitching together these ssDNA parts into genes, routine synthesis of designer genomes would be too cumbersome and expensive (8,9). Indeed, although numerous fast (seamless) *in vitro* assembly methods (10) have been developed such as: Gibson assembly, Golden Gate, ligase cycling reaction, seamless ligation cloning extract and circular polymerase extension cloning, these methods are intrinsically limited in the number of fragments that can be assembled as well as the final size of the assembled DNA. Furthermore, *Escherichia coli*, used as a propagation host (11), cannot faithfully replicate genome-size DNA constructs (12). It is *S. cerevisiae* that ultimately enabled the complete assembly and replication of the 580 Kb *Mycoplasma* genome (12). The homologous recombination machinery of *S. cerevisiae* was able to assemble 25 overlapping DNA fragments of approximately 24 kb into the *Mycoplasma* genome as well as assemble 38 overlapping ssDNA pieces of 200 bp (13,14). This high efficiency and fidelity of homologous recombination has undoubtedly contributed to *S. cerevisiae* popularity as eukaryotic model and cell factory. Yet, the full extent of *in vivo* assembly capabilities in terms of size and number of fragments has not been explored. Introduction of large genetic constructs in *S. cerevisiae* typically takes place by integration in existing chromosomes, without much considerations for potential impact on the host chromosome architecture. The recent introduction of the noscapine pathway, for example, required introduction of 31 (heterologous) genes in nine different chromosomal loci (4). Integration of large constructs might alter the chromosome structure and affect the expression of genes surrounding the newly added construct, as well as replication of the host chromosome. It might also trigger unwanted chromosomal rearrangements between native chromosomes. It is long known that *S. cerevisiae* can stably replicate and segregate Yeast Artificial Chromosomes (YACs) (15). In the present study, we explore the potential of *in vivo* assembled, supernumerary chromosomes for modular genome engineering in *S. cerevisiae*.

To enable easy remodelling of existing functions, Kuijpers *et al.* (16) developed the pathway swapping concept in *S. cerevisiae*. The genetic reduction and relocalization of the entire Embden-Meyerhoff-Parnas pathway of glycolysis to a single chromosomal locus enabled to swap this essential pathway to any new (heterologous) design in two simple steps (16,17). Combined to an orthogonal expression platform, in the form of a supernumerary Synthetic Chromosome (SynCh), pathway swapping would offer the possibility to make the introduction and expression of large product pathways in combination with the rewiring of essential native pathways, more efficient and predictable. Unlike the construction of chromosomes based on an existing structure for the *Mycoplasma* and *S. cerevisiae* genome projects (Sc2.0) (18,19), the SynCh assembly design should be based on modularity to allow for easy change in makeup and configuration of native and heterologous pathways.

The present study explores the possibility for a modular assembly of a designer, supernumerary SynCh for rational engineering of the yeast genome. A pipeline was developed for easy and rapid design, *in vivo* assembly and verification of SynChs. The limits of *in vivo* assembly in terms of number and size of assembled DNA fragments, efficiency and fidelity were explored. The stability of the Synchs and their impact on yeast physiology were probed. Finally the ability of supernumerary synthetic chromosomes to serve as landing pad for metabolic pathways and to carry essential pathways was investigated. This study paves the way for the implementation of supernumerary, synthetic chromosomes as modular expression platforms for native and heterologous functions.

## MATERIALS AND METHODS

Detailed information on materials and methods is provided in the supplementary data and includes the following: strains, maintenance and growth media, molecular biology techniques, plasmid construction, strain construction, experimental quantification of SynCh assembly efficiency, fluorescence detection by microscopy and flow cytometry, contour-clamped homogeneous electric field (CHEF) electrophoresis, sequencing, physiological characterization, SynCh stability determination, in vitro enzyme activities and statistical analysis

## RESULTS

### Design considerations and proof of concept for the modular, *de novo* assembly of supernumerary synthetic chromosomes

Establishment of a workflow for fast and flexible construction of SynChs should include a set a rules to enable easy design, assembly and screening of SynChs with different configurations of genes and auxiliary parts. This flexibility, required for pathway optimization in future strain construction programs, gave the possibility to explore structural requirements for the assembly and maintenance of the SynChs in the present study. Indeed, in this nascent field, very little is known about SynChs design requirements and even less is known about SynChs stability and impact on physiology. The second most important construction criterion was the minimization of experimental steps to construct the SynChs. Accordingly, the chosen construction workflow exploits the strengths of *in vitro* and *in vivo* assembly, by first using Golden gate cloning for stitching functional DNA parts together (e.g. construction of transcription units from promoter, gene and terminator), then assembling *de novo* these *in vitro* stitched parts using *S. cerevisiae*’s highly efficient homologous recombination (Fig. 1A). This workflow also minimized the number of PCR amplification steps, that are notoriously error-prone, to a single one.

**Figure 1:**
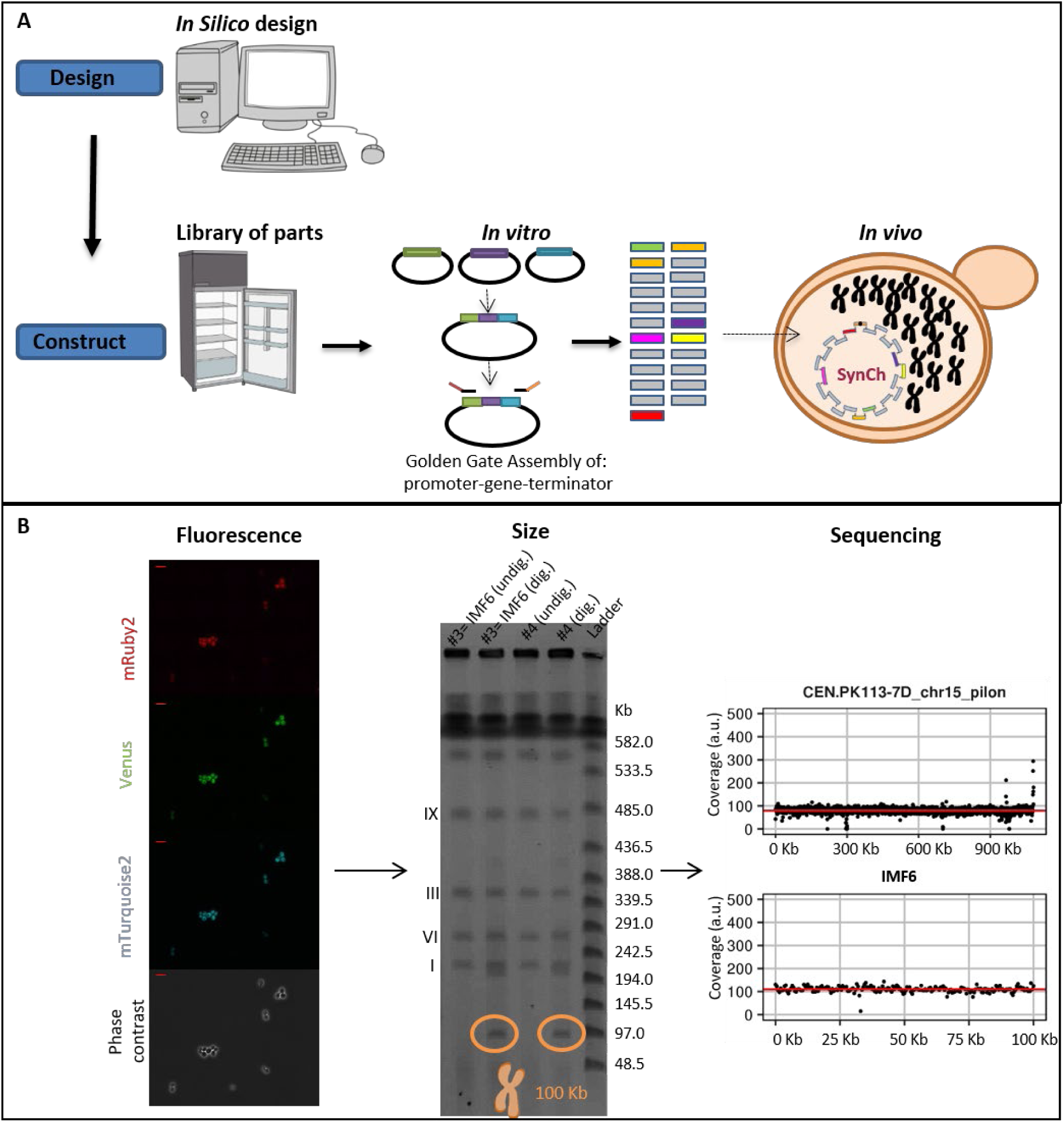
Modular genome engineering strategy. Schematic representation of the design and construction of SynChs (**A**), as well as screening based on fluorescence, size and sequencing (**B**). The data in panel B represent the results pertaining to the assembly of the 100 Kb SynCh in strain IMF6. The red bar in the top left of the fluorescence microscopy pictures (40x) represents 10 µm. The CHEF gel used to determine the size of the SynCh contains two transformants (#3 and #4) of the 100 Kb SynCh transformation. The SynChs are either undigested (undig.) by I-SceI and therefore in their circular form, or in-plug linearized by I-SceI digestion (dig.) and therefore in linear form, which can be visualised on CHEF gel.

In *vivo* SynCh assembly was promoted by framing the DNA fragments transformed to yeast with short overhangs with no homology with the yeast genome (60bp long called SHRs (20)), that can be easily and cheaply added by PCR. The SynChs were designed to carry replicative fragments, including a centromere and autonomously replicating sequences (ARS) spaced every 30-40 Kb (21), and markers to facilitate the selection of transformants with correctly assembled SynChs. To evaluate the efficiency of our SynCh construction workflow, two test chromosomes of 50 and 100 Kb were designed and assembled. To mimic assembly of pathways, typically composed of transcription units of ca. 2 to 3 Kb, but to avoid potential interference by gene expression from the SynChs, the test SynChs were assembled from 2.5 Kb non-coding *E. coli* DNA fragments. Specifically the test SynChs comprised: *CEN6/ARS4(22), ARS1* (and also ARS417 for 100 Kb SynCh), two auxotrophic selection markers (*HIS3* and *URA3*), three fluorescent proteins (Venus, mRuby2 and mTurquoise2), and 2.5 Kb non-coding *E. coli* DNA fragments used to reach the desired chromosome size (called filler fragments) and the telomerator (23). This resulted in 23 fragments for the assembly of the 50 kb SynCh and 44 fragments for the 100 Kb SynCh (Fig. 2), which were transformed in a fixed molar amounts per fragment (therefore approximately twice as much DNA was transformed for the 100 Kb SynCh with respect to the 50 Kb SynCh). Despite the large number of DNA fragments, transformation on selective growth medium (histidine and uracil deficient) resulted in a large number of colonies (ca. 2000 colonies and 275 colonies, for the 50 Kb and 100 Kb SynCh respectively). Four colonies of each transformation were checked by fluorescence microscopy and all expressed the three fluorescent markers (Fig. 1B and Suppl. Fig. S1). For both test chromosomes, one of these four colonies was randomly selected and tested by diagnostic PCR of the assembly junctions (Suppl. Fig. S2), which revealed the correct assembly of the 50 and 100 Kb SynChs. The genome of two transformants harbouring the 100 Kb SynCh (IMF1 and IMF6) and one transformant harbouring the 50 Kb chromosome (IMF2) were sequenced. While IMF2 and IMF6 were faithfully assembled, harbouring all fragments of the *in silico* design in the correct configuration, IMF1 was not. According to the sequencing data, IMF1 carried all expected fragments but was harbouring an internal deletion in filler fragment (6C) and a duplication of a 35 Kb region (Suppl. Fig. S3). This duplication, covering the middle of *mTurquoise2* to the middle of *Venus*, was confirmed by karyotyping using in-plug linearization of the SynCh (Suppl. Fig. S4-S5) and most likely resulted from a recombination event between the *mTurquoise2* and the *Venus* genes, which have a nucleotide identity of 97% (24-26). SynChs can therefore be efficiently and faithfully assembled from as many as 44 fragments, despite the presence of an internal homologous region of approximately 700 bp, as well as regions homologous to the native chromosomes (*pTEF1, pTEF2, pCCW12, tSSA1, ARS, HIS3* and *URA3* expression cassette). The 60 bp SHRs were therefore sufficient to promote the efficient assembly of the SynChs.

**Figure 2:**
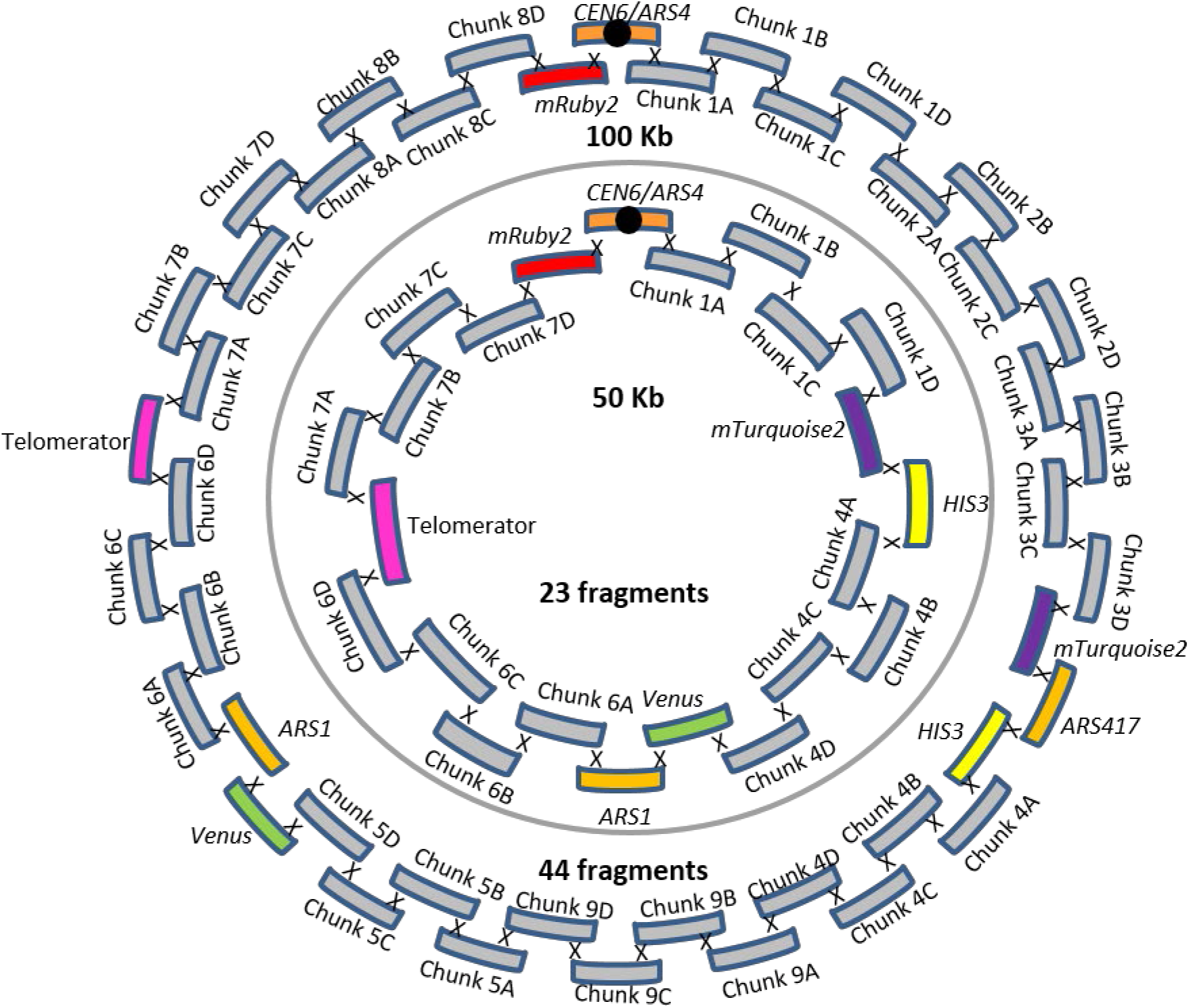
50 and 100 kb SynCh designs. The outer circle depicts the *in vivo* assembly of the 100 Kb SynCh from 44 fragments present in strain IMF6. The inner circle depicts the *in vivo* assembly of the 50 Kb SynCh from 23 fragments present in strain IMF2.

### Exploring optimal fragment size and number for efficient, modular *in vivo* assembly of SynChs

Although the assembly of the 50 Kb and 100 Kb SynChs resulted in a large number of colonies, doubling the number of transformed fragments caused a seven-fold decrease in colony count. This suggested that the number of fragments in the SynCh design might become a limiting factor for the construction of larger SynChs harbouring many native and non-native pathways. To gain more insight into transformation efficiency (i.e. CFU/10^8^ transformed cells) and to test to what extent efficiency might be influenced by fragment size and/or the number of transformed fragments, new 50 and 100 Kb SynChs were designed and assembled using DNA filler fragments of three different sizes: 2.5 Kb, 5Kb and 10 Kb, transformed in equimolar amounts. The transformations resulted in a large number of colonies (10^3^-10^4^ upon plating of the whole transformation mixture), and was most efficient with 5 Kb fragments. On average transformation with 5 Kb fragments was four-fold more efficient than 2.5 Kb fragments and three-fold more efficient than with 10 Kb fragments (one-tailed paired t-tests p ≤ 0.05), but fragments of 2.5 Kb and 10 Kb led to the same number of transformants (one-tailed paired t-test p > 0.05) (Fig. 3A). For each specific fragment size, doubling the number of fragments to assemble significantly decreased transformation efficiency by ca. 3-fold (2-tailed paired t-test p < 0.01) (Fig. 3A). Therefore, for the assembly of large SynChs it is advisable to reduce the number of fragments to transform by connecting together transcription units (e.g. via fusion PCR or Golden Gate cloning).

**Figure 3:**
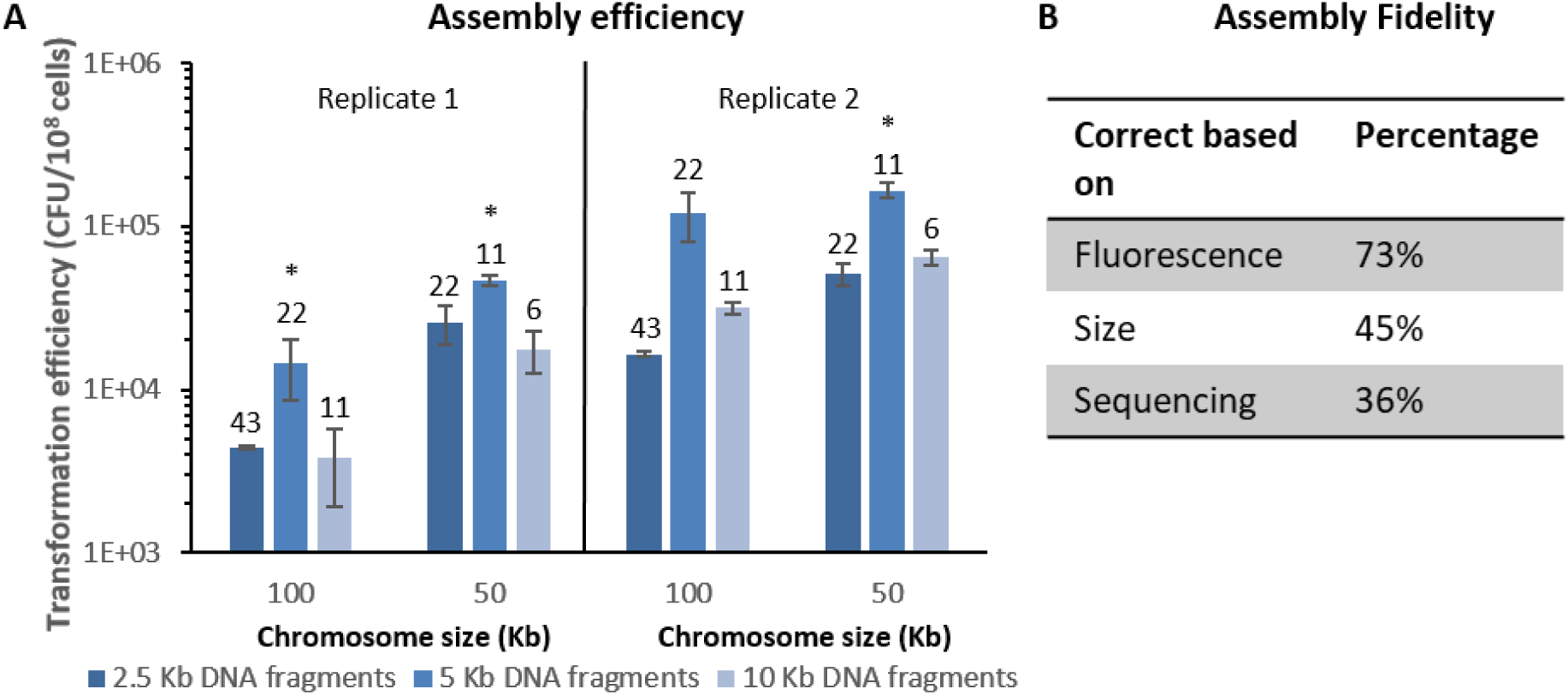
Transformation efficiency and fidelity. **A**) Transformation efficiencies for the *in vivo* assembly of 100 kb and 50 kb synthetic chromosomes. Both SynChs have been assembled with 2.5 Kb (dark blue), 5 Kb (lighter blue) and 10 Kb (lightest blue) fragments. The number above each bar indicates the number of assembled fragments. Data were normalized to 10^8^ cells using controls on YPD plates, performed in biological duplicates (replicate 1 and 2) in technical triplicates for replicate 1 and technical duplicates for replicate 2. The asterisks indicates that transformation with 5 Kb fragments resulted in a significantly different transformation efficiency compared to both neighbouring efficiencies (two-tailed paired homoscedastic t-test p<0.05). **B**) Assembly fidelity. *In vivo* assembly of the 100 Kb Syn12 from 43 fragments. Eleven colonies that grew on selective medium were tested for mRuby2 and mTurquoise2 fluorescence. SynChs were subsequently screened based on their size on CHEF gel and lastly by Illumina sequencing. The percentage of correct colonies for each screening round is indicated.

Next to transformation efficiency, fidelity of assembly (i.e. percentage of the transformants harbouring all transformed fragments in the correct order) is of paramount importance for functional pathway construction. Verification of the SynChs assembly by PCR, as performed in most other synthetic chromosome assemblies (14,18), is not only too labour intensive considering the large number of assembled fragments, but it is also unreliable. Indeed, PCR erroneously indicated a correct configuration for IMF1. Sequencing dozens of chromosomes by next generation sequencing is also time-consuming and costly. We therefore set up a screening pipeline consisting in a first selection based on the presence of auxotrophic markers distributed on the SynCh, followed by fluorescence-activated cell sorting (FACS) of transformants expressing the fluorescent markers also distributed evenly on the SynCh, the subsequent filtering based on chromosome size by Karyotyping and a final confirmation of correct assembly by sequencing (Fig. 1B).

To test the fidelity of assembly and assess the ease of our screening pipeline a novel SynCh design was made (Suppl. Fig. S6). The 100 Kb Syn12 chromosome was similar to 100 Kb SynCh carried by IMF6, except that only two fluorescent markers, *mRuby2* and *mTurquoise2*, were included, to prevent unwanted recombinations between the highly similar *mTurquoise2* and *Venus* genes, and the strongly expressed *mRuby2* was relocated away from the centromere. From the transformation of 43 fragments of ∼2.5 Kb, eleven colonies were screened for correct SynCh assembly. Eight (73%, Fig. 3B and Suppl. Fig. S7) showed double fluorescence, of which five displayed the correct SynCh size by karyotyping (45% of all transformants, Suppl. Fig. S8). Of these five correctly sized SynChs, four proved to be correctly assembled by Illumina whole genome sequencing (Suppl. Fig. S3). Therefore a remarkably high assembly fidelity of 36% was reached when assembling 43 fragments in *S. cerevisiae*. It should be noted however that although sequencing data showed coverage for all fragments in the four correct colonies, two showed uniform coverage of all fragments (Syn12.4 and Syn12.8) while in the other two, the coverage of a small number of fragments differed from the rest of the SynCh (four and two *E.coli* filler fragments with lower coverage for Syn12.1 and Syn12.3 respectively). This coverage variation most likely results from population heterogeneity, with subpopulations carrying different SynChs configurations. This heterogeneity might have occurred early during SynCh assembly, even though the strains were selected after single colony isolation, or later by recombination events occurring during strain propagation. Sequencing of IMF2 and IMF6, carrying the 50 Kb and 100 Kb original SynCh design, confirmed a uniform coverage for all fragments in these strains.

It is important that assembly of the synthetic chromosomes does not result in a large number of mutations in the native genome nor in the assembled synthetic chromosomes. Sequencing data revealed that the assembly of the SynChs resulted in a low number of mutations in the native genome as well as in the assembled SynCh. The four correctly assembled SynChs: Syn12.1 (IMF24), Syn12.3 (IMF25), Syn12.4 (IMF26) and Syn12.8 (IMF23) contained 3, 1, 13 and 12 SNPs in the SynCh, respectively (between 0% and 58% of all mutations identified in the SynChs were located in SHRs, most likely originating from amplification primers(20)); and the native genome contained 1, 0, 2 and 0 mutations in coding regions, respectively (Suppl. Table S1-S2). This was similar to the previously constructed IMF2 (50 Kb) and IMF6 (100 Kb) strain which only harbored no and a single mutation in coding regions of the native genome; and 5 and 16 mutations in the SynCh, respectively (Suppl. Table S1-S2).

### Application of synthetic chromosomes as DNA landing pads

Supernumerary SynChs can be attractive landing pads to add, remove or modify functionalities. While recent chromosome engineering efforts have demonstrated that native *S. cerevisiae* chromosomes are extremely robust to large changes in their size and number (27,28), very little is known about *S. cerevisiae*’s tolerance to *in vivo* synthetic chromosome editing (19). The 50 Kb and 100 Kb SynChs carried by strains IMF2 and IMF6 respectively, were used as landing pads for the integration of an additional 35 Kb of non-coding DNA (Fig. 4). *mTurquoise2* was targeted by CRISPR-Cas9 for introducing a double strand break and replaced by the 35 Kb construct *in vivo*-assembled from 16 DNA fragments, resulting in two chromosomes of 85 Kb and 135 Kb. FACS (loss of mTurquoise2 fluorescence), diagnostic PCR, karyotyping and sequencing revealed correct integration of all fragments (Suppl. Fig. S9, S10, S3).

**Figure 4:**
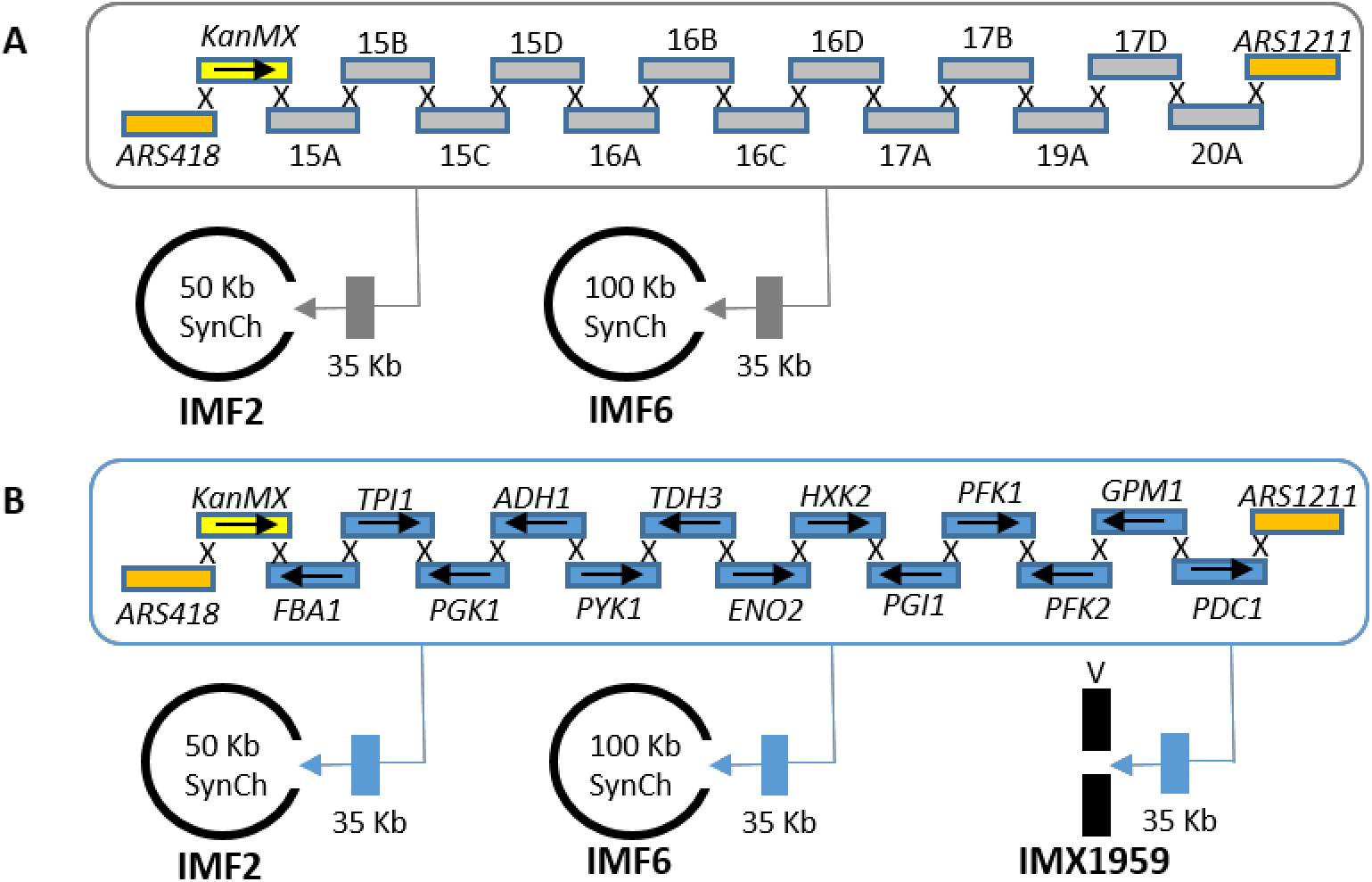
SynChs as landing pads. **A**) 35 Kb of non-coding DNA was simultaneously *in vivo* assembled and inserted by CRISPR/Cas9 at the *mTurquoise2* locus of the 100 Kb SynCh in IMF6 and of the 50 Kb SynCh in IMF2, resulting in 135 Kb (IMF11) and 85 Kb (IMF12) SynChs, respectively. **B**) 35 Kb of glycolytic genes were simultaneoulsy *in vivo* assembled and inserted using CRISPR/Cas9 at the *mTurquoise2* locus of the 100 Kb SynCh in IMF6 and of the 50 Kb SynCh of IMF2 resulting in 135 Kb (IMF13) and 85 Kb (IMF14) SynChs, respectively. As control, the same glycolytic cassettes were integrated at the native *CAN1* locus of chromosome V resulting in strain IMX1959.

Physiological characterization of the initial strains harbouring the four differently sized SynChs (50 Kb in strain IMF2, 85 Kb in strain IMF12, 100 Kb in strain IMF6 and 135 Kb in strain IMF11) revealed that they grew on average 8% slower than their prototrophic parental strain (IMX2059), irrespective of SynCh size (Fig. 5). Genome sequencing did not reveal mutations that could explain the decrease in growth rate (Suppl. Table S1-S2). Conversely, IMF23 (100 Kb Syn12.8), with the improved SynCh design, in which the highly expressed *mRuby2* was re-located from its place adjacent to the centromere and in which *Venus* (highly homologous to *mTurquoise2*) was removed, showed the same growth rate as the parental strain. As *S. cerevisiae* is tolerant to variations in the number of native chromosomes (27-29), the observed slower growth phenotype for four of the SynCh strains might be specific to the SynCh design. It has been previously shown that the size of artificial chromosomes affect their stability (30-33). In selective medium, SynCh loss would result in cells unable to grow and thereby in reduced growth rate. To evaluate the mitotic stability of the SynChs, IMF2, IMF12, IMF6, IMF11 and IMF23 were sequentially transferred in selective media and plated on both non-selective (YPD) and selective media (SMD). Along four successive culture transfers (ca. 25 generations), the viability of the strains on non-selective medium was similar to that of the prototrophic control strain CEN.PK113-7D (Fig. 5), or a control strain containing a 6.5 Kb centromeric plasmid (IMC153) (one-way ANOVA with Post-Hoc Tukey-Kramer, P>0.05).

**Figure 5:**
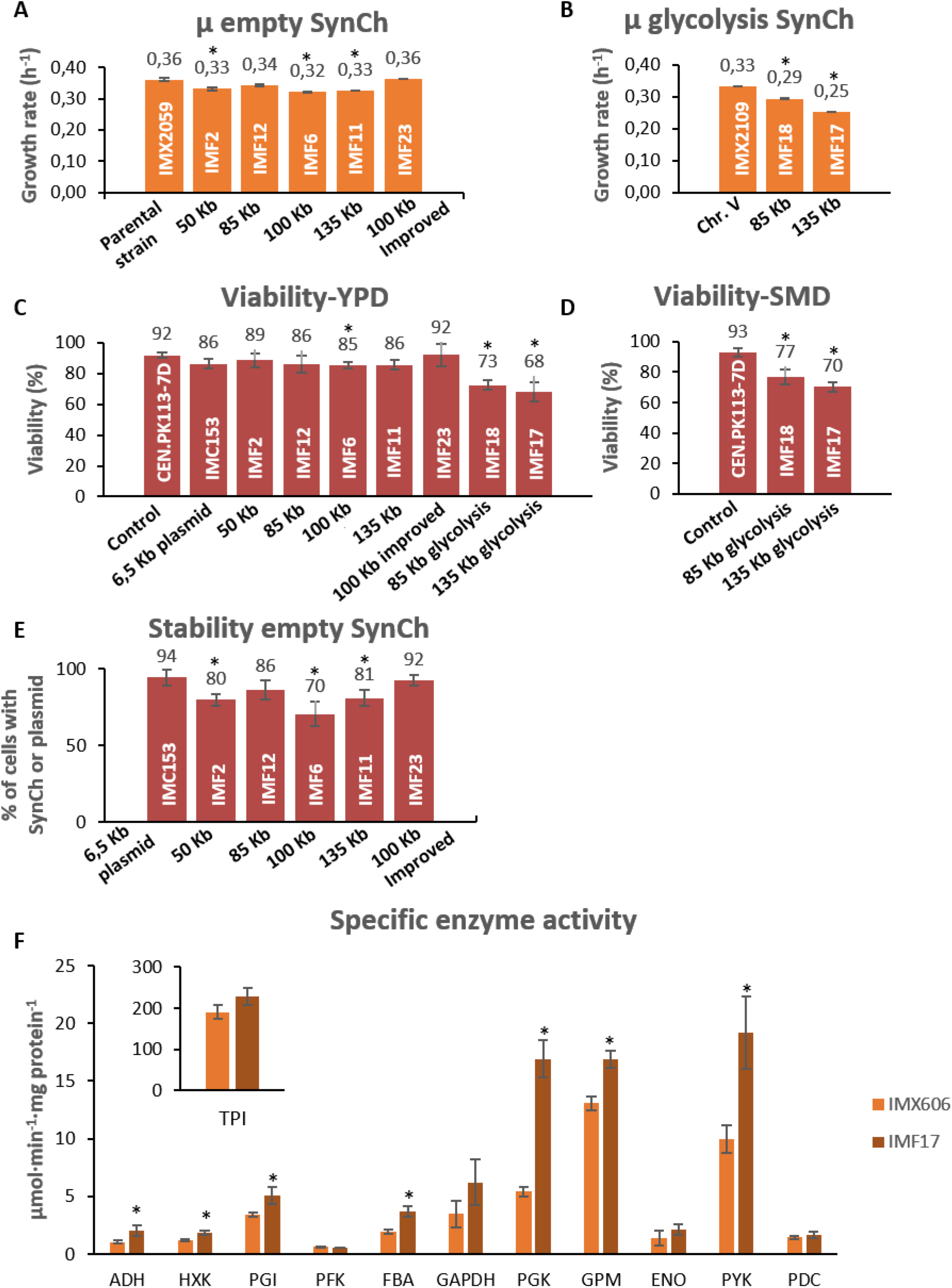
Physiological characterization of strains carrying synthetic chromosomes. **A** and **B**): Specific growth rate of strains carrying empty SynChs (Panel **A**: IMF2, IMF12, IMF6 and IMF11) and glycolytic SynChs (Panel **B**: IMF18 and IMF17). Growth rates represent the average and mean deviation of biological duplicates except for the parental strain which cultures were performed in biological quadruplicate. **C**) and **D**): viability measured as number of colonies on YPD (**C**) or SMD (**D**) sorted by FACS, divided by the total number of possible colonies (96), for the empty and glycolytic SynChs. Data represent biological duplicates and are averaged from 4 days of measurement for all strains except for the 100 Kb new design (2 samples at day 1 and day 4). **E**) Stability measured as the number of transformants on selective plates (SMD) divided by the number of colonies on non-selective plates (YPD). For each strain, the stability represents the average of 4 days of measurement in biological duplicates, except for 100 Kb new design (IMF23) for which 2 days of measurement were used (day 1 and day 4). **F**) Specific activity of glycolytic enzymes in IMF17 (135 Kb glycolysis SynCh) and a control strain expressing glycolysis from a native chromosome, IMX606 (16). Activities were measured at least in biological duplicates. For panels A, B, C, D and E all significant differences with respect to the first bar are indicated with an asterisk (one-way ANOVA with Post-Hoc Tukey-Kramer, P<0.05). For panel F, the asterisk indicates whether enzyme activities of IMF17 are significantly different with respect to IMX606 (16) (two-tailed paired homoscedastic t-test p<0.05).

Plating on selective medium revealed that for the four SynCh with the initial design (IMF2, IMF12, IMF6, IMF11), 14% to 30% of the population lost their SynCh (Fig. 5). This percentage remained stable over 25 generations (Suppl. Fig. S11), and most likely explained the reduced growth rate measured for SynCh-bearing strains. Indeed the SynCh with the improved design (IMF23), which did not show a reduced growth phenotype was highly stable with 92% of the population retaining the SynCh, which is the same as the control strain IMC153, carrying a 6.5 Kb centromeric plasmid with the same auxotrophic markers as located on the SynChs, and with the same parental strain as the SynCh strains (Fig. 5). Contrary to earlier reports (30,33) no correlation was observed between SynCh size and stability (IMF2, IMF12, IMF6 and IMF11 with increasing SynCh size showed stability of 80%, 86%, 70% and 81%, respectively). Therefore, empty SynChs can be very stable. Reduced stability observed for four of the SynCh is unlikely to have resulted from homology between *mTurquoise2* and *Venus*, since these were only both present in IMF2 and IMF6 and not in IMF12 and IMF11, which also showed reduced stability. The reduced stability was most likely caused by strong expression of *mRuby2* close to the centromere, which might cause chromosome mis-segregation or replication interference. This could result in a mother cell retaining one copy of the SynCh and the daughter cell zero copies (1:0 segregation) or the mother retaining two copies and the daughter zero (2:0 segregation) (Fig. 6C). From the coverage of reads belonging to the SynChs, we estimated that average copy numbers of 0.6, 1.5, 1.5, 1.7, for the 50 Kb (IMF2), 85Kb (IMF12), 100 Kb (IMF6) and 135 Kb (IMF11) SynChs, respectively (Fig. 6A and Suppl. Fig. S3). Since reads coverage was uniform along the SynChs, the most likely explanation for these non-integer average SynCh copy numbers far from one, is an heterogeneous population with cells carrying none, one, two, or more SynCh copies. Remarkably, the improved SynCh design (IMF23) showed an average copy number of one, confirming its stable design. Copy number variations were confirmed by monitoring mRuby2 fluorescence, which is present on all SynChs. For the SynChs with less optimal design, the larger the SynCh, the higher the fraction of cells with high fluorescence (indicating multiple copies, Fig. 6B). For the new design (IMF23) most cells showed the same fluorescence intensity as the control in which *mRuby2* (with the same promoter and terminator) was integrated in single copy in the genome of the SynCh parental strain (IMX2224) (Fig. 6B).

**Figure 6:**
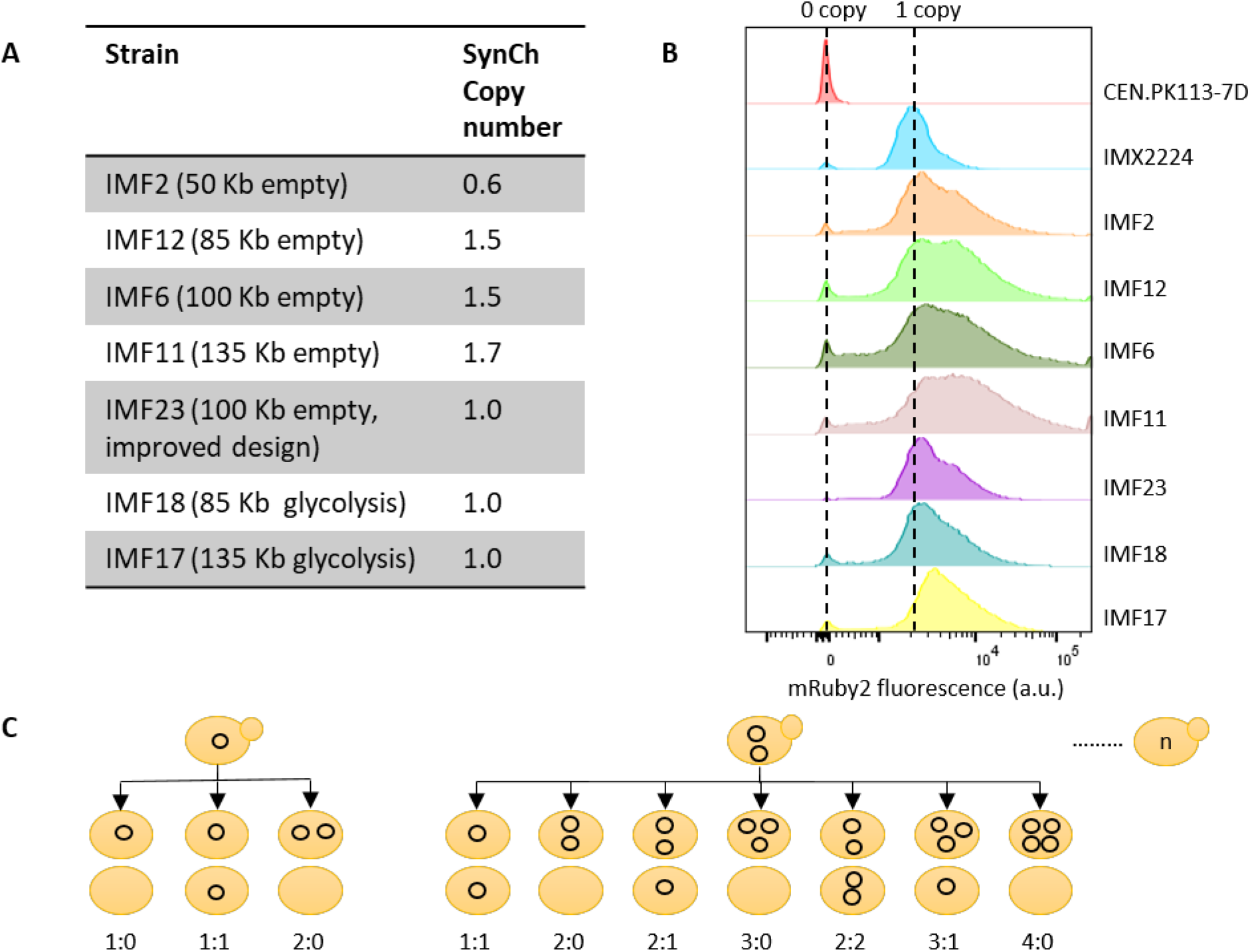
Quantification of SynCh copy number. **A**) SynCh copy number as determined by whole genome sequencing. **B**) SynCh copy number estimation based on mRuby2 fluorescence. CEN.PK113-7D with no copies of *mRuby2* was used as negative control and IMX2224 with a single copy of *mRuby2* integrated in the genome as positive control. **C**) Schematic overview of the potential segregation of circular SynCh upon cell division.

### Synthetic chromosomes as platforms for exclusive expression of (essential) pathways

SynChs have the potential to become powerful platforms for genome engineering, to rewire yeast native gene networks as well as to act as versatile expression hubs for heterologous pathways and processes. The transplantation of the complete human purine pathway to a neochromosome (34) is an encouraging demonstration, however the potential of SynChs as orthogonal expression platforms hasn’t been fully explored yet. Using the pathway swapping concept, we used SynChs as exclusive expression platform for the essential glycolytic pathway, and evaluated the impact of SynCh-borne glycolysis expression on yeast physiology. The 50 Kb (IMF2) and 100 Kb (IMF6) “empty” SynChs were assembled in the SwYG (16) ancestor strain IMX1338, in which the set of 13 genes coding for the glycolytic pathway has been relocalized to a single genomic locus. Using CRISPR-Cas9 editing combined with *in vivo* assembly of 13 glycoblocs (Fig.4), the 35 Kb glycolytic cassette was introduced in the *mTurquoise2* locus of the 50 Kb and 100 Kb SynChs. The successful removal of the single locus glycolysis from chromosome IX in these strains harbouring a second, SynCh-borne glycolytic gene set demonstrated that SynChs can serve as exclusive expression hubs for essential pathways (strain construction represented in Fig. 7). The strains carrying “glycolytic SynChs” IMF18 (glycolysis on 85 Kb SynCh) and IMF17 (glycolysis on 135 Kb SynCh) were grown in aerobic batch cultures in chemically defined medium with glucose as sole carbon source, and compared to IMX2109, a control strain carrying an identical genetic configuration of glycolysis on chromosome V. Both IMF17 and IMF18 grew significantly slower than the control strain (Fig. 5), IMF17 growing 14% slower than IMF18. Several hypotheses could explain this decreased growth rate of strains with glycolytic SynChs, i) the unfortunate occurrence of deleterious mutations in the native or synthetic chromosomes during transformation, ii) the low expression of SynCh-borne glycolytic genes and resulting decreased glycolytic flux, iii) the loss of SynChs, or iv) stalled replication due to transcription-replication clashes at the clustered glycolysis. Both IMF17 and IMF18 carried the same mutation in the *GPM1* promoter, in addition IMF17 carried non-synonymous mutations in *TPI1* and *PFK2* ORFs (Suppl. Table S2). However none of these mutations caused a decrease in the specific activity of the corresponding enzymes tested *in vitro* comparing IMF17 and the ancestral SwYG strain (16) (Fig 5), demonstrating that mutations in the glycolytic genes was most likely not responsible for the decreased growth rate of IMF17 and IMF18. A few mutations were observed in coding regions of other genes, and it cannot be excluded that they contributed to decreasing the growth rate of IMF17 and IMF18 (Suppl. Table S1). Secondly, the similar or higher specific activity in glycolytic enzymes between IMF17 and the ancestral SwYG strain(16) expressing glycolysis from a native chromosome (Fig. 5) indicated that low expression of SynCh-borne glycolytic genes was not involved in the reduced growth rate of IMF17 (and IMF18). Furthermore, previous studies have shown that an increase in specific activity of glycolytic enzymes does not result in a decrease in growth rate (35,36). Alternatively, this decreased growth rate might result from SynCh instability and potential loss in a fraction of the yeast population. Because loss of glycolytic SynChs leads to lethality in selective as well as in non-selective medium (e.g. YPD or YP ethanol), testing their stability is challenging. However, the viability of glycolytic SynCh strains on YPD could be used as indicator of SynCh loss when comparing with the viability on YPD of control strains and empty SynCh strains IMF12 and IMF11. IMF12 and IMF11 can survive SynCh loss on YPD medium, in which they display a viability similar to that of the control strain (85-92%). Conversely, the viability of IMF18 (85 Kb glycolytic SynCh) and IMF17 (135 Kb glycolytic SynCh), were significantly reduced on YPD to 73% and 68% respectively and on SMD medium to 77% and 70% respectively (Fig. 5). Assuming that this decreased viability results from SynCh loss, ca. 20% of the IMF17 and IMF18 population would lose their glycolytic SynCh. The SynCh loss would therefore be similar for SynChs with and without the glycolytic genes. The reduced growth rate of IMF17 and IMF18 does therefore most probably not result from a lower stability of glycolytic SynChs as compared to empty SynChs. Lastly, stalled replication of the SynChs due to transcription-replication clashes cannot be excluded as cause of the decreased specific growth rate.

**Figure 7:**
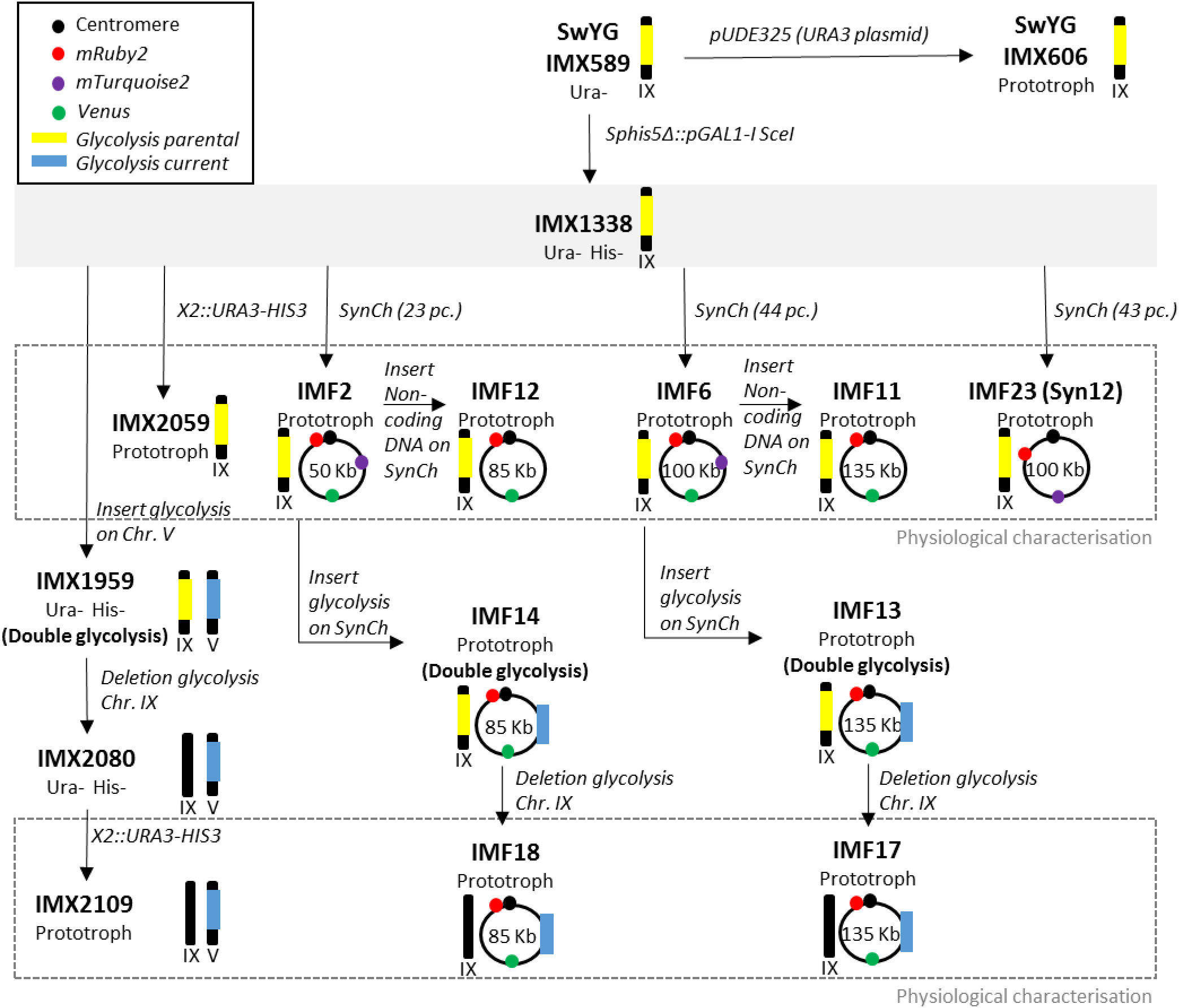
Strain construction overview. The parental strain IMX1338 is framed by a filled grey box from which its direct decedents are indicated with black arrows. The schematic representations of linear or circular chromosomes indicate the relevant native or synthetic chromosomes that were modified in this study, and the significance of the symbols is explained in the top left box. Strains characterized in this study are framed by a dashed grey box.

## DISCUSSION

The present study explores the potential of *de novo* assembled synthetic chromosomes to operate as large, modular expression islands in the popular workhorse and model strain *S. cerevisiae*. The design and construction workflow of SynChs developed in this study are highly versatile, enabling the facile construction of chromosomes with a wide variety of configurations.

Probing the limits of *in vivo* assembly, the present results demonstrate that a large number of DNA fragments of the size of expression cassettes can be efficiently assembled *de novo* in yeast. While other studies have reported this remarkable feature of *S. cerevisiae* (e.g. assembly of 38 fragments of 200 bp (13), and 25 fragments of 24 Kb (14)), quantitative information on the transformation and assembly efficiencies and fidelity was so far lacking. The SynCh assembly from 44 fragments, the record to date, showed a remarkably high efficiency, with 36% of faithfully assembled SynChs. This efficiency is substantially higher than previously reported, for instance by Annaluru et al. (37) where assembly of 27 fragments into chromosome III resulted in 0.5% of clones with the expected marker combination. Still, increasing the number of fragments was accompanied by a decrease in transformation efficiency and in the total number of clones on plates, which can become a limitation for the modular assembly of large SynChs. The construction of SynChs involves several cellular mechanisms, many of which are yet poorly understood but need to be considered to further enhance the SynChs construction efficiency. For instance, access of all required DNA fragments to the nucleus is essential for SynCh assembly, but the mechanisms involved in DNA translocation during transformation are not fully resolved and are poorly investigated. Once all DNA fragments have reached the nucleus, the HDR machinery has to be mobilized to repair a large number of lethal double strand DNA breaks (DSBs). While the number of simultaneous DNA breaks that *S. cerevisiae* tolerates is yet undefined, hundreds of simultaneous repairs most probably present a formidable challenge. Additionally, while HR is the preferred mechanism for DSB repair in *S. cerevisiae*, other, non-homologous repair mechanisms exist (38,39), and might take over when the HR machinery is inactive or saturated, thereby promoting erroneous assemblies. Inactivation of non-homologous end joining (NHEJ) is unfortunately not an option in *S. cerevisiae* as, due to the pleiotropic role of the NHEJ proteins, it leads to deleterious consequences (40-42). Last but not least, assembly in a complete SynCh is not a guaranty for cell survival, as they have to recover from the transformation stress, which is yet another poorly explored phenomenon. The combination of complex and often poorly understood events that have to take place for DNA to be assembled largely explains why typically a small fraction only of the transformed cell harbour the expected constructs (ca. one cell out of millions). One thing is however certain: the full potential of yeast *in vivo* assembly hasn’t been reached yet. While intensive efforts have been made to turn *E. coli* into a universal molecular biology tool, the power of *S. cerevisiae* as DNA assembly platform has so far been largely untapped. Despite the pivotal role played by *S. cerevisiae* in the last decade in the assembly of genomes (14,43-46) there is so far relatively little effort invested in turning this yeast into a universal and powerful DNA assembly platform (47,48), a situation that we expect to see change in the future.

SynCh stability and impact on the host physiology is a critical feature for the applicability of SynChs for synthetic biology. In the present study, introduction of four differently sized empty SynChs caused a decrease in specific growth rate and in SynCh loss in 20% of the cell population. In-depth analysis by Illumina sequencing and flow cytometric analysis showed a lower average SynCh copy number at population level and per cell for the smallest, 50 Kb chromosome. For larger SynChs the average copy number was above one, considering that part of the population did not carry a SynCh, it means that a substantial fraction of the cells carried more than one SynCh. This phenotype is most likely explained by aberrant segregation of the SynChs caused by strong expression of the gene located next to the centromere (49), impairing the centromere function. However the lower average copy number of the smallest SynCh could also be caused by failed replication or loss of the SynCh. Still, the average percentage of cells in a population (70 to 86%) containing at least one SynCh copy was similar between chromosomes of 50 to 135 Kb. Comparing the stability of the empty SynChs with published data is not straightforward as earlier work with YACs typically evaluates plasmid loss in non-selective media. However the absence of size-dependency of SynCh stability contrasts with earlier work showing that circular chromosomes increase in stability up to ∼100 Kb but decrease in stability above this size, a phenomenon tentatively attributed to the formation of dicentric dimeric circles (30,33,50). Preventing intensive transcriptional activity close to the centromere, and thereby decreasing mis-segregation events, resulted in restored growth rate and increased stability up to 92% of cells retaining the SynCh with an average copy number of one. Considering future applications in synthetic biology, population homogeneity might be further enhanced by linear SynCh design, either by linearization of the circular SynChs (for instance with the telomerator carried by the SynChs (23)) or direct, *de novo* assembly of linear SynChs. Alternatively, other strategies to improve segregation such as equipping chromosomes with synthetic kinetochores can be considered (51).

The present study demonstrates for the first time that synthetic chromosomes, fully *de novo* assembled from transcription-unit sized DNA parts can serve as exclusive, orthogonal expression platforms for essential metabolic pathways. There are few reports using synthetic and artificial chromosomes to express genes in yeast using heterologous genes or, in the case of the tRNA neochromosome, expressing a second, complete set of tRNAs, which understandably lead to a decrease in growth rate (34,50,52-55). In the present study, the native glycolytic and fermentative pathways were transplanted from a native chromosome to a SynCh with an identical genetic configuration (Fig. 4), a unique experimental set-up that enables the direct comparison of native and SynCh-borne expression of an entire pathway. This approach revealed that synch-based expression of an essential pathway reduced the host specific growth rate by ca. 14-24% and suggested a negative correlation between SynCh size and growth rate. Although this decrease in growth rate could be partly explained by aberrant segregation due to strong gene expression adjacent to the centromere as observed for the empty SynCh, other factors most probably play a role. Reduced expression and capacity of the glycolytic and fermentative enzymes or population heterogeneity in term of SynCh copy number are most likely not involved. One important aspect to consider is that the expression of the clustered glycolytic and fermentative genes is driven by some of the most highly transcriptionally active promoters, resulting in a transcriptional hotspot. An alternative explanation might therefore be a clash between the transcription and replication machineries, causing delayed replication (56). In IMX2109, carrying the same glycolytic cassette but on a native chromosome, such a conflict might be attenuated by the native genetic context (e.g. different chromatin structure, ARS sequences). Future SynCh architecture should therefore take into account directionality of transcription with respect to neighbouring ARSs and localization of highly transcribed genes.

Altogether this study highlights the potential of synthetic chromosomes to serve as platforms for modular assembly of native and heterologous pathways and paves the way towards modular genomes for synthetic biology.

## DATA AVAILABILITY

All processed data are available in the manuscript or as supplementary data. Upon request, raw data that support the findings of this paper will be made available. All genomic data for this paper have been deposited in the NCBI database (https://www.ncbi.nlm.nih.gov/) under the BioProjectID PRJNA596648.

## SUPPLEMENTARY DATA

Supplementary Data are available at xxx.

## AUTHOR CONTRIBUTION

EP, SD, JMD and PD-L designed the experiments and wrote the manuscript. EP, SD, LvB and STP performed the experiments. MvdB performed the bioinformatics analysis.

## ACKNOWLEDGEMENTS

We thank Marijke Luttik for assistance with assaying the glycolytic enzymes activity, Mark Bisschops for technical assistance with FACS analysis, Melanie Wijsman for assistance with the SynCh stability assays and Pilar de la Torre for sequencing of strains. We thank Anna Wronska and Thomas Perli for sharing the plasmids pGGkd017 and pUDR514, respectively. This project was funded by the AdLibYeast European Research Council (ERC) consolidator 648141 grant awarded to Pascale Daran-Lapujade.

## REFERENCES

1. Walker, G.M. and Walker, R.S.K. (2018) Chapter Three - Enhancing Yeast Alcoholic Fermentations. In Gadd, G. M. and Sariaslani, S. (eds.), Adv. Appl. Microbiol. Academic Press, Vol. 105, pp. 87–129.

2. Meehl, M.A. and Stadheim, T.A. (2014) Biopharmaceutical discovery and production in yeast. Curr. Opin. Biotechnol., 30, 120–127.

3. Chubukov, V., Mukhopadhyay, A., Petzold, C.J., Keasling, J.D. and Martín, H.G. (2016) Synthetic and systems biology for microbial production of commodity chemicals. Npj Syst. Biol. Appl., 2, 16009.

4. Li, Y., Li, S., Thodey, K., Trenchard, I., Cravens, A. and Smolke, C.D. (2018) Complete biosynthesis of noscapine and halogenated alkaloids in yeast. Proc. Natl. Acad. Sci. USA, 115, E3922–E3931.

5. Paddon, C.J. and Keasling, J.D. (2014) Semi-synthetic artemisinin: a model for the use of synthetic biology in pharmaceutical development. Nature Rev. Microbiol., 12, 355–367.

6. Hutchison, C.A., Chuang, R.-Y., Noskov, V.N., Assad-Garcia, N., Deerinck, T.J., Ellisman, M.H., Gill, J., Kannan, K., Karas, B.J., Ma, L. et al. (2016) Design and synthesis of a minimal bacterial genome. Science, 351, aad6253.

7. Veneziano, R., Shepherd, T.R., Ratanalert, S., Bellou, L., Tao, C. and Bathe, M. (2018) In vitro synthesis of gene-length single-stranded DNA. Sci. Rep., 8, 6548.

8. Hughes, R.A. and Ellington, A.D. (2017) Synthetic DNA Synthesis and Assembly: Putting the Synthetic in Synthetic Biology. Cold Spring Harb. Perspect. Biol., 9, a023812.

9. Kosuri, S. and Church, G.M. (2014) Large-scale de novo DNA synthesis: technologies and applications. Nat. Methods, 11, 499–507.

10. Casini, A., Storch, M., Baldwin, G.S. and Ellis, T. (2015) Bricks and blueprints: methods and standards for DNA assembly. Nat. Rev. Mol. Cell Biol., 16, 568–576.

11. Kok, S.d., Stanton, L.H., Slaby, T., Durot, M., Holmes, V.F., Patel, K.G., Platt, D., Shapland, E.B., Serber, Z., Dean, J. et al. (2014) Rapid and Reliable DNA Assembly via Ligase Cycling Reaction. ACS Synth. Biol., 3, 97–106.

12. Gibson, D.G., Benders, G.A., Andrews-Pfannkoch, C., Denisova, E.A., Baden-Tillson, H., Zaveri, J., Stockwell, T.B., Brownley, A., Thomas, D.W., Algire, M.A. et al. (2008) Complete Chemical Synthesis, Assembly, and Cloning of a *Mycoplasma genitalium* Genome. Science, 319, 1215–1220.

13. Gibson, D.G. (2009) Synthesis of DNA fragments in yeast by one-step assembly of overlapping oligonucleotides. Nucleic Acids Res., 37, 6984–6990.

14. Gibson, D.G., Benders, G.A., Axelrod, K.C., Zaveri, J., Algire, M.A., Moodie, M., Montague, M.G., Venter, J.C., Smith, H.O. and Hutchison, C.A. (2008) One-step assembly in yeast of 25 overlapping DNA fragments to form a complete synthetic *Mycoplasma genitalium* genome. Proc. Natl. Acad. Sci., 105, 20404–20409.

15. Bruschi, C., Gjuracic, K. and Tosato, V. (2006) Yeast Artificial Chromosomes. Encyclopedia of Life Sciences. John Wiley & Sons Ltd, Chichester.

16. Kuijpers, N.G., Solis-Escalante, D., Luttik, M.A., Bisschops, M.M., Boonekamp, F.J., van den Broek, M., Pronk, J.T., Daran, J.-M. and Daran-Lapujade, P. (2016) Pathway swapping: Toward modular engineering of essential cellular processes. Proc. Natl. Acad. Sci., 15060–15065.

17. Solis-Escalante, D., Kuijpers, N.G., Barrajon-Simancas, N., van den Broek, M., Pronk, J.T., Daran, J.-M. and Daran-Lapujade, P. (2015) A minimal set of glycolytic genes reveals strong redundancies in *Saccharomyces cerevisiae* central metabolism. Eukaryot. Cell, 14, 804–816.

18. Richardson, S.M., Mitchell, L.A., Stracquadanio, G., Yang, K., Dymond, J.S., DiCarlo, J.E., Lee, D., Huang, C.L.V., Chandrasegaran, S., Cai, Y. et al. (2017) Design of a synthetic yeast genome. Science, 355, 1040–1044.

19. Rideau, F., Le Roy, C., Descamps, E.C.T., Renaudin, H., Lartigue, C. and Bébéar, C. (2017) Cloning, Stability, and Modification of *Mycoplasma hominis* Genome in Yeast. ACS Synth. Biol., 6, 891–901.

20. Kuijpers, N.G., Solis-Escalante, D., Bosman, L., van den Broek, M., Pronk, J.T., Daran, J.-M. and Daran-Lapujade, P. (2013) A versatile, efficient strategy for assembly of multi-fragment expression vectors in *Saccharomyces cerevisiae* using 60 bp synthetic recombination sequences. Microb. Cell Fact., 12, 47.

21. Dhar, M.K., Sehgal, S. and Kaul, S. (2012) Structure, replication efficiency and fragility of yeast ARS elements. Res. Microbiol., 163, 243–253.

22. Sikorski, R.S. and Hieter, P. (1989) A system of shuttle vectors and yeast host strains designed for efficient manipulation of DNA in *Saccharomyces cerevisiae*. Genetics, 122, 19–27.

23. Mitchell, L.A. and Boeke, J.D. (2014) Circular permutation of a synthetic eukaryotic chromosome with the telomerator. Proc. Natl. Acad. Sci., 111, 17003–17010.

24. Lambert, T.J. (2019) FPbase: a community-editable fluorescent protein database. Nat. Methods, 16, 277–278.

25. Nagai, T., Ibata, K., Park, E.S., Kubota, M., Mikoshiba, K. and Miyawaki, A. (2002) A variant of yellow fluorescent protein with fast and efficient maturation for cell-biological applications. Nat. Biotechnol., 20, 87.

26. Goedhart, J., von Stetten, D., Noirclerc-Savoye, M., Lelimousin, M., Joosen, L., Hink, M.A., van Weeren, L., Gadella Jr, T.W.J. and Royant, A. (2012) Structure-guided evolution of cyan fluorescent proteins towards a quantum yield of 93%. Nat. Commun., 3, 751.

27. Shao, Y., Lu, N., Wu, Z., Cai, C., Wang, S., Zhang, L.-L., Zhou, F., Xiao, S., Liu, L., Zeng, X. et al. (2018) Creating a functional single-chromosome yeast. Nature, 560, 331–335.

28. Luo, J., Sun, X., Cormack, B.P. and Boeke, J.D. (2018) Karyotype engineering by chromosome fusion leads to reproductive isolation in yeast. Nature, 560, 392–396.

29. Gorter de Vries, A.R., Pronk, J.T. and Daran, J.G. (2017) Industrial Relevance of Chromosomal Copy Number Variation in *Saccharomyces* Yeasts. Appl. Environ. Microbiol., 83.

30. Murray, A.W., Schultes, N.P. and Szostak, J.W. (1986) Chromosome length controls mitotic chromosome segregation in yeast. Cell, 45, 529–536.

31. Surosky, R.T., Newlon, C.S. and Tye, B.-K. (1986) The mitotic stability of deletion derivatives of chromosome III in yeast. Proc. Natl. Acad. Sci., 83, 414–418.

32. Sleister, H.M., Mills, K.A., Blackwell, S.E., Killary, A.M., Murray, J.C. and Malone, R.E. (1992) Construction of a human chromosome 4 YAC pool and analysis of artificial chromosome stability. Nucleic Acids Res., 20, 3419–3425.

33. Hieter, P., Mann, C., Snyder, M. and Davis, R.W. (1985) Mitotic stability of yeast chromosomes: a colony color assay that measures nondisjunction and chromosome loss. Cell, 40, 381–392.

34. Agmon, N., Temple, J., Tang, Z., Schraink, T., Baron, M., Chen, J., Mita, P., Martin, J.A., Tu, B.P., Yanai, I. et al. (2019) Phylogenetic debugging of a complete human biosynthetic pathway transplanted into yeast. Nucleic Acids Res.

35. Hauf, J., Zimmermann, F.K. and Muller, S. (2000) Simultaneous genomic overexpression of seven glycolytic enzymes in the yeast *Saccharomyces cerevisiae*. Enzyme Microb. Technol., 26, 688–698.

36. Schaaff, I., Heinisch, J. and Zimmermann, F.K. (1989) Overproduction of glycolytic enzymes in yeast. Yeast, 5, 285–290.

37. Annaluru, N., Muller, H., Mitchell, L.A., Ramalingam, S., Stracquadanio, G., Richardson, S.M., Dymond, J.S., Kuang, Z., Scheifele, L.Z., Cooper, E.M. et al. (2014) Total Synthesis of a Functional Designer Eukaryotic Chromosome. Science, 344, 55–58.

38. Aylon, Y. and Kupiec, M. (2004) DSB repair: the yeast paradigm. DNA Repair (Amst), 3, 797–815.

39. Heyer, W.D., Ehmsen, K.T. and Liu, J. (2010) Regulation of homologous recombination in eukaryotes. Annu. Rev. Genet., 44, 113–139.

40. Barnes, G. and Rio, D. (1997) DNA double-strand-break sensitivity, DNA replication, and cell cycle arrest phenotypes of Ku-deficient *Saccharomyces cerevisiae*. Proc. Natl. Acad. Sci., 94, 867–872.

41. Laroche, T., Martin, S.G., Gotta, M., Gorham, H.C., Pryde, F.E., Louis, E.J. and Gasser, S.M. (1998) Mutation of yeast Ku genes disrupts the subnuclear organization of telomeres. Curr. Biol., 8, 653–656.

42. Gravel, S., Larrivee, M., Labrecque, P. and Wellinger, R.J. (1998) Yeast Ku as a regulator of chromosomal DNA end structure. Science, 280, 741–744.

43. Benders, G.A., Noskov, V.N., Denisova, E.A., Lartigue, C., Gibson, D.G., Assad-Garcia, N., Chuang, R.-Y., Carrera, W., Moodie, M., Algire, M.A. et al. (2010) Cloning whole bacterial genomes in yeast. Nucleic Acids Res., 38, 2558–2569.

44. Tagwerker, C., Dupont, C.L., Karas, B.J., Ma, L., Chuang, R.-Y., Benders, G.A., Ramon, A., Novotny, M., Montague, M.G., Venepally, P. et al. (2012) Sequence analysis of a complete 1.66 Mb *Prochlorococcus marinus* MED4 genome cloned in yeast. Nucleic Acids Res., 40, 10375–10383.

45. Karas, B.J., Molparia, B., Jablanovic, J., Hermann, W.J., Lin, Y.C., Dupont, C.L., Tagwerker, C., Yonemoto, I.T., Noskov, V.N., Chuang, R.Y. et al. (2013) Assembly of eukaryotic algal chromosomes in yeast. J. Biol. Eng., 7, 30.

46. Venetz, J.E., Del Medico, L., Wölfle, A., Schächle, P., Bucher, Y., Appert, D., Tschan, F., Flores-Tinoco, C.E., van Kooten, M., Guennoun, R. et al. (2019) Chemical synthesis rewriting of a bacterial genome to achieve design flexibility and biological functionality. Proc. Natl. Acad. Sci., 116, 8070–8079.

47. DiCarlo, J.E., Conley, A.J., Penttila, M., Jantti, J., Wang, H.H. and Church, G.M. (2013) Yeast oligo-mediated genome engineering (YOGE). ACS Synth. Biol., 2, 741–749.

48. Yu, S.C., Kuemmel, F., Skoufou-Papoutsaki, M.N. and Spanu, P.D. (2019) Yeast transformation efficiency is enhanced by TORC1- and eisosome-dependent signaling. MicrobiologyOpen, 8, e00730.

49. Hill, A. and Bloom, K. (1987) Genetic manipulation of centromere function. Mol. Cell. Biol., 7, 2397–2405.

50. Walker, R.S.K. (2017), The university of Edinburgh.

51. Lacefield, S., Lau, D.T. and Murray, A.W. (2009) Recruiting a microtubule-binding complex to DNA directs chromosome segregation in budding yeast. Nat. Cell Biol., 11, 1116–1120.

52. Naesby, M., Nielsen, S.V., Nielsen, C.A., Green, T., Tange, T.O., Simon, E., Knechtle, P., Hansson, A., Schwab, M.S., Titiz, O. et al. (2009) Yeast artificial chromosomes employed for random assembly of biosynthetic pathways and production of diverse compounds in *Saccharomyces cerevisiae*. Microb. Cell Fact., 8, 45.

53. Klein, J., Heal, J.R., Hamilton, W.D., Boussemghoune, T., Tange, T.O., Delegrange, F., Jaeschke, G., Hatsch, A. and Heim, J. (2014) Yeast synthetic biology platform generates novel chemical structures as scaffolds for drug discovery. ACS Synth. Biol., 3, 314–323.

54. Essani, K., Glieder, A. and Geier, M. (2015) Combinatorial pathway assembly in yeast. Vol. 2, pp. 423–436.

55. Hughes, S.R., Cox, E.J., Bang, S.S., Pinkelman, R.J., López-Núñez, J.C., Saha, B.C., Qureshi, N., Gibbons, W.R., Fry, M.R. and Moser, B.R. (2015) Process for assembly and transformation into *Saccharomyces cerevisiae* of a synthetic yeast artificial chromosome containing a multigene cassette to express enzymes that enhance xylose utilization designed for an automated platform. J. Lab. Autom., 20, 621–635.

56. Hamperl, S. and Cimprich, K.A. (2016) Conflict Resolution in the Genome: How Transcription and Replication Make It Work. Cell, 167, 1455–1467.

