## Supplementary material for "A supernumerary designer chromosome for modular *in vivo* pathway assembly in *Saccharomyces cerevisiae*"

### SUPPLEMENTARY DATA

#### Strains, maintenance and growth media

All *S. cerevisiae* strains used in this study (Suppl. Table S3-S4) were derived from the CEN.PK family (1). For non-selective growth and propagation, the yeast strains were grown on Yeast extract-Peptone (YP) medium containing: 10 g $L^{-1}$ Bacto yeast extract and 20 g $L^{-1}$ Bacto peptone. For selective growth, Synthetic Medium (SM) was used, consisting of: 3 g $L^{-1}$ $KH_2PO_4$, 0.5 g $L^{-1}$ $MgSO_4 \cdot 7H_2O$, 5 g $L^{-1}$ $(NH_4)_2SO_4$ and 1 mL $L^{-1}$ of a trace element solution (2). Both YP and SM medium were set to pH 6.0 using 2 M KOH and sterilised by autoclaving at 121°C for 20 min. Subsequently, synthetic medium was supplemented with 1 ml $L^{-1}$ of a filter sterilised vitamin solution. As a carbon source 20 g $L^{-1}$ glucose, sterilised at 110°C for 20 min, was supplemented to SM or YP medium, resulting in SMD or YPD medium, respectively. When auxothrophic strains were cultivated in SMD, it was supplemented with 125 mg $L^{-1}$ histidine and/or 150 mg $L^{-1}$ uracil. Screening of strains lacking the *URA3* gene was performed on SMD with 150 mg $L^{-1}$ uracil and 1 g $L^{-1}$ 5-FluoroOrotic Acid (SMD–5-FOA). For the selection based on the dominant markers *hphNT1* and *KanMX,* 200 mg $L^{-1}$ hygromycin (Hyg) and 200 mg $L^{-1}$ G418 were added to the medium, respectively. When G418 was used for selection in SMD medium, the 5 g $L^{-1}$ $(NH_4)_2SO_4$ in the SM was replaced with either 1.0 g $L^{-1}$ of L-glutamate (monosodium: $C_5H_8NO_4Na$; $H_2O$ content of 1 mol $mol^{-1}$) or 2.3 g $L^{-1}$ urea. Moreover 6.6 g $L^{-1}$ $K_2SO_4$ was added to compensate for sulfate supply. Solid medium was obtained by adding 20 g $L^{-1}$ Bacto agar to the YP or SM medium before autoclaving. Liquid yeast culture were grown in 500 mL shake flask with 100 mL medium at 30°C and 200 rpm in an Innova incubator (New Brunswick Scientific, Edison, NJ), unless stated otherwise.

*Escherichia coli* XL1-blue was used for propagation and isolation of plasmids. *E.coli* was grown in 5 mL Lysogeny Broth (10 g $L^{-1}$ Bacto tryptone, 5 g $L^{-1}$ Bacto yeast extract and 5 g $L^{-1}$ NaCl) supplemented with 100 mg $L^{-1}$ ampicillin when required. Cultivation was performed in 5 mL Greiner tubes or 25 mL shake flasks at 37°C and 200 rpm in an Innova 4000 shaker (New Brunswick Scientific).

For storage of *S. cerevisiae* and *E.coli* strains, the cultures were mixed with glycerol (30% v/v) and stored in 1 mL vials at -80°C.

#### Molecular biology techniques

Genomic DNA from *S. cerevisiae* used for amplification of integrative cassettes or fragments to build synthetic chromosomes, was isolated with the YeaStar genomic DNA kit (Zymo Research, Irvine, CA) or the QIAGEN Blood & Cell Culture Kit with 100/G Genomic-tips (Qiagen, Hilden, Germany) following the manufacturer's recommendations. Genomic DNA of *E.coli*, to serve as template for fragments of the synthetic chromosomes, was isolated by pelleting 1.5 mL of overnight grown culture and resuspension of the cell pellet in 600 µl of Lysis buffer (10 mM Tris-HCl (pH 8.0), 1 mM EDTA, 0.6% SDS (v/v), 0.12 g $L^{-1}$ proteinase K). This mixture was incubated for 1 hour at 37°C, whereafter the protocol I of the YeaStar genomic DNA kit (Zymo research) was followed from step 4 onwards (600 µl instead of 250 µl of chloroform was added). Plasmids were isolated from *E.coli* using the Sigma GenElute Plasmid DNA miniprep kit (Sigma-Aldrich, St. Louis, MO).

All DNA fragments for transformation in *S. cerevisiae* or *E.coli* of less than 10 Kb were amplified by PCR using Phusion High-Fidelity DNA Polymerase (Thermo Fisher Scientific, Waltham, MA), all fragments of 10 Kb or longer were amplified with LongRange PCR (Qiagen) according to the manufacturer's instructions. *E.coli* DNA was amplified with desalted primers while coding DNA from *S. cerevisiae* or plasmids (Suppl. Table S19-S20) was amplified with PAGE-purified oligonucleotides (Suppl. Tables S5-S18) (Sigma-Aldrich). PCR products were purified by either one of three different methods. First, by separation using electrophoresis on a 1% (w/v) agarose gel (Thermo Fisher Scientific) in 1X TAE buffer (Thermo Fisher Scientific) with subsequent purification with the Zymoclean Gel DNA Recovery kit (Zymo Research). Second, by the GenElute PCR Clean-Up kit (Sigma-Aldrich). Third by size selection using the AMPure XP beads (Beckman Coulter, Brea, CA) according to the supplier's protocol. Purity of fragments was checked using the NanoDrop 2000 spectrophotometer (Thermo Fisher Scientific) and concentration was measured either by the NanoDrop 2000 (Thermo Fisher Scientific) or by the Qubit dsDNA BR Assay kit (Thermo Fisher Scientific) using the Qubit 2.0 Fluorometer (Invitrogen, Carlsbad, CA).

Transformation in *S. cerevisiae* was executed using the lithium acetate/polyethylene glycol method (3). To verify correct transformants by diagnostic PCR, genomic DNA was isolated using an SDS/lithium acetate protocol (4) or using the YeaStar genomic DNA kit (Zymo Research). Unless stated otherwise, single-colony isolates were obtained by three consecutive re-streaks on selective solid medium and verification of the accurate genotype by diagnostic PCR before storage at -80°C. Plasmids were transformed in *E.coli* XL1-blue using chemical transformation (5) and verified by restriction analysis or diagnostic PCR after isolation. All diagnostic PCRs were performed using DreamTaq PCR Master Mix (Thermo Fisher Scientific) according to the manufacturers instruction.

#### Plasmid construction

Plasmids used in this study are described in Suppl. Table S19. Plasmids carrying the fluorescent markers *mRuby2* (pUDC191), *mTurquoise2* (pUDC192) and *Venus* (pUDC193) were constructed by Golden Gate *Bsa*I assembly using the plasmid parts provided in the Yeast Toolkit (6). First the GFP dropout plasmid pUD538 was built using Golden Gate *BsaI* cloning, which consisted of ConLS (pYTK002), *sfGFP* (pYTK047), ConR1 (pYTK067), AmpR (pYTK083), *URA3* (pYTK074) and *CEN6/ARS4* (pYTK081). pUDC191 was assembled from the GFP dropout plasmid pUD538, *pCCW12* (pYTK010), *mRuby2* (pYTK034) and *tENO1* (pYTK051). pUDC192 was assembled from pUD538, *pTEF2* (pYTK014), *mTurquoise2* (pYTK032) and *tSSA1* (pYTK052). Finally, pUDC193 was assembled from pUD538, *pTEF1* (pYTK013), *Venus* (pYTK033) and *tTDH1* (pYTK056). A golden gate reaction mixture was made by using 20 fmol of each part plasmid or dropout plasmid, 1 µL T4 DNA ligase buffer (Thermo Fisher Scientific, Waltham, MA), 0.5 µL T7 DNA ligase (NEB New England Biolabs, Ipswich, MA), 0.5 µL FastDigest *Eco31*I (*Bsa*I) (Thermo Fisher Scientific) and filled to a final volume of 10 µL with MilliQ $H_2O$. The reaction was performed in a thermocycler with 25 cycles of digestion at 42°C for 2 min followed by ligation at 16°C for 5 min. The reaction was ended by a final digestion at 60°C for 10 min followed by heat inactivation of the enzyme at 80°C for 10min. 1 µl of the assembly reaction was transformed in chemical competent *E.coli* XL1-blue cells as described in the supplementary materials. Correct transformants were screened based on the presence/absence of the GFP gene, which could be identified by colony colour on the Safe Imager 2.0 Blue-Light Transilluminator (Thermo Fisher Scientific). The transformants were subsequently also confirmed by PCR.

The transcriptional units of the major glycolytic and fermentation enzymes *FBA1*, *TPI1*, *PGK1*, *ADH1*, *PYK1*, *TDH3*, *ENO2*, *HXK2*, *PGI1*, *PFK1*, *PFK2*, *GPM1* and *PDC1* were also cloned in plasmids by Golden Gate assembly to facilitate amplification by PCR (plasmids listed in Suppl. Table S19). The promoter (800bp)-gene-terminator (300bp) sequences were amplified from the genome of CEN.PK113-7D using primers containing *Bsa*I sites (Suppl. Table S5). Using *Bsa*I Golden Gate assembly as described earlier, the entry plasmid pGGkd017 was constructed from pYTK002, pYTK047, pYTK072, pYTK074, pYTK082 and pYTK083 and used as backbone for assembly of the glycolysis plasmids.

pUDR400, a guide RNA (gRNA) plasmid for introducing Cas9-mediated double strand breaks in *mTurquoise2*, was constructed as described by Mans *et al.(7)* with pROS12 as backbone and gRNA-containing primers listed in Suppl. Table S6. Similarly gRNA plasmids pUDR514, pUDR547 and pUDR557 were constructed targeting the YPRcTau3 locus (primer 'YPRCtau3_targetRNA FW'), the intergenic *X2* locus (primer 'FW_X-2_gRNA') and the *sga1* SinLoG (primer 'CRISPR RNA Recycl fw') locus, respectively (Suppl. Table S6).

#### Strain construction

The strains constructed in this study are derived from SwYG described in Kuipers *et al.* (8) (Fig. 7). SwYG is characterized by the genetic reduction and relocalization to the *SGA1* locus of the set of genes encoding the glycolytic and fermentative pathways. As host strain for synthetic chromosome assembly, IMX1338 was constructed by integration in SwYG's genome of an inducible expression cassette for the meganuclease I-SceI. To this end, an I-SceI expression cassette consisting of *pGAL1-I-SceI-tCYC1* was amplified from pUDC073. Two fragments to assemble a gRNA plasmid for a double strand break (DSB) in *SpHIS5* were amplified as described by Mans *et al.(7)*. The expression cassette and two gRNA fragments were transformed in strain IMX589 and correct colonies were confirmed by diagnostic PCR. The assembled *URA3*-based gRNA plasmid was removed by counter selection on plates containing 5-FOA before the strain was stocked.

For the construction of strains carrying *in vivo* assembled synthetic chromosomes (IMF1, IMF2, IMF6), the amount of DNA fragments transformed was kept constant for all experiments with 200 fmol of each *E.coli* filler fragment, 200 fmol of each fluorescent marker and 100 fmol of the *CEN6/ARS4*, *ARS* and selectable markers. The total amount of transformed DNA therefore depended on the number and nature of DNA parts in the chromosome design (Suppl. Table S20-S22). DNA fragments for assembly of synthetic chromosomes were amplified from genomic DNA of *E.coli* XL1 blue/*E.coli* BL21, genomic DNA of *S. cerevisiae* CEN.PK113-7D or plasmids as specified in Suppl. Table S20. All primers for amplification of synthetic chromosome fragments consisted of a primer binding part of 18 bp to 25 bp, flanked by 60 bp Synthetic Homologous Recombination sequences (SHR) (9). IMF1, IMF2 (50 Kb) and IMF6 (100 Kb) were assembled from ~2.5 Kb *E.coli* filler fragments, three fluorescent markers: *mRuby2*, *mTurquoise2*, *Venus*, Autonomously Replicating Sequences (ARS), *CEN6/ARS4*, a Telomerator fragment and a *HIS3* selective marker (Fig. 2, Suppl. Table S21-S22). After amplification and purification of these fragments they were concentrated by chromatography with Vivacon 500 PCR grade columns (Sartorius, Gottingen, Germany), according to the manufacturer's instruction. The fragments for each SynCh were pooled using the amounts described above. This was supplemented with MilliQ $H_2O$ to a final volume of 50 µl, before transformation. The transformants were checked by fluorescent microscopy, FACS, CHEF and Whole Genome Sequencing (WGS) as described below.

The synthetic chromosomes were used as landing pads for *in vivo* assembly and integration of two types of DNA constructs, both 35 Kb long. Both constructs were flanked by *ARS* sequences and carried the *KanMX* selectable marker, but differed by the presence of non-coding *E.coli* filler fragments for one, and 13 functional glycolytic and fermentative genes from yeast for the other (Suppl. Table S23-S26). Integration of these constructs in the 50 Kb SynCh of IMF2 and 100 Kb SynCh of IMF6 was facilitated by Cas9-mediated editing of the SynChs at the *mTurquoise2* locus. Integration of the *E. coli* and glycolytic constructs in IMF6 resulted in strains IMF11 and IMF13, respectively; and in IMF2 resulted in IMF12 and IMF14, respectively. As control, the same 35 Kb glycolytic construct was also integrated in IMX1338 genomic DNA at the *CAN1* locus using Cas9-mediated editing (strain IMX1959).

In more detail, for construction if IMF12 (85 Kb) and IMF11 (135 Kb), *E.coli* fragments adding up to ~35 Kb were inserted at the *mTurquoise2* locus of IMF2 (50 Kb) and IMF6 (100 Kb). To this end, IMF2 and IMF6 were transformed with pUDR400 for introduction of a DSB at *mTurquoise2* and with sixteen DNA fragments flanked with 60 bp SHRs for *in vivo* assembly and insertion at that locus (Fig. 4 and Suppl. Table S25-S26). The gRNA plasmid was removed by non-selective growth and correct transformants were verified by PCR, CHEF and WGS. For construction of IMF13 (135 Kb) and IMF14 (85 Kb), the major paralogs of glycolysis flanked by *ARS* sequences and the *KanMX* selective marker, were inserted at the *mTurquoise2* locus of IMF2 (50 Kb) and IMF6 (100 Kb). The glycolytic expression cassettes consisted of 800bp promoters, the gene and 300 bp terminators. The 16 DNA fragments (Fig. 4 and Suppl. Table S23-S24), forming a total of ~35 Kb, were *in vivo* assembled and inserted at the *mTurquoise2* locus, facilitated by the DSB induced by pUDR400. For construction of IMX1959, the same DNA fragments were also transformed in IMX1338 for insertion of the same configuration of the single locus glycolysis in the *can1* locus. To induce the DSB, the gRNA plasmid p426-SNR52p-gRNA.CAN1.Y-SUP4t was used. Again, gRNA plasmids were removed from all strains by non-selective growth and correct transformants were verified by PCR, CHEF and WGS.

Strains IMF13, IMF14 and IMX1959 contained two genetic copies of the glycolytic and fermentative pathways. The copy harboured in the *SGA1* locus on chromosome IX was removed to obtain strains with a single locus glycolysis, either carried by a SynCh (IMF17 and IMF18) or by chromosome V (CAN1 locus, strain IMX2080). Removal of glycolysis from IMF13, IMF14 and IMX1959 resulted in strains IMF17, IMF18 and IMX2080 respectively. More precisely, strains IMF13, IMF14 and IMX1959 were transformed with pUDR557 and a repair fragment (synthesised by annealing the primers COUNTER SELECT oligo fw and COUNTER SELECT oligo rv (Suppl. Table S15)). Removal of the SinLoG at *sga1* was confirmed by PCR and gRNA plasmids were removed by non-selective growth. Finally strain IMX2080 was made prototrophic by inserting an expression cassette for *URA3* and *HIS3* in the *X2* locus. Two fragments were amplified $X2_{flank}$*-pURA3-URA3-tURA3-SHR DT* from the *S. cerevisiae* CEN.PK113-7D genome and *SHR DT-pHIS3-HIS3-tHIS3-*$X2_{flank}$ from pLM092 (Suppl. Table S16). These fragments were transformed together with gRNA plasmid pUDR547 in IMX2080. After validation of correct insertion at the *X2* locus by diagnostic PCR the gRNA plasmid was removed by non-selective growth and the strain was stocked as IMX2109. The original IMX1338 strain (single locus glycolysis on chromosome IX) was made prototrophic in the same way, resulting in strain IMX2059. The strains IMF17, IMF18 and IMX2109 were confirmed by WGS. Strain IMX2059 was confirmed by diagnostic PCR.

Fluorescent control strains IMX2224 (*mRuby2)*, IMX2225 (*Venus*) and IMX2226 (*mTurquoise2)* were constructed, with stable expression from the genome, with the same promoters and terminators as used for expression from the SynCh and in the same genetic background as for the SynCh strains. *pCCW12-mRuby2-tENO1* was amplified with primers YPRcTau3_pCCW12_fw and YPRcTau3_tENO1_rv from pUDC191, *pTEF1-Venus-tTDH1* was amplified with primers YPRcTau3_pTEF1_fwd and YPRcTau3_tTDH1_rev from pUDC193 and *pTEF2-mTurquoise2-tSSA1* was amplified with primers

YPRcTau3_pTEF2_fw and YPRcTau3_tSSA1_rv from pUDC193 (Suppl. Table S17). The strain were made by transforming the individual fragments together with pUDR514 in IMX2059. gRNA plasmids were removed from all strains by non-selective growth and correct transformants were verified by PCR and FACS.

The prototrophic stability control strain IMC153 was made by transforming a centromeric 6.5 Kb plasmid (pLM092) in IMX1338. Fluorescent control strains for mRuby2 (IMC111), mTurquoise2 (IMC112) and Venus (IMC113) were made by transforming centromeric plasmids with these markers in IMX581 (CEN.PK113-5D with Cas9).

All strains used in this study are listed in Suppl. Table S3-S4 and Figure 7 gives a graphic overview of the most important strains in this study.

#### Experimental quantification of SynCh assembly efficiency

To compare transformation efficiency of differently sized DNA fragments, 2.5 Kb, 5 Kb and 10 Kb fragments were transformed in IMX1338 to build SynChs. The SynCh design was the same as 100 Kb SynCh of IMF6 and the 50 Kb SynCh of IMF2, except the telomerator fragment was omitted since it had only 30 bp homology in comparison to the 60 bp of the other fragments. Therefore colonies were selected based on *HIS3* prototrophy alone instead of the *URA3* and *HIS3* prototrophy of IMF2 and IMF6. 2.5 Kb *E.coli* filler fragments were amplified from *E.coli* XL1 blue/*E.coli* BL21 genome. 5 Kb and 10 Kb fragments were amplified from genomic DNA isolated from strain IMF1 (Suppl. Fig. S12). The fragments were purified and concentrated before transformation. The fragments outlined in Suppl. Tables S27-S32 were transformed in the concentrations described for the construction of IMF1, IMF2 and IMF6 into IMX1338. Both a 50 Kb chromosome as well as a 100 Kb chromosome were assembled from all three fragment sizes. In the transformation the IMX1338 competent cells were also transformed with 50 µl of MilliQ $H_2O$ and several dilutions (1.0 x $10^5$ and 1.0 x $10^6$) were plated on YPD. For the transformation with the different size fragments also several dilutions of the transformation mix were plated (100x, 20x and 10x), which were considered as technical replicates when single colonies could be counted. Based on the colony forming units (CFU) counted on the plates, and the viability of the cells based on the YPD control, the efficiency (CFU/$10^8$ cells) was calculated. The experiment was performed in biological duplicate, by performing the same transformations on different days.

Syn12, a 100 Kb SynCh with a different design, contained two fluorescent markers: *mRuby2*, *mTurquoise2* (to prevent homology between *mTurquoise2* and *Venus*), Autonomously Replicating Sequences (ARS), *CEN6/ARS4*, a Telomerator fragment and a *HIS3* selective marker (Suppl. Table S33 and Suppl. Fig. S6). After one restreak, the total number of correct transformants were assessed based on screening by FACS, CHEF and WGS (Suppl. Table S34). Four correct transformants were stocked as IMF23 to IMF26.

#### Fluorescence detection by microscopy and flow cytometry

To validate the expression of the fluorescent proteins mRuby2, Venus and mTurquoise2, from their genes on the SynCh, the transformants were checked by fluorescent microscopy and/or by flow cytometry.

For validation based on fluorescent microscopy, single colony isolates were cultivated overnight in SMD. Subsequently, 5 µl of culture were spotted on a 18x18 mm coverslip and placed on a microscopic slide. Fluorescence of the cells was observed by the ZEISS Axio Imager Z1 microscope (Carl Zeiss AG, Oberkochen, Germany) , with three different filter sets. *mRuby2* was detected by Filter set 14 with excitation at 535 nm and emission at 590 nm. *Venus* was visualized using filter set 9, with excitation at 470 nm and emission at 515 nm. Finally filter set 47 was used to detect *mTurquoise2* at an excitation of 436 nm and an emission of 480 nm.

The BDFACSAria™ II Cell Sorter in combination with the BDFACSDiva software (BD Biosciences, Franklin Lakes, NJ) was used for flow cytometry analysis. The machine was equipped with 355 nm, 445 nm, 488 nm, 561 nm and 640 nm lasers and a 70 µm nozzle, and operated with filtered FACSFlow™ (BD Biosciences). Prior to every experiment, a CST cycle with the corresponding CS&T Beads (BD Biosciences) was run to evaluate a correct cytometer performance. Furthermore if the FACS was used for sorting an Auto Drop Delay cycle with the corresponding Accudrop Beads (BD Biosciences) was run to determine the drop delay. mTurquoise2 was excited by the 445 nm laser and the emission was detected using a 525 nm bandpass filter with a bandwidth of 50 nm. Venus was excited by the 488 nm laser and the emission was detected using a 545 nm bandpass filter with a bandwidth of 30 nm. mRuby2 was excited by the 561 nm laser and the emission was detected using a 582 nm bandpass filter with a bandwidth of 15 nm.

For FACS analysis, single colony isolates were grown overnight in SMD. In the morning the culture was transferred to new SMD at an $OD_{660}$ of 0.5. The culture was grown for 2 generations to an $OD_{660}$ of ~2.0. The level of fluorescence was analysed by plotting the according fluorescence versus the area of the cell of 10,000 or 100,000 events. The sorting regions (‘gates’) of fluorescence or no fluorescence were based on the reference strain CEN.PK113-7D taken along in the experiments. FACS data were analysed using FlowJo® software (version 3.05230, FlowJo, LLC, Ashland, OR, USA).

#### Contour-clamped homogeneous electric field (CHEF) electrophoresis

Agarose plugs containing genomic DNA were made using the CHEF Yeast Genomic DNA Plug Kit (Bio-Rad laboratories, Hercules, CA) following the manufacturer's instructions. For in-plug linearization of the synthetic chromosomes, an I-SceI digestion (Thermo Fischer Scientific) was performed in the plugs. For this, the plugs were washed for 1 hour in 1 mL 0.1x wash buffer (Bio-Rad laboratories) per plug. Subsequently, the wash buffer was decanted and fresh wash buffer was added to sufficiently cover the plugs. The wash buffer was aspirated and the plugs were then equilibrated for 1 hour in 1 mL of 1x Tango buffer without Mg-acetate per plug. The buffer was aspirated again and 100 µL fresh Tango buffer without Mg-acetate and 2 µL of I-SceI enzyme were added to each plug. This was incubated for 2 hours on ice, after which 10 µL of 100 mM Mg-acetate was added to each plug. After 1h or overnight incubation, the buffer was removed and the plugs were kept in 1 mL 1x wash buffer each. One-half of each plug was used per well and 1 gradation (on the syringe) thin slice of the Lambda PFG Ladder (New England Biolabs) was used as standard in a 1% Certified™ Megabase Agarose (Bio-Rad laboratories) gel in 0.5x TBE Electrophoresis Buffer (prepared from a 10x TBE Electrophoresis Buffer stock; Thermo Fischer Scientific).

The chromosomes were separated using a Bio-Rad Electrophoresis Cell in combination with a CHEF-DR® II Control Module and a CHEF-DR® II Drive module (Bio-Rad laboratories). The CHEF electrophoresis was run for 24 hours with the following parameters: an initial and final switch time of 35 seconds, a pulse angle of 120°, a voltage of 5.3 V/cm, a flow of 70 (about 0.75 L/min), a set temperature of 14°C and a buffer volume of 1.5 L. The CHEF gel was stained for 15 minutes with ethidium bromide (3 µg $mL^{-1}$) in 0.5x TBE buffer and unstained for 15 minutes with 0.5x TBE buffer. The gel was visualised with an InGenius LHR gel Imaging System (Syngene, Bangalore, India).

#### Sequencing

For whole genome sequencing, genomic DNA was isolated using the QIAGEN Blood & Cell Culture Kit with 100/G Genomic-tips (Qiagen), according to the manufacturer's instructions. The Miseq Reagent Kit v3 (Illumina, San Diego, CA, USA), was used to obtain 300 bp reads for paired-end sequencing. DNA was sheared to 550 bp with the M220 ultrasonicator (Covaris, Wolburn, MA) and subsequently the TruSeq DNA PCR-Free Library Preparation kit (Illumina) was employed for a six strain library. The samples were quantified by qPCR on a Rotor-Gene Q PCR cycler (Qiagen) using the KAPA Library Quantification Kit (Kapa Biosystems, Wilmington, MA). Finally the library was sequenced using an illumina MiSeq sequencer (Illumina, San Diego, CA). Data were mapped to the CEN.PK113-7D (10) genome with corresponding *in silico* SynCh design using the Burrows-Wheeler alignment tool (11) (version 0.7.15). Data was visualised using the Integrated Genomics Viewer (IGV) (12) (version 2.4.0). Chromosomal copy number variations (CNV) were quantified with the Magnolya algorithm (13) (version 0.15). Finally an overview of single nucleotide polymorphisms (SNPs) were made using the SAMtools (11) (version 1.3.1) and Pilon (14) (with -vcf setting; version 1.18). Sequenced strains are indicated in Suppl. Table S3.

Sanger sequencing (Baseclear, Leiden, The Netherlands) was performed on the original purified *E.coli* chunks used for assembly of the SynCh (primers in Suppl. Table S18).

#### Physiological characterization

A 1 mL aliquot stored at -80°C was used to inoculate a wake-up culture of 100 mL SMD in a 500 mL shake flask. The culture was grown until late exponential phase and used to inoculate a pre-culture. The pre-culture was grown until mid-exponential phase and used to inoculate a measuring culture in biological duplicate at an initial $OD_{660}$ of 0.3. Optical densities at 660 nm of the cultures were monitored by a Jenway 7200 spectrophotometer in technical duplicate (Cole-Parmer, Vernon Hills, IL). A maximum specific growth rate ($\mu_{max}$) was calculated from at least 5 data points in the exponential phase.

#### SynCh stability determination

SynCh stability was assessed based on plating on selective and non-selective growth medium with respect to the SynCh borne *URA3* and *HIS3* markers. Strains were inoculated from a freezer stock in 20 mL of selective SMD medium in a 100 mL shake flask and grown overnight. The strains were then inoculated in biological duplicate to fresh 20 mL medium in a 100 mL shake flask at an $OD_{660}$ of 0.5 and were grown for 2 generations until an approximate $OD_{660}$ of 2.0. From the culture 3 mL were taken, washed in sterile COULTER® ISOTON® II Diluent (Beckman Coulter, Brea, CA) and subsequently resuspended in 3 mL sterile COULTER® ISOTON® II Diluent (Beckman Coulter) to be sorted by the FACS. 96 cells of the entire population were sorted on both solid non-selective YPD and solid selective SMD medium in a single well OmniTray (Thermo Fischer Scientific).

The viability was determined by:

$$Viability\ (\%) = \frac{number\ of\ colonies\ grown\ on\ YPD\ plate}{96} \times 100\%$$

The stability was determined by:

$$\text{Stability (\%)} = \frac{number\ of\ colonies\ grown\ on\ SMG\ plate}{number\ of\ colonies\ on\ YPD\ plate} \times 100\%$$

After the samples for the FACS were taken the cultures were placed back in the incubator and grown until the morning, after which the biological duplicates of each strain were transferred to new medium at an $OD_{660}$ of 0.5 and the abovementioned protocol was repeated. In total, the strains were analysed by FACS for 4 consecutive days, except for IMF23 which was analysed on the first and the fourth day.

#### *In vitro* enzyme activities

For determining the glycolytic enzyme activities, strain IMF17 was inoculated from a freezer stock in 100 mL selective SMD medium in a 500 mL shake flask. It was transferred in biological duplicate to fresh 100 mL SMD medium and grown until mid-exponential phase at an $OD_{660}$ of 8. 50 mL Of cell culture was harvested from each biological duplicate by centrifuging for 10 minutes at 5000 rpm at 4˚C. Cell extracts were prepared as described by Postma *et al.*(15). Assays were prepared in at least technical duplicates for HXK, PGI, FBA, TPI, GAPDH (16.5 U of enzyme instead of 25U), PGK, GPM, ENO, PYK, PDC and ADH as described by Jansen *et al.* (16) and PFK according to Cruz *et al*. (17). Cell extracts were diluted with 100 mM $KH_2PO_4$ buffer with 2 mM $MgCl_2$ (pH 7.5) when necessary, except for the assay for TPI, for which the cell extract was diluted with demineralized water. The spectrophotometric assays were carried out in an U-3010 spectophotometer (Hitachi, Tokyo, Japan) at 30 °C and 340 nm. The protocol by Lowry *et al.* (18) was used to determine protein concentrations in the cell extract, using bovine serum albumin (fatty acid-free) as standard.

#### Statistical analysis

If replicates were performed, data are presented as mean ± s.d. and the number of replicates are indicated. Statistical differences were determined either by using a t-test (paired or unpaired is indicated) for comparison of two samples using excel or for comparison of multiple samples by ANOVA with Post-Hoc Tukey-Kramer using GraphPad Prism 4 (Graphpad, San Diego, CA). For both tests, differences were considered significant when $P<0.05$.

**Table S1 Amino acid substitutions in native genome identified in the constructed strains as compared to most relevant parental strain.**

| Systematic name | Name | Type | Amino acid change |
|---|---|---|---|
| **IMF2 compared to IMX1338** | | | |
| No mutations observed | | | |
| **IMF6 compared to IMX1338** | | | |
| YEL074W | - | Non-synonymous | Pro-34-His |
| **IMF24 (genome of Syn12.1) compared to IMX1338** | | | |
| YER116C | SLX8 | Non-synonymous | Pro-150-Leu |
| **IMF25 (genome Syn12.3) compared to IMX1338** | | | |
| No mutations observed | | | |
| **IMF26 (genome Syn12.4) compared to IMX1338** | | | |
| YCL052C | PBN1 | Non-synonymous | Ile-109-Thr |
| YOL076W | MDM20 | Non-synonymous | Leu-249-Ile |
| **IMF23 (genome Syn12.8) compared to IMX1338** | | | |
| No mutations observed | | | |
| **IMF12 compared to IMF2** | | | |
| YMR323W | ERR3 | Non-synonymous | Leu-447-Phe |
| **IMF11 compared to IMF6** | | | |
| YKL205W | LOS1 | Synonymous | Val-860-Val |
| **IMF18 compared to IMF2** | | | |
| YBR084W | MIS1 | Non-synonymous | Phe-547-Leu |
| YDL159W | STE7 | Non-synonymous | Cys-447-Ser |
| YDR295C | HDA2 | Non-synonymous | Ala-556-Glu |
| YGL150C | INO80 | Non-synonymous | Ala-882-Val |
| YBL113C | - | Intron | - |
| YPL006W | NCR1 | Synonymous | Gly-119-Gly |
| YPR147C | - | Synonymous | Leu-52-Leu |
| **IMF17 compared to IMF6** | | | |
| YFL025C | BST1 | Non-synonymous | Asn-791-His |
| YIL169C | CSS1 | Non-synonymous | Thr-906-Ser |
| YKL176C | LST4 | Non-synonymous | Ser-705-Tyr |
| **IMX2109 compared to IMX1338** | | | |
| YJR162C | - | Synonymous | Thr-33-Thr |
| YIL130W | ASG1 | Non-synonymous | Asp-617-Asn |
| YKR004C | ECM9 | Non-synonymous | Ser-47-Tyr |
| YPL283W-A | - | Intron | |
| YLR172C | DPH5 | Non-synonymous | Ala-79-Thr |
| YNR050C | LYS9 | Synonymous | Phe-150-Phe |

#### Table S2 Mutation identified in Synthetic Chromosomes and IMX2109 SinLoG as compared with *in silico* design and with the most relevant parental strain

For the SynChs containing *E.coli* DNA it was discovered that the template DNA used for PCR amplification and subsequent transformation was a mix of two *E.coli* strains: XL1-Blue and BL21. Therefore all mutations identified from the *in silico* SynCh design (based on *E.coli* BL21) which were identical to the sequence of *E.coli* XL1-Blue were discarded. To verify that indeed these variation was already present in PCR fragments used for transformation and did not occur during SynCh assembly/propagation, a part of the PCR fragments used for transformation were sanger sequenced (Suppl. Fig. S13). For IMF11, IMF12, IMF17 and IMF18, five SNPs were observed for ARS418 and 2 for ARS1211 with respect to the in silico design. However these were verified not to be real SNPs, since for the in silico template S228C was used while ARS418 and ARS1211 were amplified from CEN.PK113-7D. These strains indeed differ in the 7 SNPs observed for the two ARSs.

*The SNP in *TPI1* for IMF17 was a non-synonymous Val-7-Phe mutation.

#The SNP in *PFK2* for IMF17 was a non-synonymous Met-390-Ile mutation.

| Position | Fragment | Mutation type |
|---|---|---|
| **SynCh2 (IMF2)** | | |
| 12242 | *HIS3* promoter | C to CA |
| 18451 | SHR BD | C to CT |
| 24938 | *TDH1* terminator (*Venus*) | 34 bp deletion |
| 42085 | SHR BK | C to CA |
| 48959 | SHR AD | G to A |
| **SynCh1 (IMF6)** | | |
| 1703 | Chunk 1A | A to T |
| 22970 | Chunk 3A | C to T |
| 30653 | SHR AH | AC to A |
| 32769 | *HIS3* promoter | C to CA |
| 38952 | SHR BD | T to TG |
| 46433 | SHR BF | CA to C |
| 54835 | Chunk 5A | A to T |
| 56692 | SHR BG | T to TC |
| 59309 | SHR BH | AT to A |
| 59315 | SHR BH | AC to A |
| 65869 | *TDH1* terminator (*Venus*) | 34 bp deletion |
| 71313 | SHR BQ | C to CT |
| 77341 | Telomerator (*URA3* CDS) | G to GA (in stretch of A’s) |
| 77997 | SHR AO | C to CA |
| 87376 | Chunk 7D | G to A |
| 100069 | SHR AD | A to AG |
| **Syn12.1 (IMF24)** | | |
| 17843 | SHR BO | G to GC |
| 39753 | Chunk 15C | Insertion of 12 bp |
| 97403 | *SSA1* terminator (*mTurquoise2*) | CAT to C |
| **Syn12.3 (IMF25)** | | |
| 64232 | *HIS3* promoter | C to CA |
| **Syn12.4 (IMF26)** | | |
| 7596 | SHR BL | A to AT |
| 20696 | *CCW12* promoter (*mRuby2*) | CA to C |
| 22174 | *ARS1* | A to AC |
| 24751 | Chunk 18A | GGT to G |
| 37589 | SHR DH | C to T |
| 43110 | *CEN6/ARS4* | AT to A |
| 51015 | SHR DM | GA to G |
| 53542 | SHR DN | C to T |
| 64232 | *HIS3* promoter | C to CA |

| | | |
|---|---|---|
| 65169 | *HIS3* terminator | C to T |
| 70141 | Chunk 4B | C to T |
| 75441 | SHR AK | T to TC |
| 97403 | *SSA1* terminator (*mTurquoise2*) | CAT to C |
| **Syn12.8 (IMF23)** | | |
| 43 | SHR AO | AT to A |
| 20696 | *CCW12* promoter (*mRuby2*) | C to CA |
| 40152 | SHR DI | A to AC |
| 40171 | SHR DI | AG to A |
| 41708 | Chunk 15D | G to T |
| 56968 | Chunk 17B | C to T |
| 64232 | *HIS3* promoter | C to CA |
| 77907 | SHR BF | A to AG |
| 77911 | SHR BF | AC to A |
| 80507 | SHR BS | Deletion of 12 bp |
| 93363 | SHR BI | GA to G |
| 97595 | SHR DS | T to TC |
| **Syn20 (IMF12) as compared to SynCh2 (IMF2)** | | |
| 10248 | SHR AH | C to CG |
| 14530 | SHR DF | G to C |
| 23462 | Chunk 16A | G to A |
| 33079 | Chunk 17A | A to G |
| 40176 | Chunk 17D | TA to T |
| **Syn19 (IMF11) as compared to SynCh1 (IMF6)** | | |
| 40004 | SHR DI | GC to G |
| 62440 | Chunk 17D | C to T |
| 63119 | SHR DR | AG to A |
| **Syn16 (IMF18) as compared to SynCh2 (IMF2)** | | |
| 23389 | SHR 0 | CT to C |
| 30653 | SHR C | A to AG |
| 43179 | *GPM1* promoter | Deletion of 18 bp (AT's) |
| **Syn15 (IMF17) as compared to SynCh1 (IMF6)** | | |
| 32334 | SHR AA | AT to A |
| 35439 | *TPI1** | G to T |
| 41106 | SHR N | T to TG |
| 60046 | *PFK2* [#] | C to G |
| 62020 | SHR L | AT to A |
| 63582 | *GPM1* promoter | Deletion of 18 bp (AT's) |
| 63970 | SHR M | T to TG |
| 105411 | Chunk 6B | Insertion of 59 bp |
| 105412 | Chunk 6B | T to C |
| 105415 | Chunk 6B | C to T |
| 105417 | Chunk 6B | G to A |
| 105421 | Chunk 6B | T to C |
| 105430 | Chunk 6B | G to A |
| 8590 | *tADH1* | A to T |
| 20368 | *tHXK2* | AT to A |
| 30676 | *pPFK2* | GA to G |
| 32946 | *pGPM1* | Deletion of 18 bp (AT's) |

#### Table S3 *S. cerevisiae* strains used in this study

Strains that were whole genome sequence in this study are marked with a *. SHRs are differently annotated than in Kuijpers *et al.* (8). SHRs are annotated in subscript between de genetic fragments that they join together.

| Strain | Relevant genotype | Source |
|---|---|---|
| **CEN.PK113-7D** | *MATa URA3 HIS3 LEU2 TRP1 MAL2-8c SUC2* | Entian *et al.*(1) |
| **IMX581** | *MATa ura3-52 HIS3 LEU2 TRP1 MAL2-8c SUC2 can1 Δ::( pTEF1-Spcas9-tCYC1 natNT1)* | Mans *et al.*(7) |
| **IMX589** | *MATa ura3-52 his3-1 leu2-3,112 MAL2-8c SUC2 glk1Δ:: (pAgTEF1-SpHIS5-tAgTEF1) hxk1Δ::KlLEU2 tdh1Δ tdh2Δ gpm2Δ gpm3Δ eno1Δ pyk2Δ pdc5Δ pdc6Δ adh2Δ adh5Δ adh4Δ sga1Δ::(* $_{G}$ *tFBA1-FBA1-pFBA1* $_{H}$ *pTPI1-TPI1-tTPI1* $_{P}$ *tPGK1-PGK1-pPGK1* $_{Q}$ *tADH1-ADH1-pADH1* $_{N}$ *pPYK1-PYK1-tPYK1* $_{O}$ *tTDH3-TDH3-pTDH3* $_{A}$ *pENO2-ENO2-tENO2* $_{B}$ *pHXK2-HXK2-tHXK2* $_{C}$ *pPGI-PGI1-tPGI1* $_{D}$ *pPFK1-PFK1-tPFK1* $_{J}$ *tPFK2-PFK2-pPFK2* $_{K}$ *pAgTEF1-AmdSYM-tAgTEF1* $_{L}$ *tGPM1-GPM1-pPGM1* $_{M}$ *pPDC1-PDC1-tPDC1-SYN* $_{F}$*) pyk1Δ pgi1Δ tpi1Δ tdh3Δ pfk2Δ::(pTEF1-Spcas9-tCYC1 natNT1) pgk1Δ gpm1Δ fba1Δ hxk2Δ pfk1Δ adh1Δ pdc1Δ eno2Δ* | Kuijpers *et al.*(8) |
| **IMX1338*** | *MATa ura3-52 his3-1 leu2-3,112 MAL2-8c SUC2 glk1Δ::(pAgTEF1-SpHIS5-tAgTEF1)Δ::(pGAL1-I SceI-tCYC1) hxk1Δ::KlLEU2 tdh1Δ tdh2Δ gpm2Δ gpm3Δ eno1Δ pyk2Δ pdc5Δ pdc6Δ adh2Δ adh5Δ adh4Δ sga1Δ::(* $_{G}$ *tFBA1-FBA1-pFBA1* $_{H}$ *pTPI1-TPI1-tTPI1* $_{P}$ *tPGK1-PGK1-pPGK1* $_{Q}$ *tADH1-ADH1-pADH1* $_{N}$ *pPYK1-PYK1-tPYK1* $_{O}$ *tTDH3-TDH3-pTDH3* $_{A}$ *pENO2-ENO2-tENO2* $_{B}$ *pHXK2-HXK2-tHXK2* $_{C}$ *pPGI-PGI1-tPGI1* $_{D}$ *pPFK1-PFK1-tPFK1* $_{J}$ *tPFK2-PFK2-pPFK2* $_{K}$ *pAgTEF1-AmdSYM-tAgTEF1* $_{L}$ *tGPM1-GPM1-pPGM1* $_{M}$ *pPDC1-PDC1-tPDC1-SYN* $_{F}$*) pyk1Δ pgi1Δ tpi1Δ tdh3Δ pfk2Δ::(pTEF1-Spcas9-tCYC1 natNT1) pgk1Δ gpm1Δ fba1Δ hxk2Δ pfk1Δ adh1Δ pdc1Δ eno2Δ* | This study |
| **IMF1*** | *MATa ura3-52 his3-1 leu2-3,112 MAL2-8c SUC2 glk1Δ::(pAgTEF1-SpHIS5-tAgTEF1)Δ::(pGAL1-I SceI-tCYC1) hxk1Δ::KlLEU2 tdh1Δ tdh2Δ gpm2Δ gpm3Δ eno1Δ pyk2Δ pdc5Δ pdc6Δ adh2Δ adh5Δ adh4Δ sga1Δ::(* $_{G}$ *tFBA1-FBA1-pFBA1* $_{H}$ *pTPI1-TPI1-tTPI1* $_{P}$ *tPGK1-PGK1-pPGK1* $_{Q}$ *tADH1-ADH1-pADH1* $_{N}$ *pPYK1-PYK1-tPYK1* $_{O}$ *tTDH3-TDH3-pTDH3* $_{A}$ *pENO2-ENO2-tENO2* $_{B}$ *pHXK2-HXK2-tHXK2* $_{C}$ *pPGI-PGI1-tPGI1* $_{D}$ *pPFK1-PFK1-tPFK1* $_{J}$ *tPFK2-PFK2-pPFK2* $_{K}$ *pAgTEF1-AmdSYM-tAgTEF1* $_{L}$ *tGPM1-GPM1-pPGM1* $_{M}$ *pPDC1-PDC1-tPDC1-SYN* $_{F}$*) pyk1Δ pgi1Δ tpi1Δ tdh3Δ pfk2Δ::(pTEF1-Spcas9-tCYC1 natNT1) pgk1Δ gpm1Δ fba1Δ hxk2Δ pfk1Δ adh1Δ pdc1Δ eno2Δ SynCh1 (wrong assembly: duplication)* | This study |
| **IMF2*** | *MATa ura3-52 his3-1 leu2-3,112 MAL2-8c SUC2 glk1Δ::(pAgTEF1-SpHIS5-tAgTEF1)Δ::(pGAL1-I SceI-tCYC1) hxk1Δ::KlLEU2 tdh1Δ tdh2Δ gpm2Δ gpm3Δ eno1Δ pyk2Δ pdc5Δ pdc6Δ adh2Δ adh5Δ adh4Δ sga1Δ::(* $_{G}$ *tFBA1-FBA1-pFBA1* $_{H}$ *pTPI1-TPI1-tTPI1* $_{P}$ *tPGK1-PGK1-pPGK1* $_{Q}$ *tADH1-ADH1-pADH1* $_{N}$ *pPYK1-PYK1-tPYK1* $_{O}$ *tTDH3-TDH3-pTDH3* $_{A}$ *pENO2-ENO2-tENO2* $_{B}$ *pHXK2-HXK2-tHXK2* $_{C}$ *pPGI-PGI1-tPGI1* $_{D}$ *pPFK1-PFK1-tPFK1* $_{J}$ *tPFK2-PFK2-pPFK2* $_{K}$ *pAgTEF1-AmdSYM-tAgTEF1* $_{L}$ *tGPM1-GPM1-pPGM1* $_{M}$ *pPDC1-PDC1-tPDC1-SYN* $_{F}$*) pyk1Δ pgi1Δ tpi1Δ tdh3Δ pfk2Δ::(pTEF1-Spcas9-tCYC1 natNT1) pgk1Δ gpm1Δ fba1Δ hxk2Δ pfk1Δ adh1Δ pdc1Δ eno2Δ SynCh2* | This study |
| **IMF6*** | *MATa ura3-52 his3-1 leu2-3,112 MAL2-8c SUC2 glk1Δ::(pAgTEF1-SpHIS5-tAgTEF1)Δ::(pGAL1-I SceI-tCYC1) hxk1Δ::KlLEU2 tdh1Δ tdh2Δ gpm2Δ gpm3Δ eno1Δ pyk2Δ pdc5Δ pdc6Δ adh2Δ adh5Δ adh4Δ sga1Δ::(* $_{G}$ *tFBA1-FBA1-pFBA1* $_{H}$ *pTPI1-TPI1-tTPI1* $_{P}$ *tPGK1-PGK1-pPGK1* $_{Q}$ *tADH1-ADH1-pADH1* $_{N}$ *pPYK1-PYK1-tPYK1* $_{O}$ *tTDH3-TDH3-pTDH3* $_{A}$ *pENO2-ENO2-tENO2* $_{B}$ *pHXK2-HXK2-tHXK2* $_{C}$ *pPGI-PGI1-tPGI1* $_{D}$ *pPFK1-PFK1-tPFK1* $_{J}$ *tPFK2-PFK2-pPFK2* $_{K}$ *pAgTEF1-AmdSYM-tAgTEF1* $_{L}$ *tGPM1-GPM1-pPGM1* $_{M}$ *pPDC1-PDC1-tPDC1-SYN* $_{F}$*) pyk1Δ pgi1Δ tpi1Δ tdh3Δ pfk2Δ::(pTEF1-* | This study |

| | | |
|---|---|---|
| | Spcas9-tCYC1 natNT1) pgk1Δ gpm1Δ fba1Δ hxk2Δ pfk1Δ adh1Δ pdc1Δ eno2Δ SynCh1 | |
| **IMF11*** | MATa ura3-52 his3-1 leu2-3,112 MAL2-8c SUC2 glk1Δ::(pAgTEF1-SpHIS5-tAgTEF1)Δ::(pGAL1-I SceI-tCYC1) hxk1Δ::KlLEU2 tdh1Δ tdh2Δ gpm2Δ gpm3Δ eno1Δ pyk2Δ pdc5Δ pdc6Δ adh2Δ adh5Δ adh4Δ sga1Δ::( $_{G}$ tFBA1-FBA1-pFBA1 $_{H}$ pTPI1-TPI1-tTPI1 $_{P}$ tPGK1-PGK1-pPGK1 $_{Q}$ tADH1-ADH1-pADH1 $_{N}$ pPYK1-PYK1-tPYK1 $_{O}$ tTDH3-TDH3-pTDH3 $_{A}$ pENO2-ENO2-tENO2 $_{B}$ pHXK2-HXK2-tHXK2 $_{C}$ pPGI-PGI1-tPGI1 $_{D}$ pPFK1-PFK1-tPFK1 $_{J}$ tPFK2-PFK2-pPFK2 $_{K}$ pAgTEF1-AmdSYM-tAgTEF1 $_{L}$ tGPM1-GPM1-pPGM1 $_{M}$ pPDC1-PDC1-tPDC1-SYN $_{F}$) pyk1Δ pgi1Δ tpi1Δ tdh3Δ pfk2Δ::(pTEF1-Spcas9-tCYC1 natNT1) pgk1Δ gpm1Δ fba1Δ hxk2Δ pfk1Δ adh1Δ pdc1Δ eno2Δ Syn19 | This study |
| **IMF12*** | MATa ura3-52 his3-1 leu2-3,112 MAL2-8c SUC2 glk1Δ::(pAgTEF1-SpHIS5-tAgTEF1)Δ::(pGAL1-I SceI-tCYC1) hxk1Δ::KlLEU2 tdh1Δ tdh2Δ gpm2Δ gpm3Δ eno1Δ pyk2Δ pdc5Δ pdc6Δ adh2Δ adh5Δ adh4Δ sga1Δ::( $_{G}$ tFBA1-FBA1-pFBA1 $_{H}$ pTPI1-TPI1-tTPI1 $_{P}$ tPGK1-PGK1-pPGK1 $_{Q}$ tADH1-ADH1-pADH1 $_{N}$ pPYK1-PYK1-tPYK1 $_{O}$ tTDH3-TDH3-pTDH3 $_{A}$ pENO2-ENO2-tENO2 $_{B}$ pHXK2-HXK2-tHXK2 $_{C}$ pPGI-PGI1-tPGI1 $_{D}$ pPFK1-PFK1-tPFK1 $_{J}$ tPFK2-PFK2-pPFK2 $_{K}$ pAgTEF1-AmdSYM-tAgTEF1 $_{L}$ tGPM1-GPM1-pPGM1 $_{M}$ pPDC1-PDC1-tPDC1-SYN $_{F}$) pyk1Δ pgi1Δ tpi1Δ tdh3Δ pfk2Δ::(pTEF1-Spcas9-tCYC1 natNT1) pgk1Δ gpm1Δ fba1Δ hxk2Δ pfk1Δ adh1Δ pdc1Δ eno2Δ Syn20 | This study |
| **IMF13** | MATa ura3-52 his3-1 leu2-3,112 MAL2-8c SUC2 glk1Δ::(pAgTEF1-SpHIS5-tAgTEF1)Δ::(pGAL1-I SceI-tCYC1) hxk1Δ::KlLEU2 tdh1Δ tdh2Δ gpm2Δ gpm3Δ eno1Δ pyk2Δ pdc5Δ pdc6Δ adh2Δ adh5Δ adh4Δ sga1Δ::( $_{G}$ tFBA1-FBA1-pFBA1 $_{H}$ pTPI1-TPI1-tTPI1 $_{P}$ tPGK1-PGK1-pPGK1 $_{Q}$ tADH1-ADH1-pADH1 $_{N}$ pPYK1-PYK1-tPYK1 $_{O}$ tTDH3-TDH3-pTDH3 $_{A}$ pENO2-ENO2-tENO2 $_{B}$ pHXK2-HXK2-tHXK2 $_{C}$ pPGI-PGI1-tPGI1 $_{D}$ pPFK1-PFK1-tPFK1 $_{J}$ tPFK2-PFK2-pPFK2 $_{K}$ pAgTEF1-AmdSYM-tAgTEF1 $_{L}$ tGPM1-GPM1-pPGM1 $_{M}$ pPDC1-PDC1-tPDC1-SYN $_{F}$) pyk1Δ pgi1Δ tpi1Δ tdh3Δ pfk2Δ::(pTEF1-Spcas9-tCYC1 natNT1) pgk1Δ gpm1Δ fba1Δ hxk2Δ pfk1Δ adh1Δ pdc1Δ eno2Δ Syn13 | This study |
| **IMF14** | MATa ura3-52 his3-1 leu2-3,112 MAL2-8c SUC2 glk1Δ::(pAgTEF1-SpHIS5-tAgTEF1)Δ::(pGAL1-I SceI-tCYC1) hxk1Δ::KlLEU2 tdh1Δ tdh2Δ gpm2Δ gpm3Δ eno1Δ pyk2Δ pdc5Δ pdc6Δ adh2Δ adh5Δ adh4Δ sga1Δ::( $_{G}$ tFBA1-FBA1-pFBA1 $_{H}$ pTPI1-TPI1-tTPI1 $_{P}$ tPGK1-PGK1-pPGK1 $_{Q}$ tADH1-ADH1-pADH1 $_{N}$ pPYK1-PYK1-tPYK1 $_{O}$ tTDH3-TDH3-pTDH3 $_{A}$ pENO2-ENO2-tENO2 $_{B}$ pHXK2-HXK2-tHXK2 $_{C}$ pPGI-PGI1-tPGI1 $_{D}$ pPFK1-PFK1-tPFK1 $_{J}$ tPFK2-PFK2-pPFK2 $_{K}$ pAgTEF1-AmdSYM-tAgTEF1 $_{L}$ tGPM1-GPM1-pPGM1 $_{M}$ pPDC1-PDC1-tPDC1-SYN $_{F}$) pyk1Δ pgi1Δ tpi1Δ tdh3Δ pfk2Δ::(pTEF1-Spcas9-tCYC1 natNT1) pgk1Δ gpm1Δ fba1Δ hxk2Δ pfk1Δ adh1Δ pdc1Δ eno2Δ Syn14 | This study |
| **IMF17*** | MATa ura3-52 his3-1 leu2-3,112 MAL2-8c SUC2 glk1Δ::(pAgTEF1-SpHIS5-tAgTEF1)Δ::(pGAL1-I SceI-tCYC1) hxk1Δ::KlLEU2 tdh1Δ tdh2Δ gpm2Δ gpm3Δ eno1Δ pyk2Δ pdc5Δ pdc6Δ adh2Δ adh5Δ adh4Δ sga1Δ pyk1Δ pgi1Δ tpi1Δ tdh3Δ pfk2Δ::(pTEF1-Spcas9-tCYC1 natNT1) pgk1Δ gpm1Δ fba1Δ hxk2Δ pfk1Δ adh1Δ pdc1Δ eno2Δ Syn15 | This study |
| **IMF18*** | MATa ura3-52 his3-1 leu2-3,112 MAL2-8c SUC2 glk1Δ::(pAgTEF1-SpHIS5-tAgTEF1)Δ::(pGAL1-I SceI-tCYC1) hxk1Δ::KlLEU2 tdh1Δ tdh2Δ gpm2Δ gpm3Δ eno1Δ pyk2Δ pdc5Δ pdc6Δ adh2Δ adh5Δ adh4Δ sga1Δ pyk1Δ pgi1Δ tpi1Δ tdh3Δ pfk2Δ::(pTEF1-Spcas9-tCYC1 natNT1) pgk1Δ gpm1Δ fba1Δ hxk2Δ pfk1Δ adh1Δ pdc1Δ eno2Δ Syn18 | This study |
| **IMF23*, IMF24*, IMF25*, IMF26*** | MATa ura3-52 his3-1 leu2-3,112 MAL2-8c SUC2 glk1Δ::(pAgTEF1-SpHIS5-tAgTEF1)Δ::(pGAL1-I SceI-tCYC1) hxk1Δ::KlLEU2 tdh1Δ tdh2Δ gpm2Δ gpm3Δ eno1Δ pyk2Δ pdc5Δ pdc6Δ adh2Δ adh5Δ adh4Δ sga1Δ pyk1Δ pgi1Δ tpi1Δ tdh3Δ pfk2Δ::(pTEF1-Spcas9- | This study |

| | | |
|---|---|---|
| | *tCYC1 natNT1) pgk1Δ gpm1Δ fba1Δ hxk2Δ pfk1Δ adh1Δ pdc1Δ eno2Δ Syn12* | |
| **IMX1959** | *MATa ura3-52 his3-1 leu2-3,112 MAL2-8c SUC2 glk1Δ::(pAgTEF1-SpHIS5-tAgTEF1)Δ::(pGAL1-I SceI-tCYC1) hxk1Δ::KlLEU2 tdh1Δ tdh2Δ gpm2Δ gpm3Δ eno1Δ pyk2Δ pdc5Δ pdc6Δ adh2Δ adh5Δ adh4Δ sga1Δ::( G tFBA1-FBA1-pFBA1 H pTPI1-TPI1-tTPI1 P tPGK1-PGK1-pPGK1 Q tADH1-ADH1-pADH1 N pPYK1-PYK1-tPYK1 O tTDH3-TDH3-pTDH3 A pENO2-ENO2-tENO2 B pHXK2-HXK2-tHXK2 C pPGI-PGI1-tPGI1 D pPFK1-PFK1-tPFK1 J tPFK2-PFK2-pPFK2 K pAgTEF1-AmdSYM-tAgTEF1 L tGPM1-GPM1-pPGM1 M pPDC1-PDC1-tPDC1-SYN F) pyk1Δ pgi1Δ tpi1Δ tdh3Δ pfk2Δ::(pTEF1-*Spcas9-*tCYC1 natNT1) pgk1Δ gpm1Δ fba1Δ hxk2Δ pfk1Δ adh1Δ pdc1Δ eno2Δ can1Δ:: (ARS418 G pAgTEF-KanMX-tAgTEF AA tFBA1-FBA1-pFBA1 H pTPI1-TPI1-tTPI1 P tPGK1-PGK1-pPGK1 Q tADH1-ADH1-pADH1 N pPYK1-PYK1-tPYK1 O tTDH3-TDH3-pTDH3 A pENO2-ENO2-tENO2 B pHXK2-HXK2-tHXK2 C pPGI-PGI1-tPGI1 D pPFK1-PFK1-tPFK1 J tPFK2-PFK2-pPFK2 L tGPM1-GPM1-pPGM1 M pPDC1-PDC1-tPDC1 AR ARS1211)* | This study |
| **IMX2080** | *MATa ura3-52 his3-1 leu2-3,112 MAL2-8c SUC2 glk1Δ::(pAgTEF1-SpHIS5-tAgTEF1)Δ::(pGAL1-I SceI-tCYC1) hxk1Δ::KlLEU2 tdh1Δ tdh2Δ gpm2Δ gpm3Δ eno1Δ pyk2Δ pdc5Δ pdc6Δ adh2Δ adh5Δ adh4Δ sga1Δ pyk1Δ pgi1Δ tpi1Δ tdh3Δ pfk2Δ::(pTEF1-*Spcas9-*tCYC1 natNT1) pgk1Δ gpm1Δ fba1Δ hxk2Δ pfk1Δ adh1Δ pdc1Δ eno2Δ can1Δ:: (ARS418 G pAgTEF-KanMX-tAgTEF AA tFBA1-FBA1-pFBA1 H pTPI1-TPI1-tTPI1 P tPGK1-PGK1-pPGK1 Q tADH1-ADH1-pADH1 N pPYK1-PYK1-tPYK1 O tTDH3-TDH3-pTDH3 A pENO2-ENO2-tENO2 B pHXK2-HXK2-tHXK2 C pPGI-PGI1-tPGI1 D pPFK1-PFK1-tPFK1 J tPFK2-PFK2-pPFK2 L tGPM1-GPM1-pPGM1 M pPDC1-PDC1-tPDC1 AR ARS1211)* | This study |
| **IMX2109*** | *MATa ura3-52 his3-1 leu2-3,112 MAL2-8c SUC2 glk1Δ::(pAgTEF1-SpHIS5-tAgTEF1)Δ::(pGAL1-I SceI-tCYC1) hxk1Δ::KlLEU2 tdh1Δ tdh2Δ gpm2Δ gpm3Δ eno1Δ pyk2Δ pdc5Δ pdc6Δ adh2Δ adh5Δ adh4Δ sga1Δ pyk1Δ pgi1Δ tpi1Δ tdh3Δ pfk2Δ::(pTEF1-*Spcas9-*tCYC1 natNT1) pgk1Δ gpm1Δ fba1Δ hxk2Δ pfk1Δ adh1Δ pdc1Δ eno2Δ can1Δ:: (ARS418 G pAgTEF-KanMX-tAgTEF AA tFBA1-FBA1-pFBA1 H pTPI1-TPI1-tTPI1 P tPGK1-PGK1-pPGK1 Q tADH1-ADH1-pADH1 N pPYK1-PYK1-tPYK1 O tTDH3-TDH3-pTDH3 A pENO2-ENO2-tENO2 B pHXK2-HXK2-tHXK2 C pPGI-PGI1-tPGI1 D pPFK1-PFK1-tPFK1 J tPFK2-PFK2-pPFK2 L tGPM1-GPM1-pPGM1 M pPDC1-PDC1-tPDC1 AR ARS1211) X2::(pURA3-URA3-tURA3 DT pHIS3-HIS3-tHIS3)* | This study |
| **IMX2059** | *MATa ura3-52 his3-1 leu2-3,112 MAL2-8c SUC2 glk1Δ::(pAgTEF1-SpHIS5-tAgTEF1)Δ::(pGAL1-I SceI-tCYC1) hxk1Δ::KlLEU2 tdh1Δ tdh2Δ gpm2Δ gpm3Δ eno1Δ pyk2Δ pdc5Δ pdc6Δ adh2Δ adh5Δ adh4Δ sga1Δ::( G tFBA1-FBA1-pFBA1 H pTPI1-TPI1-tTPI1 P tPGK1-PGK1-pPGK1 Q tADH1-ADH1-pADH1 N pPYK1-PYK1-tPYK1 O tTDH3-TDH3-pTDH3 A pENO2-ENO2-tENO2 B pHXK2-HXK2-tHXK2 C pPGI-PGI1-tPGI1 D pPFK1-PFK1-tPFK1 J tPFK2-PFK2-pPFK2 K pAgTEF1-AmdSYM-tAgTEF1 L tGPM1-GPM1-pPGM1 M pPDC1-PDC1-tPDC1-SYN F) pyk1Δ pgi1Δ tpi1Δ tdh3Δ pfk2Δ::(pTEF1-*Spcas9-*tCYC1 natNT1) pgk1Δ gpm1Δ fba1Δ hxk2Δ pfk1Δ adh1Δ pdc1Δ eno2Δ X2::(pURA3-URA3-tURA3 DT pHIS3-HIS3-tHIS3)* | This study |
| **IMX2224** | *MATa ura3-52 his3-1 leu2-3,112 MAL2-8c SUC2 hxk1Δ::KlLEU2 tdh1Δ tdh2Δ gpm2Δ gpm3Δ eno1Δ pyk2Δ pdc5Δ pdc6Δ adh2Δ adh5Δ adh4Δ sga1Δ::(FBA1GH TPI1HP PGK1PQ ADH1QN PYK1NO TDH3OA ENO2AB HXK2BC PGI1CD PFK1DJ PFK2JK AmdSYMKL GPM1LM PDC1-SYNMF) pyk1Δ pgi1Δ tpi1Δ tdh3Δ pfk2Δ::(pTEF-cas9-tCYC1 natNT1) pgk1Δ gpm1Δ fba1Δ hxk2Δ pfk1Δ adh1Δ pdc1Δ eno2Δ glk1Δ::Sphis5Δ::(pGAL1-I SceI-tCYC1) X2::pURA3-URA3-tURA3-SHR DT-pHIS3-HIS3-tHIS3 YPRCtau3Δ::pCCW12-mRuby2-tENO1* | This study |

| | | |
|---|---|---|
| **IMX2225** | *MATa ura3-52 his3-1 leu2-3,112 MAL2-8c SUC2 hxk1Δ::KlLEU2 tdh1Δ tdh2Δ gpm2Δ gpm3Δ eno1Δ pyk2Δ pdc5Δ pdc6Δ adh2Δ adh5Δ adh4Δ sga1Δ::(FBA1GH TPI1HP PGK1PQ ADH1QN PYK1NO TDH3OA ENO2AB HXK2BC PGI1CD PFK1DJ PFK2JK AmdSYMKL GPM1LM PDC1-SYNMF) pyk1Δ pgi1Δ tpi1Δ tdh3Δ pfk2Δ::(pTEF-cas9-tCYC1 natNT1) pgk1Δ gpm1Δ fba1Δ hxk2Δ pfk1Δ adh1Δ pdc1Δ eno2Δ glk1Δ::Sphis5Δ::(pGAL1-I SceI-tCYC1) X2::pURA3-URA3-tURA3-SHR DT-pHIS3-HIS3-tHIS3 YPRCtau3Δ::pTEF1-Venus-tTDH1* | This study |
| **IMX2226** | *MATa ura3-52 his3-1 leu2-3,112 MAL2-8c SUC2 hxk1Δ::KlLEU2 tdh1Δ tdh2Δ gpm2Δ gpm3Δ eno1Δ pyk2Δ pdc5Δ pdc6Δ adh2Δ adh5Δ adh4Δ sga1Δ::(FBA1GH TPI1HP PGK1PQ ADH1QN PYK1NO TDH3OA ENO2AB HXK2BC PGI1CD PFK1DJ PFK2JK AmdSYMKL GPM1LM PDC1-SYNMF) pyk1Δ pgi1Δ tpi1Δ tdh3Δ pfk2Δ::(pTEF-cas9-tCYC1 natNT1) pgk1Δ gpm1Δ fba1Δ hxk2Δ pfk1Δ adh1Δ pdc1Δ eno2Δ glk1Δ::Sphis5Δ::(pGAL1-I SceI-tCYC1) X2::pURA3-URA3-tURA3-SHR DT-pHIS3-HIS3-tHIS3 YPRCtau3Δ::pTEF2-mTurquoise2-tSSA1* | This study |
| **IMC111** | *MATa ura3-52 can1 ::cas9-natNT2 TRP1 LEU2 HIS3 pUDC191* | This study |
| **IMC112** | *MATa ura3-52 can1 ::cas9-natNT2 TRP1 LEU2 HIS3 pUDC192* | This study |
| **IMC113** | *MATa ura3-52 can1 ::cas9-natNT2 TRP1 LEU2 HIS3 pUDC193* | This study |
| **IMC153** | *MATa ura3-52 his3-1 leu2-3,112 MAL2-8c SUC2 glk1Δ::(pAgTEF1-SpHIS5-tAgTEF1)Δ::(pGAL1-I SceI-tCYC1) hxk1Δ::KlLEU2 tdh1Δ tdh2Δ gpm2Δ gpm3Δ eno1Δ pyk2Δ pdc5Δ pdc6Δ adh2Δ adh5Δ adh4Δ sga1Δ::( G tFBA1-FBA1-pFBA1 H pTPI1-TPI1-tTPI1 P tPGK1-PGK1-pPGK1 Q tADH1-ADH1-pADH1 N pPYK1-PYK1-tPYK1 O tTDH3-TDH3-pTDH3 A pENO2-ENO2-tENO2 B pHXK2-HXK2-tHXK2 C pPGI-PGI1-tPGI1 D pPFK1-PFK1-tPFK1 J tPFK2-PFK2-pPFK2 K pAgTEF1-AmdSYM-tAgTEF1 L tGPM1-GPM1-pPGM1 M pPDC1-PDC1-tPDC1-SYN F) pyk1Δ pgi1Δ tpi1Δ tdh3Δ pfk2Δ::(pTEF1-Spcas9-tCYC1 natNT1) pgk1Δ gpm1Δ fba1Δ hxk2Δ pfk1Δ adh1Δ pdc1Δ eno2Δ pLM092* | This study |

#### Table S4 SynCh configurations

SHRs are differently annotated than in Kuijpers *et al.* (8). SHRs are annotated in subscript between the genetic fragments that they join together.

| Name | Size | Size chunks | Notes | Stocked name | SynCh configuration |
|---|---|---|---|---|---|
| **SynCh1** | 100 Kb | 2.5 Kb | | IMF1 (wrong), IMF6 | $_{AO'}$ 7A $_{BJ}$ 7B $_{BK}$ 7C $_{BL}$ 7D $_{AP}$ 8A $_{BM}$ 8B $_{BN}$ 8C $_{BO}$ 8D $_{AC}$ *pCCW12-mRuby2-tENO1* $_{AD}$ *CEN6/ARS4* $_{AE}$ 1A $_{AT}$ 1B $_{AS}$ 1C $_{AU}$ 1D $_{AF}$ 2A $_{AV}$ 2B $_{AW}$ 2C $_{AX}$ 2D $_{AG}$ 3A $_{AY}$ 3B $_{AZ}$ 3C $_{BA}$ 3D $_{AH}$ *pTEF2-mTurquoise2-tSSA1* $_{AI}$ ARS417 $_{BU}$ *pHIS3-HIS3-tHIS3* $_{AJ}$ 4A $_{BC}$ 4B $_{BD}$ 4C $_{BE}$ 4D $_{AK}$ 9A $_{BF}$ 9B $_{BS}$ 9C $_{BT}$ 9D $_{AQ}$ 5A $_{BG}$ 5B $_{BH}$ 5C $_{BI}$ 5D $_{AL}$ *pTEF1-Venus-tTDH1* $_{AM}$ ARS1 $_{AN}$ 6A $_{BP}$ 6B $_{BQ}$ 6C $_{BR}$ 6D $_{AO`}$ Telomerator $_{AO`}$ |
| **SynCh2** | 50 Kb | 2.5 Kb | | IMF2 | $_{AO'}$ 7A $_{BJ}$ 7B $_{BK}$ 7C $_{BL}$ 7D $_{AC}$ *pCCW12-mRuby2-tENO1* $_{AD}$ *CEN6/ARS4* $_{AE}$ 1A $_{AT}$ 1B $_{AS}$ 1C $_{AU}$ 1Ds $_{AH}$ *pTEF2 - mTurquoise2 - tSSA1* $_{AI}$ *pHIS3-HIS3-tHIS3* $_{AJ}$ 4A $_{BC}$ 4B $_{BD}$ 4C $_{BE}$ 4D $_{AL}$ *pTEF1 - Venus - tTDH1* $_{AM}$ *ARS1* $_{AN}$ 6A $_{BP}$ 6B $_{BQ}$ 6C $_{BR}$ 6D $_{AO'}$ Telomerator $_{AO'}$ |
| **SynCh3.1** | 100 Kb | 2.5 Kb | | - | $_{AO}$ 7A $_{BJ}$ 7B $_{BK}$ 7C $_{BL}$ 7D $_{AP}$ 8A $_{BM}$ 8B $_{BN}$ 8C $_{BO}$ 8D $_{AC}$ *pCCW12-mRuby2-tENO1* $_{AD}$ *CEN6/ARS4* $_{AE}$ 1A $_{AT}$ 1B $_{AS}$ 1C $_{AU}$ 1D $_{AF}$ 2A $_{AV}$ 2B $_{AW}$ 2C $_{AX}$ 2D $_{AG}$ 3A $_{AY}$ 3B $_{AZ}$ 3C $_{BA}$ 3D $_{AH}$ *pTEF2-mTurquoise2-tSSA1* $_{AI}$ ARS417 $_{BU}$ *pHIS3-HIS3-tHIS3* $_{AJ}$ 4A $_{BC}$ 4B $_{BD}$ 4C $_{BE}$ 4D $_{AK}$ 9A $_{BF}$ 9B $_{BS}$ 9C $_{BT}$ 9D $_{AQ}$ 5A $_{BG}$ 5B $_{BH}$ 5C $_{BI}$ 5D $_{AL}$ *pTEF1-Venus-tTDH1* $_{AM}$ ARS1 $_{AN}$ 6A $_{BP}$ 6B $_{BQ}$ 6C $_{BR}$ 6D $_{AO}$ |
| **SynCh3.2** | 100 Kb | 5 Kb | | - | $_{AO}$ 7A $_{BJ}$ 7B $_{BK}$ 7C $_{BL}$ 7D $_{AP}$ 8A $_{BM}$ 8B $_{BN}$ 8C $_{BO}$ 8D $_{AC}$ *pCCW12-mRuby2-tENO1* $_{AD}$ *CEN6/ARS4* $_{AE}$ 1A $_{AT}$ 1B $_{AS}$ 1C $_{AU}$ 1D $_{AF}$ 2A $_{AV}$ 2B $_{AW}$ 2C $_{AX}$ 2D $_{AG}$ 3A $_{AY}$ 3B $_{AZ}$ 3C $_{BA}$ 3D $_{AH}$ *pTEF2-mTurquoise2-tSSA1* $_{AI}$ ARS417 $_{BU}$ *pHIS3-HIS3-tHIS3* $_{AJ}$ 4A $_{BC}$ 4B $_{BD}$ 4C $_{BE}$ 4D $_{AK}$ 9A $_{BF}$ 9B $_{BS}$ 9C $_{BT}$ 9D $_{AQ}$ 5A $_{BG}$ 5B $_{BH}$ 5C $_{BI}$ 5D $_{AL}$ *pTEF1-Venus-tTDH1* $_{AM}$ ARS1 $_{AN}$ 6A $_{BP}$ 6B $_{BQ}$ 6C $_{BR}$ 6D $_{AO}$ |
| **SynCh3.3** | 100 Kb | 10 Kb | | - | $_{AO}$ 7A $_{BJ}$ 7B $_{BK}$ 7C $_{BL}$ 7D $_{AP}$ 8A $_{BM}$ 8B $_{BN}$ 8C $_{BO}$ 8D $_{AC}$ *pCCW12-mRuby2-tENO1* $_{AD}$ *CEN6/ARS4* $_{AE}$ 1A $_{AT}$ 1B $_{AS}$ 1C $_{AU}$ 1D $_{AF}$ 2A $_{AV}$ 2B $_{AW}$ 2C $_{AX}$ 2D $_{AG}$ 3A $_{AY}$ 3B $_{AZ}$ 3C $_{BA}$ 3D $_{AH}$ *pTEF2-mTurquoise2-tSSA1* $_{AI}$ ARS417 $_{BU}$ *HIS3* $_{AJ}$ 4A $_{BC}$ 4B $_{BD}$ 4C $_{BE}$ 4D $_{AK}$ 9A $_{BF}$ 9B $_{BS}$ 9C $_{BT}$ 9D $_{AQ}$ 5A $_{BG}$ 5B $_{BH}$ 5C $_{BI}$ 5D $_{AL}$ *pTEF1-Venus-tTDH1* $_{AM}$ ARS1 $_{AN}$ 6A $_{BP}$ 6B $_{BQ}$ 6C $_{BR}$ 6D $_{AO}$ |
| **SynCh4.1** | 50 Kb | 2.5 Kb | | - | $_{AO}$ 7A $_{BJ}$ 7B $_{BK}$ 7C $_{BL}$ 7D $_{AC}$ *pCCW12-mRuby2-tENO1* $_{AD}$ *CEN6/ARS4* $_{AE}$ 1A $_{AT}$ 1B $_{AS}$ 1C $_{AU}$ 1Ds $_{AH}$ *pTEF2 - mTurquoise2 - tSSA1* $_{AI}$ *pHIS3-HIS3-tHIS3* $_{AJ}$ 4A $_{BC}$ 4B $_{BD}$ 4C $_{BE}$ 4D $_{AL}$ *pTEF1 - Venus - tTDH1* $_{AM}$ *ARS1* $_{AN}$ 6A $_{BP}$ 6B $_{BQ}$ 6C $_{BR}$ 6D $_{AO}$ |
| **SynCh4.2** | 50 Kb | 5 Kb | | - | $_{AO}$ 7A $_{BJ}$ 7B $_{BK}$ 7C $_{BL}$ 7D $_{AC}$ *pCCW12-mRuby2-tENO1* $_{AD}$ *CEN6/ARS4* $_{AE}$ 1A $_{AT}$ 1B $_{AS}$ 1C $_{AU}$ 1Ds $_{AH}$ *pTEF2 - mTurquoise2 - tSSA1* $_{AI}$ *pHIS3-HIS3-tHIS3* $_{AJ}$ 4A $_{BC}$ 4B $_{BD}$ 4C $_{BE}$ 4D $_{AL}$ *pTEF1 - Venus - tTDH1* $_{AM}$ *ARS1* $_{AN}$ 6A $_{BP}$ 6B $_{BQ}$ 6C $_{BR}$ 6D $_{AO}$ |

| | | | | | |
|---|---|---|---|---|---|
| **SynCh4.3** | 50 Kb | 10 Kb | | - | AO 7A BJ 7B BK 7C BL 7D AC *pCCW12-mRuby2-tENO1* AD *CEN6/ARS4* AE 1A AT 1B AS 1C AU 1Ds AH *pTEF2 - mTurquoise2 - tSSA1*AI *pHIS3-HIS3-tHIS3*AJ 4A BC 4B BD 4C BE 4D AL *pTEF1 - Venus - tTDH1* AM *ARS1* AN 6A BP 6B BQ 6C BR 6D AO |
| **Syn12** | 100 Kb | 2.5 Kb | | IMF23 (col. 8), IMF24 (col. 1), IMF25 (col. 3), IMF26 (col. 4) | AO 7A BJ 7B BK 7C BL 7D AP 8A BM 8B BN 8C BO 8D AC *pCCW12-mRuby2-tENO1* AD *ARS1*AN 18A BP 18B BQ 19C BR 19D DE 15A DF 15B DH 15C DI 15D DJ *CEN6/ARS4*AE 16A DK 16B DL 16C DM 16D DN 17A DO 17B DP 19A DQ 17D DR *ARS417* BU *pHIS3-HIS3-tHIS3*AJ 4A BC 4B BD 4C BE 4D AK 9A BF 9B BS 9C BT 9D AQ 5A BG 5B BH 5C BI 5D AL *tSSA1-mTurquoise2-pTEF2* DS Telomerator AO |
| **Syn15** | 135 Kb | 2.5 Kb | Glycolysis (800bp promoters and 300bp terminators) integrated by CRISPR/Cas 9 at *mTurquoise2* of IMF6 | IMF13 (double glycolysis), IMF17 (single glycolysis) | AO' 7A BJ 7B BK 7C BL 7D AP 8A BM 8B BN 8C BO 8D AC *pCCW12-mRuby2-tENO1* AD *CEN6/ARS4*AE 1A AT 1B AS 1C AU 1D AF 2A AV 2B AW 2C AX 2D AG 3A AY 3B AZ 3C BA 3D AH ARS418 G *pAgTEF-KanMX-tAgTEF* AA *tFBA1-FBA1-pFBA1* H *pTPI1-TPI1-tTPI1* P *tPGK1-PGK1-pPGK1* Q *tADH1-ADH1-pADH1* N *pPYK1-PYK1-tPYK1* O *tTDH3-TDH3-pTDH3* A *pENO2-ENO2-tENO2* B *pHXK2-HXK2-tHXK2* C *pPGI-PGI1-tPGI1* D *pPFK1-PFK1-tPFK1* J *tPFK2-PFK2-pPFK2* L *tGPM1-GPM1-pPGM1* M *pPDC1-PDC1-tPDC1* AR *ARS1211* BU *pHIS3-HIS3-tHIS3*AJ 4A BC 4B BD 4C BE 4D AK 9A BF 9B BS 9C BT 9D AQ 5A BG 5B BH 5C BI 5D AL *pTEF1-Venus-tTDH1*AM ARS1 AN 6A BP 6B BQ 6C BR 6D AO` Telomerator AO` |
| **Syn16** | 85 Kb | 2.5 Kb | Glycolysis (800bp promoters and 300bp terminators) integrated by CRISPR/Cas 9 at *mTurquoise2* of IMF2 | IMF14 (double glycolysis), IMF18 (single glycolysis) | AO' 7A BJ 7B BK 7C BL 7D AC *pCCW12-mRuby2-tENO1* AD *CEN6/ARS4* AE 1A AT 1B AS 1C AU 1Ds AH *ARS418* G *pAgTEF-KanMX-tAgTEF* AA *tFBA1-FBA1-pFBA1* H *pTPI1-TPI1-tTPI1* P *tPGK1-PGK1-pPGK1* Q *tADH1-ADH1-pADH1* N *pPYK1-PYK1-tPYK1* O *tTDH3-TDH3-pTDH3* A *pENO2-ENO2-tENO2* B *pHXK2-HXK2-tHXK2* C *pPGI-PGI1-tPGI1* D *pPFK1-PFK1-tPFK1* J *tPFK2-PFK2-pPFK2* L *tGPM1-GPM1-pPGM1* M *pPDC1-PDC1-tPDC1* AR *ARS1211*AI *pHIS3-HIS3-tHIS3*AJ 4A BC 4B BD 4C BE 4D AL *pTEF1 - Venus - tTDH1* AM *ARS1* AN 6A BP 6B BQ 6C BR 6D AO' Telomerator AO' |
| **Syn19** | 135 Kb | 2.5 Kb | *E.coli* chunks integrated by CRISPR/Cas 9 at *mTurquoise2* of IMF6 | IMF11 | AO' 7A BJ 7B BK 7C BL 7D AP 8A BM 8B BN 8C BO 8D AC *pCCW12-mRuby2-tENO1* AD *CEN6/ARS4*AE 1A AT 1B AS 1C AU 1D AF 2A AV 2B AW 2C AX 2D AG 3A AY 3B AZ 3C BA 3D AH ARS418 G *pAgTEF-KanMX-tAgTEF* AA 15A DF 15B DH 15C DI 15D DJ 16A DK 16B DL 16C DM 16D DN 17A DO 17B DP 19A DQ 17D DR 20A AR ARS1211 BU *pHIS3-HIS3-tHIS3*AJ 4A BC 4B BD 4C BE 4D AK 9A BF 9B BS 9C BT 9D AQ 5A BG 5B BH 5C BI 5D AL *pTEF1-Venus-tTDH1*AM ARS1 AN 6A BP 6B BQ 6C BR 6D AO` Telomerator AO` |

| | | | | | |
|---|---|---|---|---|---|
| **Syn20** | 85 Kb | 2.5 Kb | *E.coli* chunks integrated by CRISPR/Cas 9 at *mTurquoise2* of IMF2 | IMF12 | AO' 7A BJ 7B BK 7C BL 7D AC *pCCW12-mRuby2-tENO1* AD *CEN6/ARS4* AE 1A AT 1B AS 1C AU 1Ds AH ARS418 G *pAgTEF-KanMX-tAgTEF* AA 15A DF 15B DH 15C DI 15D DJ 16A DK 16B DL 16C DM 16D DN 17A DO 17B DP 19A DQ 17D DR 20A AR ARS1211 AI *pHIS3-HIS3-tHIS3* AJ 4A BC 4B BD 4C BE 4D AL *pTEF1 - Venus - tTDH1* AM *ARS1* AN 6A BP 6B BQ 6C BR 6D AO' Telomerator AO' |

#### Table S5 Primers to construct yeast toolkit plasmids.

| Primer Name | Sequence 5' - 3' |
|---|---|
| HXK2 sc prom fw Ytoolkit | AAGCATCGTCTCATCGGTCTCAAACGCTGGTAAAGTACAGCTACATTC |
| YTK HXK2 REV | ATGCCGTCTCAGGTCTCACAGCACGCTACAAAAGAAAGTACGCAAG |
| PGI1 sc prom fw Ytoolkit | AAGCATCGTCTCATCGGTCTCAAACGTATTCTTAGTGGATAACATGCG |
| PGI1 sc term rv Ytoolkit | TTATGCCGTCTCAGGTCTCACAGCGAAATAGGACCTGATATCCTCC |
| PFK1 sc prom fw Ytoolkit | AAGCATCGTCTCATCGGTCTCAAACGCGGCTAGTAAAAAAGAAAATTAATATCTCATTAAC |
| YTK PFK1 REV | ATGCCGTCTCAGGTCTCACAGCCACATTCAGAGCAATTTGTAGTAC |
| PFK2 sc prom fw Ytoolkit | AAGCATCGTCTCATCGGTCTCAAACGCCATTCTCTGCTGCTTTGTTG |
| YTK PFK2 REV | ATGCCGTCTCAGGTCTCACAGCATAAGAGAACAAAGTATTTAACGC |
| FBA1 sc prom fw Ytoolkit | AAGCATCGTCTCATCGGTCTCAAACGCAATACCAGCCTTCCAACTTC |
| FBA1 sc term rv Ytoolkit | TTATGCCGTCTCAGGTCTCACAGCCGCGAACTCCAAAATGAGC |
| TPI1 sc prom fw Ytoolkit | AAGCATCGTCTCATCGGTCTCAAACGACCCAGAGATGTTGTTGTCC |
| TPI1 sc term rv Ytoolkit | TTATGCCGTCTCAGGTCTCACAGCCGGTACACTTCTGAGTAAC |
| TDH3 sc prom fw Ytoolkit | AAGCATCGTCTCATCGGTCTCAAACGCGAATATATACTAGCGTTGAATGTTAG |
| TDH3 sc term rv Ytoolkit | TTATGCCGTCTCAGGTCTCACAGCGTAACTTCAGAATCGTTATCCTGG |
| PGK1 sc prom fw Ytoolkit | AAGCATCGTCTCATCGGTCTCAAACGTATTTTAGATTCCTGACTTCAACTC |
| PGK1 sc term rv Ytoolkit | TTATGCCGTCTCAGGTCTCACAGCCGAAATAATATCCTTCTCGAAAG |
| pGPM1 sc fw Ytookit | AAGCATCGTCTCATCGGTCTCAAACGGTGATACTTTGACAGGAGC |
| GPM1 sc term rv Ytoolkit | TTATGCCGTCTCAGGTCTCACAGCCATTAAACTACGATGTAAACATC |
| ENO2 sc prom fw Ytoolkit | AAGCATCGTCTCATCGGTCTCAAACGGGATGATGAAAACACTAAACGAAG |
| YTK ENO2 REV | ATGCCGTCTCAGGTCTCACAGCAGGTATCATCTCCATCTCCC |
| PYK1 sc prom fw Ytoolkit | AAGCATCGTCTCATCGGTCTCAAACGCCCTGGTCAAACTTCAGAAC |
| PYK1 sc term rev Ytoolkit | TTATGCCGTCTCAGGTCTCACAGCGTATCCTTTCGCCATCCTG |
| PDC1 sc prom fw Ytoolkit | AAGCATCGTCTCATCGGTCTCAAACGCATGCGACTGGGTGAGCATATG |
| PDC1 rv term rv Ytoolkit | TTATGCCGTCTCAGGTCTCACAGCCAGTGTTCCTTAATCAAGGATACC |
| ADH1 sc prom fw Ytoolkit | AAGCATCGTCTCATCGGTCTCAAACGAAGTCCAATGCTAGTAGAGAAG |
| ADH1 sc term rv Ytoolkit | TTATGCCGTCTCAGGTCTCACAGCCAACAGGTGTTGTCCTCTG |

#### Table S6 Primers used to construct guide RNA plasmids

| Primer Name | Sequence 5' - 3' |
|---|---|
| mTurquoise2_gRNA1_fw | TGCGCATGTTTCGGCGTTCGAAACTTCTCCGCAGTGAAAGATAAATGATCTACTGCTGCTGGTATTACCTGTTTTAGAGCTAGAAATAGCAAGTTAAAATAAG |
| mTurquoise2_gRNA2_fw | TGCGCATGTTTCGGCGTTCGAAACTTCTCCGCAGTGAAAGATAAATGATCCCTTAGTCACTACTTTATCTGTTTTAGAGCTAGAAATAGCAAGTTAAAATAAG |
| CRISPR RNA Recycl fw | TGCGCATGTTTCGGCGTTCGAAACTTCTCCGCAGTGAAAGATAAATGATCTTACAATATAGTGATAATCGGTTTTAGAGCTAGAAATAGCAAGTTAAAATAAGGCTAGTCCGTTATCAAC |
| FW_X-2_gRNA | TGCGCATGTTTCGGCGTTCGAAACTTCTCCGCAGTGAAAGATAAATGATCGGCGACTAGGAAGAGAGTAGGTTTTAGAGCTAGAAATAGCAAGTTAAAATAAG |
| YPRCtau3_targetRNA FW | TGCGCATGTTTCGGCGTTCGAAACTTCTCCGCAGTGAAAGATAAATGATCAAACATTCAAATATATTCCAGTTTTAGAGCTAGAAATAGCAAGTTAAAATAAG |

#### Table S7 Primers for *pGAL1-I SceI-tCYC1* expression cassette

Flanks for homologous recombination are marked in bold.

| Primer Name | Sequence 5’ - 3’ |
|---|---|
| glk1oh_prGAL1_FW | **AAACCACAACACCACCACTAATACAACTCTATCATACACAAGATG**GCAGTGAGCGCAACGCAATTAATG |
| glk1oh_terCYC1_RV | **GTACGGTGGGATACGTACACAAACCAAAAAAATGTAAAAAGATCA**CGACTCACTATAGGGCGAATTGG |
| **Diagnostic primers to check integration in genome** | |
| GLK1FW4 | GCCCGACAGGGTAACATATTATC |
| I-SceI inside rv | GAACCAGTATGCCAGAGACATC |
| GAL1fw1 | GTTCCTGAAACGCAGATGTG |

#### Table S8 Primers used for fragments for IMF1/IMF6 assembly

SHRs are marked in bold.

| Primer Name | Sequence 5’ - 3’ |
|---|---|
| Ecoli_ch1_fw | **ATGTTCGGAAGAGCTGTCTCTATCTGGTACGAGTGCTGCTGTCATCATCCTCATGCGTTG**GTGCCGATGAAGTTGACGTG |
| Chunk_1A_rv | **CCTGACACTGCGTCGAGATATATAGGATTGCTGAAGAGCCGATCGCAGTTCAGAGCGCAT**CGGAGTCAGCGGAGGTATAG |
| Chunk_1B_fw | **ATGCGCTCTGAACTGCGATCGGCTCTTCAGCAATCCTATATATCTCGACGCAGTGTCAGG**CCAGATTGCCTGCCACGAAG |
| Chunk_1B_rv | **GCGGTATGTCGTGCTATAATAGCATACATATACTCGCCGTCGGCAATATGATGATAGGTC**GTTCCCGCGTGACCAACCTC |
| Chunk_1C_fw | **GACCTATCATCATATTGCCGACGGCGAGTATATGTATGCTATTATAGCACGACATACCGC**TGCCTTTCGCCGCCAGTTTC |
| Chunk_1C_rv | **GAGGATCATGTCTGTCACTCGCATCGTGTCTAGTGATATGCGTATCTCTGATGAGTGATG**AAGACGCACCTCGGTGATGG |
| Chunk_1D_fw | **CATCACTCATCAGAGATACGCATATCACTAGACACGATGCGAGTGACAGACATGATCCTC**CCAACGGTGGCGCGTAAC |
| Ecoli_ch1_rv | **TGAGAGCTTGTGATAACTGCTCGCCAGTTGTGGTGATCTCCCAGTCGGTGTAGCAGCAAT**TAATCTCGACAGCCTCTTCC |
| Ecoli_ch2_fw | **ATTGCTGCTACACCGACTGGGAGATCACCACAACTGGCGAGCAGTTATCACAAGCTCTCA**CAAACTGCTCGTTCAGTACTTCACCCA |
| Chunk_2A_rv | **CCGCTATCTGCGCGTAGTATCATGATTACAACCTATGCGCTTTGAGATTTGTCGAGAGCC**GTCGCCGTTATCAGCAGCAA |
| Chunk_2B_fw | **GGCTCTCGACAAATCTCAAAGCGCATAGGTTGTAATCATGATACTACGCGCAGATAGCGG**AAATGGCACCGCCGCAGATC |
| Chunk_2B_rv | **TGAACAGTTGTATTGCTGCCATCCATGTCACGACCTATGAGGATGACTTCGATGTAAGCT**GCGCCGCCATAGCGAAGTTG |
| Chunk_2C_fw | **AGCTTACATCGAAGTCATCCTCATAGGTCGTGACATGGATGGCAGCAATACAACTGTTCA**GTGGCGAAATCGAAGTACCG |
| Chunk_2C_rv | **GCGCATGATTCAGATGGTCCTAGCGGCATTGGCGAAATCTCGTAGCGATGGTATCAGTCT**TTTGACGGCTGTAAGCGCTCAC |
| Chunk_2D_fw | **AGACTGATACCATCGCTACGAGATTTCGCCAATGCCGCTAGGACCATCTGAATCATGCGC**GATGGCGCAAATTCTTCAATGGATCG |
| Ecoli_ch2_rv | **GGTGAATTGAGAGCTATCCTATATTATAGCAGATGCCGGGTATGCAGCTTGGTAGAATGC**CCAGTGATGTTGAATATTACTTTGAACCAGTGAC |
| Ecoli_ch3_fw | **GCATTCTACCAAGCTGCATACCCGGCATCTGCTATAATATAGGATAGCTCTCAATTCACC**CGCGTAAATGTTGCGATGTAATATTCCGT |
| Chunk_3A_rv | **CTACTAAATGTGAGCAGGTGGCATCCTGTGCTCGCATATAATGAGAGCTGATGCCGCACT**GGCGGTTATCAGTGAAATGGTACATCG |
| Chunk_3B_fw | **AGTGCGGCATCAGCTCTCATTATATGCGAGCACAGGATGCCACCTGCTCACATTTAGTAG**GGAACTTCCCAGAGGATGACTTCAC |
| Chunk_3B_rv | **ATGGCCGCTCGTATCGCAATCACAGATGGCGGTCGATACATTTCGTGCTTTGGAGACAGC**TTCATTGACAGAAGGCAGCGTCAG |
| Chunk_3C_fw | **GCTGTCTCCAAAGCACGAAATGTATCGACCGCCATCTGTGATTGCGATACGAGCGGCCAT**TGAATTGTTCGATATCATCGACCCGATTG |
| Chunk_3C_rv | **TAAGTCTCTTGACATCTCGGAACATATCCACTCAGCGGTGTATCATTCTGTGGTCGGCGC**GCTTGGCATCGACAGTTTTGTG |
| Chunk_3D_fw | **GCGCCGACCACAGAATGATACACCGCTGAGTGGATATGTTCCGAGATGTCAAGAGACTTA**CGGCATATTCGTTGATTCGCGCAG |
| Ecoli_ch3_rv | **CTCAGCCTTAGCCAATATGATCATGTCGTTGCGTCTCGGACCATCTAGTCTACTCTGAAG**TAAATGGCCTGGATGTAGAGGTTTATCACTG |
| prTEF2_Turquoise2_tSSA1_fw | **CTTCAGAGTAGACTAGATGGTCCGAGACGCAACGACATGATCATATTGGCTAAGGCTGAG**GAACGTTGATAGGTCAAGATCAATG |
| prTEF2_Turquoise2_tSSA1_rv | **CAGTGACGTGAGTGCCATCTGCAGGTCATGTGATGCTATCAGCTACACTGCCAGCAATGA**CGTCTCATTGGCAGCATAAA |

| | |
|---|---|
| **ARS417_fw** | **TCATTGCTGGCAGTGTAGCTGATAGCATCACATGACCTGCAGATGGCACTCACGTCACTG**ACAAGTCCTTAAGAATATACAAAAAGCTACATAAATATAA |
| **ARS417_rv** | **AATCATGTGACCCAGGCTTGCGCATACATGATCCTTCTTGCGCTGCATGGGCGACTATAT**CTTACGCTCAATTCCTTTATTTTTTTATATTTATGTAGCTTTTT |
| **BU-His3_fw** | **ATATAGTCGCCCATGCAGCGCAAGAAGGATCATGTATGCGCAAGCCTGGGTCACATGATT**TGCGGCATCAGAGCAGATTG |
| **terHIS3_rv** | **ACGCAATATCGGCCATCGTGCGAGTGTCTCAAACTATCTGTATGCAAATTCGTGCGTGTG**GCATCTGTGCGGTATTTCACAC |
| **Ecoli_ch4_fw** | **CACACGCACGAATTTGCATACAGATAGTTTGAGACACTCGCACGATGGCCGATATTGCGT**CCGCGCATATTCCAGCAAGGAG |
| **Chunk_4A_rv** | **CTAGGCTCTGCTGCATGTCAGTGATTTCTATTAGGCAGCGCTTACCCATGATTAGCGCAG**CTACGGTCAGTTCGCCTTTC |
| **Chunk_4B_fw** | **CTGCGCTAATCATGGGTAAGCGCTGCCTAATAGAAATCACTGACATGCAGCAGAGCCTAG**GTGTCAGCTTTCGTGGTGTG |
| **Chunk_4B_rv** | **AGTCACGCTGAGTCCATGCTGACCATGATTCACACTCAGTGCCGATAATTCCATAGTCTG**CTCTCCGGCATTGACGGAAC |
| **Chunk_4C_fw** | **CAGACTATGGAATTATCGGCACTGAGTGTGAATCATGGTCAGCATGGACTCAGCGTGACT**GCCAATTTCCGTGTTGTAGG |
| **Chunk_4C_rv** | **TCAATCATTCGTTCTCGCAGATCTACAATCGTCCTGAGCTCTGTGAGTGATGTACGCTCC**CTAACGCGTCAATGCACTCC |
| **Chunk_4D_fw** | **GGAGCGTACATCACTCACAGAGCTCAGGACGATTGTAGATCTGCGAGAACGAATGATTGA**TACCGTCACGCCACGTCCAC |
| **Ecoli_ch4_rv** | **TCAGACAATTCTATACGCGGACTGATATGGCAGAAGCTAGGAGACGTTATGCGATCTTAG**TGGTACGGTTTACTCCTTCACCTGTG |
| **Ecoli_ch9_fw** | **CTAAGATCGCATAACGTCTCCTAGCTTCTGCCATATCAGTCCGCGTATAGAATTGTCTGA**CATTACCGGCGTAGCCCCATG |
| **Chunk_9.2A_rv** | **GCGCGACGTGTCTCGTATATTAGTGAAGTTGGATCTGTCCATGAATCCTCGGCTCTGGTG**TGCAAGCGGTATGAGGAAAG |
| **Chunk_9.2B_fw** | **CACCAGAGCCGAGGATTCATGGACAGATCCAACTTCACTAATATACGAGACACGTCGCGC**TATACTGCGGGTAGGAAAGG |
| **Chunk_9.2B_rv** | **GTTCAGGATTCTGTCGATGCCACATCGAGTCAGTCGTAGTAACATGGAACGCAGTGCATC**TCACGTTCTGGTATTGGGTGC |
| **Chunk_9.2C_fw** | **GATGCACTGCGTTCCATGTTACTACGACTGACTCGATGTGGCATCGACAGAATCCTGAAC**GCGTCGCTTTACGCCAGGTC |
| **Chunk_9.2C_rv** | **CAGATACTGGGCAGGCTCTATAGGAGCTTGTACCGCATTGGCTTTGCCACTCATTCGAGA**GCTGCCGCCGATGAGATCGC |
| **Chunk_9.2D_fw** | **TCTCGAATGAGTGGCAAAGCCAATGCGGTACAAGCTCCTATAGAGCCTGCCCAGTATCTG**GACTATCTGCTGACTGAGTTGCTGTTG |
| **Chunk_9.2D_rv** | **ATAAGGAATGATGCACGCGCGACGCTGCTTCAATCAGATGTATAGCATTGCCTTCTGCGC**GGTTGCATACTGTGGCAACTGAC |
| **Ecoli_ch5_fw** | **GCGCAGAAGGCAATGCTATACATCTGATTGAAGCAGCGTCGCGCGTGCATCATTCCTTAT**CTGGGGAAACTGGCGAGCG |
| **Chunk_5A_rv** | **GAGGCTTCACAGTGCTTTATTAGTATGATTGCCTAGCTGGTATATGTGTTCCTGGAGCGC**TGTGGATCTGGCGGTTACGG |
| **Chunk_5B_fw** | **GCGCTCCAGGAACACATATACCAGCTAGGCAATCATACTAATAAAGCACTGTGAAGCCTC**GCTGATTACCGCAGCCTGAA |
| **Chunk_5B_rv** | **AGGATCGCTCGCGTACTCATGCATTCTCCCACATATTGAGGCCCTGATTCCATGCAATGT**GGAAAATCTCCGCCATTCCC |
| **Chunk_5C_fw** | **ACATTGCATGGAATCAGGGCCTCAATATGTGGGAGAATGCATGAGTACGCGAGCGATCCT**ATGGCTTACGGCAGCATTGG |
| **Chunk_5C_rv** | **TCTGTCAGTTGGTTAAGCGCCGCTACGATTACTACACATGCCACAGACTGATCTACAATG**GTACCGCTCTGCACCACAGG |
| **Chunk_5D_fw** | **CATTGTAGATCAGTCTGTGGCATGTGTAGTAATCGTAGCGGCGCTTAACCAACTGACAGA**ACAAAGTACCGCCAGCCAGG |
| **Ecoli_ch5_rv** | **GCGTCCTCTGTATTAAGATGTCATGGTGTTGAGTCTGCACATGCAATGGCAATGAGTCAG**TTAGCCCGCAGTGGATCCTCC |

| | |
|---|---|
| **prTEF1_Venus_tENO2_fw** | **CTGACTCATTGCCATTGCATGTGCAGACTCAACACCATGACATCTTAATACAGAGGACGC**CCTTGCCAACAGGGAGTTCTTCAG |
| **prTEF1_Venus_tTDH1_rv** | **CAGTGACATGCCGCTCAGTACTCGTATCTTACATGACGTGGGCATGGGTTCCGCTCATAT**TGGCAGCCGTTCAGGGTAAT |
| **ARS1_fw** | **ATATGAGCGGAACCCATGCCCACGTCATGTAAGATACGAGTACTGAGCGGCATGTCACTG**GGCCTTTTGAAAAGCAAGCATAAAAGATCTAAAC |
| **ARS1_rv** | **ATACATCATGCACGCCTGAAAGCATCCCTGACGCGAGTATGACGCAGTTCACACATCTTA**CTTGTTATTTTACAGATTTTATGTTTAGATCTTTTATGCTTGCTTTTCAAAA |
| **Ecoli_ch6_fw** | **TAAGATGTGTGAACTGCGTCATACTCGCGTCAGGGATGCTTTCAGGCGTGCATGATGTAT**CCCGCTACGAAATTGCGGATACTAAGG |
| **Chunk_6A_rv** | **GAGATGACTGGGTCCACTCTTTCGTGTATTTCGAGAGAGCGATACGCATGTCTCCATCGT**TTCACTCTGCCCGGTGTTCC |
| **Chunk_6B_fw** | **ACGATGGAGACATGCGTATCGCTCTCTCGAAATACACGAAAGAGTGGACCCAGTCATCTC**CGAAAGAGCTGTGGCTAACC |
| **Chunk_6B_rv** | **CAGATCAGTGTCAATGAAGGTAGGCTGCTTGGCAATGCTTCTGGTGACTGGTAGATCATC**GTCTGTGCGACCACCAATCCC |
| **Chunk_6C_fw** | **GATGATCTACCAGTCACCAGAAGCATTGCCAAGCAGCCTACCTTCATTGACACTGATCTG**CTAGTCCGCCACGGTGAAACG |
| **Chunk_6C_rv** | **AGCTGGCGTCGCGCATAAATGCATGATCTGTCCTGGCTGACACGCATCTGCACTAATGAT**TAGTCGTCAGCGCCAATTTC |
| **Chunk_6D_fw** | **ATCATTAGTGCAGATGCGTGTCAGCCAGGACAGATCATGCATTTATGCGCGACGCCAGCT**CGTCGGGTGAAGAAGTTCTC |
| **Ecoli_ch6_rv** | **TGACGCGCTTTCAACCATGTTGCTGGGATTTCAGATTAGCCACGTACCTGGCATCAATGA**GGCCATATTCAGCTTCAGGTTCTGCTTC |
| **Telomerator_pos7n_fw** | **CGGAAGCAGAACCTGAAGCTGAATATGGCCTCATTGATGCCAGGTACGTGGCTAATCTGA**CCCGGGGGATCCGGTGATTG |
| **Telomerator_pos7_rv** | **CGCCTGGAGAATGAGCGGGAAGCTAACCATTGACGCGCTTTCAACCATGTTGCTGGGATT**CCAAAGCTGGAGCTCCACCG |
| **Ecoli_ch7_fw** | **TCATTGATGCCAGGTACGTGGCTAATCTGAAATCCCAGCAACATGGTTGAAAGCGCGTCA**ATGGTTAGCTTCCCGCTCATTCTCC |
| **Chunk_7A_rv** | **GGCGCACATGGTATATTATGATCGGAGATGCGGCAACATAGCTGGGTGTGATCCTCTCTA**CGCCATCCGTGGGTCTTTTC |
| **Chunk_7B_fw** | **TAGAGAGGATCACACCCAGCTATGTTGCCGCATCTCCGATCATAATATACCATGTGCGCC**TGGGCGTCATTGTCCGGAGT |
| **Chunk_7B_rv** | **GAGCATACTGTCCTATCATGTCGACTCTTGTCACATCTGACGCCTCTCTGCGATAGGATT**TCCGGCGGCAGCCATCAAAG |
| **Chunk_7C_fw** | **AATCCTATCGCAGAGAGGCGTCAGATGTGACAAGAGTCGACATGATAGGACAGTATGCTC**CCAGGCGGAAGAAGTCTTTGAAGAC |
| **Chunk_7C_rv** | **CAGCAAGTGCGTAGAGATCAGCATTATCTGACTGTGGATGATCCTACATCGTCATCAGAG**CGCCGCTTCATAAGCGCCAA |
| **Chunk_7D_fw** | **CTCTGATGACGATGTAGGATCATCCACAGTCAGATAATGCTGATCTCTACGCACTTGCTG**AATGGCGATCCCCGAGCAAC |
| **Ecoli_ch7_rv** | **CTCAAGGCTGTGCTGACGTAGGACTGATTCGAGCTATCTCTTGGTGTATTTGAGCAGACC**CAGACATGGCGTAACCCCGTG |
| **Ecoli_ch8_fw** | **GGTCTGCTCAAATACACCAAGAGATAGCTCGAATCAGTCCTACGTCAGCACAGCCTTGAG**GCATTTACGCTGTACGGACACCTT |
| **Chunk_8A_rv** | **ACAATGAGAATCGAGCGCCGCTGCTTAATCTGTCAGTCGATCCTATGGTTGCTGCTGAGC**ACAGTGCAGCGCGTTTGGTC |
| **Chunk_8B_fw** | **GCTCAGCAGCAACCATAGGATCGACTGACAGATTAAGCAGCGGCGCTCGATTCTCATTGT**CTGGCGGGTTACTGGCTGTG |
| **Chunk_8B_rv** | **GGCCGCTGTGTAGTCTCTATGCATGTACTTAGATCCTAGCGCATCTTCGCCAGCTATATT**TGCCGACTTACGCCGTGGTT |
| **Chunk_8C_fw** | **AATATAGCTGGCGAAGATGCGCTAGGATCTAAGTACATGCATAGAGACTACACAGCGGCC**TGATCGGACTGGGCGATCAC |

| | |
|---|---|
| **Chunk_8C_rv** | **GGCATTCGGCGTGATTCCATCATGCTATGCACTGATCTCGCACATAATCTCGGTCGGCTG**CGGTATGACCCTGGCGGAAG |
| **Chunk_8D_fw** | **CAGCCGACCGAGATTATGTGCGAGATCAGTGCATAGCATGATGGAATCACGCCGAATGCC**TTTCAGCGTGCTCTGTTTACCC |
| **Ecoli_ch8_rv** | **GCTACATCTTCCGTACTATGCTGTAGTCTCATGGTCGAGTTCTATTGCTGTTCGGCGGCA**GGGCAATATTACCAACCCGTTTTGC |
| **prCCW12_mRuby_tENO1_fw** | **TGCCGCCGAACAGCAATAGAACTCGACCATGAGACTACAGCATAGTACGGAAGATGTAGC**AACGCACCCATGAACCACAC |
| **prCCW12_mRuby_tENO1_rv** | **CTCCACTGTACTGCATGTAGCATTCGCCGATCTGCATGATGTGTGACATTCTGCTATCGG**GGCAGCATACATGGGTGACCAAA |
| **CEN6_ARS4_fw** | **CCGATAGCAGAATGTCACACATCATGCAGATCGGCGAATGCTACATGCAGTACAGTGGAG**GGTCCTTTTCATCACGTGCTATAAAAATAATTATAATTTAAATTTTTTAATATAAATATA |
| **CEN6_ARS4_rv** | **CAACGCATGAGGATGATGACAGCAGCACTCGTACCAGATAGAGACAGCTCTTCCGAACAT**GGACGGATCGCTTGCCTGTAAC |

#### Table S9 Additional primers used for fragments of assembly of IMF2

SHRs are marked in bold.

| Primer Name | Sequence 5' - 3' |
|---|---|
| Chunk_1-AH_rv | **CTCAGCCTTAGCCAATATGATCATGTCGTTGCGTCTCGGACCATCTAGTCTACTCTGAAG**TAATCTCGACAGCCTCTTCC |
| prHIS3_fw | **TCATTGCTGGCAGTGTAGCTGATAGCATCACATGACCTGCAGATGGCACTCACGTCACTG**TGCGGCATCAGAGCAGATTG |
| Chunk_4-AL_rv | **GCGTCCTCTGTATTAAGATGTCATGGTGTTGAGTCTGCACATGCAATGGCAATGAGTCAG**TGGTACGGTTTACTCCTTCACCTGTG |
| Chunk_7-AC_rv | **GCTACATCTTCCGTACTATGCTGTAGTCTCATGGTCGAGTTCTATTGCTGTTCGGCGGCA**CAGACATGGCGTAACCCCGTG |

#### Table S10 Additional primers used for amplification of fragments of Syn12

SHRs are marked in bold.

| Primer Name | Sequence 5' - 3' |
|---|---|
| ARS1-AD_fw | **CCGATAGCAGAATGTCACACATCATGCAGATCGGCGAATGCTACATGCAGTACAGTGGAG**GGCCTTTTGAAAAGCAAGCATAAAAGATCTAAAC |
| Chunk_18A_fw | **TAAGATGTGTGAACTGCGTCATACTCGCGTCAGGGATGCTTTCAGGCGTGCATGATGTAT** |
| Chunk_18A_rv | **GAGATGACTGGGTCCACTCTTTCGTGTATTTCGAGAGAGCGATACGCATGTCTCCATCGT**GCTAACTGTCACCCAACATAC |
| Chunk_18B_fw | **ACGATGGAGACATGCGTATCGCTCTCTCGAAATACACGAAAGAGTGGACCCAGTCATCTC**GATCCGCAAGTTCTTCATCG |
| Chunk_18B_rv | **CAGATCAGTGTCAATGAAGGTAGGCTGCTTGGCAATGCTTCTGGTGACTGGTAGATCATC**GCGATGTGCAATGTTCTTTGTTAC |
| Chunk_18C_fw | **GATGATCTACCAGTCACCAGAAGCATTGCCAAGCAGCCTACCTTCATTGACACTGATCTG**TGACAGTTCAATCAGCATCAG |
| Chunk_18C_rv | **AGCTGGCGTCGCGCATAAATGCATGATCTGTCCTGGCTGACACGCATCTGCACTAATGAT**ACTCTTGCCTGAGGAATAGC |
| Chunk_18D_fw | **ATCATTAGTGCAGATGCGTGTCAGCCAGGACAGATCATGCATTTATGCGCGACGCCAGCT**GACTCTGCTGATTATGCTTTAGTC |
| Chunk_18D_rv | **GCAGTAGCTTCCAGTACCTTCTCATACGTTATCACACTGGATATGCCATCGCGTCGAGGA**CACACCACGCCACTAAGCAG |
| Chunk_15A_fw | **TCCTCGACGCGATGGCATATCCAGTGTGATAACGTATGAGAAGGTACTGGAAGCTACTGC**TGCGAGCTGAATGCCATGAC |
| Chunk_15A_rv | **GATGAACGTGCCTTCGATTTATAGAAACTGCGCTGCCCTGTGATGAATTGTCTTAGCGCG**AAAGCGGCAGGTTGAGGTCC |
| Chunk_15B_fw | **CGCGCTAAGACAATTCATCACAGGGCAGCGCAGTTTCTATAAATCGAAGGCACGTTCATC**CTGCGACCACGCAGTTTGAG |
| Chunk_15B_rv | **CGTGCCGGTTAATGAGCTATGCGTGTCATGTATCCTTAGGCATATCCTTAACACGCAGTG**CGCCTTTGGCATGATCGAACAG |
| Chunk_15C_fw | **CACTGCGTGTTAAGGATATGCCTAAGGATACATGACACGCATAGCTCATTAACCGGCACG**AACCGGCAGGTTATAGCTGATG |
| Chunk_15C_rv | **CGGGTCATTAGAGATAGTCTCTCAGGATTCAACTAGATGGTGATCTATTGTCTACGCGGC**ATGGCCCATATACACTTCGAGCAC |
| Chunk_15D_fw | **GCCGCGTAGACAATAGATCACCATCTAGTTGAATCCTGAGAGACTATCTCTAATGACCCG**ATGCGTGAATGGCTGGCAGAG |
| Chunk_15D_rv | **CGCTGACCTGTCTAACGTATCAACAGAATGCACGTCAGTCGTATGCTTGACGTGTCTGCC**CAGTATCAACCACCGGGTAAC |
| CEN6_ARS4_DJ_fw | **GGCAGACACGTCAAGCATACGACTGACGTGCATTCTGTTGATACGTTAGACAGGTCAGCG**GGTCCTTTTCATCACGTGCTATAAAAATAATTATAATTTAAATTTTTTAATATAAATATA |
| Chunk_16A_fw | **ATGTTCGGAAGAGCTGTCTCTATCTGGTACGAGTGCTGCTGTCATCATCCTCATGCGTTG**TGATGCGCGATGCTTATCAGG |
| Chunk_16A_rv | **CGCCGCTCTTAGAAGGCTATACGAGCTATGAGAGAGACTCGCTATCCATTCCGCTGAGTT**CGTCAGCGATGAGACGTTAC |
| Chunk_16B_fw | **AACTCAGCGGAATGGATAGCGAGTCTCTCTCATAGCTCGTATAGCCTTCTAAGAGCGGCG**GCATCGGTGAACAGGGTGCTAAG |
| Chunk_16B_rv | **CGCAAATGTCCCATCGTATTTCAGAACCTTGTCACTCATGCGAGCAAGTGTGACAGCTATA**TGGCGTTCTCCGCCCAGTATG |
| Chunk_16C_fw | **ATAGCTGTCACACTTGCTCGCATGAGTGACAAGGTTCTGAAATACGATGGGACATTTGCG**CCGGCGCAGATCACTTTCATAG |
| Chunk_16C_rv | **CGACAAAGTGCTCGTCACGTGCGTCAGCTTCGAGGCATATCAAGCACCTGCCGGATGATT**ATGCCCGTGAATGGCAAAGCG |
| Chunk_16D_fw | **AATCATCCGGCAGGTGCTTGATATGCCTCGAAGCTGACGCACGTGACGAGCACTTTGTCG**AACACCGGACGGCCTTTGCTAC |

| | |
|---|---|
| **Chunk_16D_rv** | **CCTCCGCTGCGTAGAGTAATCCTGGCTCTCGCGTGTATATTGATAGATTGTCTGTCAGGC**TCGCGCAGGTATGGTTCAGG |
| **Chunk 17A_fw** | **GCCTGACAGACAATCTATCAATATACACGCGAGAGCCAGGATTACTCTACGCAGCGGAGG**TACGCAGTTTATCGGCCAGTTG |
| **Chunk 17A_rv** | **CAACCACCTGACTAGAGTGTCAAAGCGTGCTCCTACATAGGTAGAGTTGCATAATCTGGC**AAATCGCTGAAGCGTTCC |
| **Chunk 17B_fw** | **GCCAGATTATGCAACTCTACCTATGTAGGAGCACGCTTTGACACTCTAGTCAGGTGGTTG**TTGATAATCGCGGATGGACG |
| **Chunk 17B_rv** | **ATTCAGCGGGTGATCCGACTTGACTACATTTAGGTGTGGCCTCCTTACTACTCTGAGATG**CATCCGGTGAAAGCGTACCC |
| **Chunk 19A_fw** | **CATCTCAGAGTAGTAAGGAGGCCACACCTAAATGTAGTCAAGTCGGATCACCCGCTGAAT**GCGATGGTCATTATTTACGGTAG |
| **Chunk 19A_rv** | **ATGCCGGTGGCCGAATCTATGGTCCACATTATTTGCTGCACAAGATAGTGCAGTAGCGTT**CCATTATTGGCAGGATACTTTGAG |
| **Chunk 17D_fw** | **AACGCTACTGCACTATCTTGTGCAGCAAATAATGTGGACCATAGATTCGGCCACCGGCAT**TGGGTGTTTATGCCCGGACTAGC |
| **Chunk 17D_rv** | **ATCGACGGTCCTCGCAAGATCTCAATGTGCAGTGGTATGCTGATAACTTGTGCCTGTGGC**GGGATTTAGATCCCACATTAACG |
| **ARS417_DR_fw** | **GCCACAGGCACAAGTTATCAGCATACCACTGCACATTGAGATCTTGCGAGGACCGTCGAT**ACAAGTCCTTAAGAATATACAAAAAGCTACATAAATATAA |
| **Turquoise_AL_rv** | **CTGACTCATTGCCATTGCATGTGCAGACTCAACACCATGACATCTTAATACAGAGGACGC**CGTCTCATTGGCAGCATAAA |
| **Turquoise_DS_fw** | **ATCAAGACTGAGGAGTACGTCAGGTTGCAGAGGATCACTTGTAATGAATGTGTGCTCGCT**GAACGTTGATAGGTCAAGATCAATG |
| **Telomerator_r_fw** | **AGCGAGCACACATTCATTACAAGTGATCCTCTGCAACCTGACGTACTCCTCAGTCTTGAT**CCGGGGGATCCGGTGATTG |
| **Telomerator_l_rv** | **TGACGCGCTTTCAACCATGTTGCTGGGATTTCAGATTAGCCACGTACCTGGCATCAATGA**CCAAAGCTGGAGCTCCACCG |

#### Table S11 Primers used to insert glycolysis in the SynCh1 (IMF6), SynCh2 (IMF2) and in the *can1* locus

SHRs are marked in bold.

| Primer Name | Sequence 5' - 3' |
|---|---|
| ARS418_fw + can1 | **GGTGTATGACTTATGAGGGTGAGAATGCGAAATGGCGTGGGAATGTGATTAAAGGTAATA**TGAAAGTTTATGTTTTTTCACTGGA |
| ARS418 fw_SHR AH | **CTTCAGAGTAGACTAGATGGTCCGAGACGCAACGACATGATCATATTGGCTAAGGCTGAG**GTGTACCGAAGACTGCATTGAAAG |
| ARS418 rv_tag G | **AAGGGCCATGACCACCTGATGCACCAATTAGGTAGGTCTGGCTATGTCTATACCTCTGGC**CATAGACACAGTACTTACATTTAATAAC |
| KanMX_fw+ G | **GCCAGAGGTATAGACATAGCCAGACCTACCTAATTGGTGCATCAGGTGGTCATGGCCCTT**GCGACATGGAGGCCCAGAATACC |
| KanMX_rv + AA | **ATAGCATAGGTGCAAGGCTCTCGCCGCTTGTCGAGCTATTGGCATGGATGTGCTCCCTAA**AGTATAGCGACCAGCATTCACATACG |
| FBA1_rv + AA | **TTAGGGAGCACATCCATGCCAATAGCTCGACAAGCGGCGAGAGCCTTGCACCTATGCTAT**AATGAGCTATCAAAAACGATAGATC |
| FBA1 Fw + H | **GTCACGGGTTCTCAGCAATTCGAGCTATTACCGATGATGGCTGAGGCGTTAGAGTAATCT**AACGTGAACAACAATACCAGCCTTC |
| TPI FW + H | **AGATTACTCTAACGCCTCAGCCATCATCGGTAATAGCTCGAATTGCTGAGAACCCGTGAC**AACGAAGACCCAGAGATGTTGTTGT |
| TPI Rv + P | **CTGATAGTGCTGTAAGTCGCCTCCATCTTAGCAGAGCTGTCCCTGAATGCGTACTCGTGA**TGAGTAACCCATATAGAGATCGTAC |
| PGK1 Rv + P | **TCACGAGTACGCATTCAGGGACAGCTCTGCTAAGATGGAGGCGACTTACAGCACTATCAG**AAATAATATCCTTCTCGAAAGCTTT |
| PGK1 Fw + Q | **GAGCTGAATGTATATGCTGCGGGATCATTGCACAGCTCTGAGAGCCCTGCAACGCGATAT**TCTTTTTATTAACCTTAATTTTTAT |
| ADH1 Rv + Q | **ATATCGCGTTGCAGGGCTCTCAGAGCTGTGCAATGATCCCGCAGCATATACATTCAGCTC**TTGTCCTCTGAGGACATAAAATACA |
| ADH1 Fw + N | **TTCTAGGCTTTGATGCAAGGTCCACATATCTTCGTTAGGACTCAATCGTGGCTGCTGATC**AACGAAGTCCAATGCTAGTAGAGAA |
| PYK1 Fw + N | **GATCAGCAGCCACGATTGAGTCCTAACGAAGATATGTGGACCTTGCATCAAAGCCTAGAA**AACGTGGTCAAACTTCAGAACTAAG |
| PYK1 Rv + O | **ATACTCCCTGCACAGATGAGTCAAGCTATTGAACACCGAGAACGCGCTGAACGATCATTC**ATAATCATGATAACCTTGAGGGAAG |
| TDH3 Rv + O | **GAATGATCGTTCAGCGCGTTCTCGGTGTTCAATAGCTTGACTCATCTGTGCAGGGAGTAT**ATCCTGGCGGAAAAAATTCATTTGT |
| TDH3 Fw + A | **GTGCCTATTGATGATCTGGCGGAATGTCTGCCGTGCCATAGCCATGCCTTCACATATAGT**ATACTAGCGTTGAATGTTAGCGTCA |
| ENO2 Fw + A | **ACTATATGTGAAGGCATGGCTATGGCACGGCAGACATTCCGCCAGATCATCAATAGGCAC**AACGGATGATGAAAACACTAAACGA |
| ENO2 Rv + B | **GTTGAACATTCTTAGGCTGGTCGAATCATTTAGACACGGGCATCGTCCTCTCGAAAGGTG**TAACGAAGACGTTACCAGCTGATTG |
| HXK2 Fw + B | **CACCTTTCGAGAGGACGATGCCCGTGTCTAAATGATTCGACCAGCCTAAGAATGTTCAAC**AACGACGCTGGTAAAGTACAGCTAC |
| HXK2 Rv + C | **CTAGCGTGTCCTCGCATAGTTCTTAGATTGTCGCTACGGCATATACGATCCGTGAGACGT**ACTTGAACAATAAATACGAAATCCT |
| PGI1 Rv + C | **ACGTCTCACGGATCGTATATGCCGTAGCGACAATCTAAGAACTATGCGAGGACACGCTAG**TTTTAAACAGTTGATGAGAACCTTT |
| PGI1 Fw + D | **AATCACTCTCCATACAGGGTTTCATACATTTCTCCACGGGACCCACAGTCGTAGATGCGT**AACGTATTCTTAGTGGATAACATGC |
| PFK1 Fw + D | **ACGCATCTACGACTGTGGGTCCCGTGGAGAAATGTATGAAACCCTGTATGGAGAGTGATT**AACGCGGCTAGTAAAAAAGAAAATT |
| PFK1 Rv + J | **CGACGAGATGCTCAGACTATGTGTTCTACCTGCTTGGACATCTTCGCGTATATGACGGCC**ATTCCATAGCTTAGTTTAATCAAGG |

| | |
|---|---|
| **PFK2 Rv + J** | **GGCCGTCATATACGCGAAGATGTCCAAGCAGGTAGAACACATAGTCTGAGCATCTCGTCG**AAATCGTCTATATCACATATTCCAG |
| **PFK2_fw + L** | **GCCGTAGCTTCCGCAAGTATGCCGTAGTTGAAGAGCATTTGCCGTCGGTTCAGGTCATAT**AACGATTCTCTGCTGCTTTGTTGC |
| **GPM1 Rv + L** | **ATATGACCTGAACCGACGGCAAATGCTCTTCAACTACGGCATACTTGCGGAAGCTACGGC**TATTGCTATAACATGTCATGTCACC |
| **GPM1 Fw + M** | **ACGAGAGATGAAGGCTCACCGATGGACTTAGTATGATGCCATGCTGGAAGCTCCGGTCAT**AACGGTGATACTTTGACAGGAGCTA |
| **PDC1 Fw + M** | **ATGACCGGAGCTTCCAGCATGGCATCATACTAAGTCCATCGGTGAGCCTTCATCTCTCGT**AACGCATGCGACTGGGTGAGCATAT |
| **PDC1 RV + AR** | **TGACGAGATTTGAGAAGTCCCCAATATCGACTCGTGATGTGCCATGCGTGCTGTCAGTAT**ACAGTGTTCCTTAATCAAGGATACC |
| **ARS1211_fw +AR** | **ATACTGACAGCACGCATGGCACATCACGAGTCGATATTGGGGACTTCTCAAATCTCGTCA**GACATAGTATTTCGCAACCTTTCAG |
| **ARS1211_rv + BU** | **AATCATGTGACCCAGGCTTGCGCATACATGATCCTTCTTGCGCTGCATGGGCGACTATAT**GACAGGCGTTTCTGTACCGCTGTTA |
| **ARS1211_rv + AI** | **CAGTGACGTGAGTGCCATCTGCAGGTCATGTGATGCTATCAGCTACACTGCCAGCAATGA**GACAGGCGTTTCTGTACCGCTGTTA |
| **ARS1211_rv + Can1** | **GATGAGAAAAGTAAAGAATTGTATCCATTGCGCTCTTTCCCGACGAGAGTAAATGGCGAG**GACAGGCGTTTCTGTACCGCTGTTA |

#### Table S12 Diagnostic primers used to check insertion of glycolysis in SynCh1 (IMF6), SynCh2 (IMF2) and in the *can1* locus

| Name | Sequence (5' - 3') |
|---|---|
| AE_dg_new_rv | TGCCGGACTATCGAAACGAG |
| dg_chunk1/2_ fw | AGAACTTGCTGGTGGAGATG |
| CAN1 PAGE rv | TGAAGGAGTTTCAAATGCTTCTAC |
| TPI1-forward | CATAGACGCAGTACTTACATTTAATAACAG |
| TPI1 Flank left RV | GAGAAGATGTTCTTATTCAAATTTCAACTG |
| FK186 | CTGCAGCGAGGAGCCGTAAT |
| c KanMX fw | GACATCATCTGCCCAGATGC |
| FBA1 Flank left rv | ATTTTACTCACGCTTGAAATTAACGGC |
| FBA1 Flank right FW | CGTATTACGATAATCCTGCTGTC |
| gntK_RV_seq4 | ACCGAGGTGGTATCCGAGAG |
| FK115 | GTGGCATGTGAGATTCTCC |
| pPGK1_fw 1 | GCTGCTACTGTTGCAAAGGC |
| YAT2_2 | CAGCTCTGGAACAACGACATCTG |
| Wan5543 Seq Primer RV 3 | TGTGGAAGCCCGATGTCTGG |
| I-forward | TGAGCCACTTAAATTTCGTGAATG |
| dg_PGK1_rv_tagP | CTATGTTCGGGTTCAGCGTA |
| AL_ctrl_rv | TGAGGGTAACATCAATTCAAGAAG |
| ADH1 Flank left RV | CGAGCAAATGCCTGCAAATCG |
| YAT2_2 | CAGCTCTGGAACAACGACATCTG |
| Wan5543 Seq Primer RV 3 | TGTGGAAGCCCGATGTCTGG |
| tPYK1_fwd | TCCAATTGTCGTCATAACGATGAGG |
| TDH3 Flank left RV | GGCGCTCTTAATAATTTTGGGG |
| Ptdh3 Ctrl Rv2 | GGGCATGTACGGGTTACAG |
| ENO2 Flank left RV | GGCGTGCGGTGTGTAGATGTATC |
| FK244 | CACCTTTCGAGAGGACGATG |
| B-reverse | ACGGAATAGAACACGATATTTGC |
| Probe HXK2 fw | CACAAGCCAGAAAGGGTTCC |
| C-reverse | CCCACGATGCTTCTACCAAC |
| qPCR HXK2-FW | TTGGTGCTAGAGCTGCTAGATTG |
| PGI1 Flank left RV | CACCTTGAACTTGCGAAAAAGGTTCTC |
| PGI1 Flank right FW | CTACCCACATCCGAACATTGC |
| D-reverse | AATCATGTTGATGACGACAATGG |
| FW_PFK1_seq2 | CAATTGCGTGACGGTAGAAC |
| Q-PFK2-FW | TTGGTGGTTTCGAAGCTTTTG |
| FG-forward | TGTCTTACCCTGGACGGTATC |
| tGPM1_fwd | TTTTCAGCCTGTCGTGGTAGC |
| GPM1_P_FW | TATTGTAATATGTGTGTTTGTTTGGATTATTAAG |
| pGPM1_rev | GCATCACTGCATGTGTTAACCG |
| PDC1_P_RV | TTTGATTGATTTGACTGTGTTATTTTGCG |
| tPDC1-fwd | GAATTGGCTTAAGTCTGGGTCC |
| tPDC1 FW | GCGATTTAATCTCTAATTATTAGTTAAAG |
| dg_ARS1211_rv | GACAGGCGTTTCTGTACC |
| fCan1_qPCR_fw | CTCTTTCCCGACGAGAG |
| FK074 | CGCTTTACTAGGGCTTTCTG |
| CAT2promDiagfw | GCCGGTCACAACCCTCTTTC |

#### Table S13 Additional primers needed for insertion of *E.coli* chunks in SynCh1 (IMF6) and SynCh2 (IMF2)

SHRs are marked in bold.

| Primer Name | Sequence 5’ - 3’ |
|---|---|
| **Chunk_15A_fw + AA** | **TTAGGGAGCACATCCATGCCAATAGCTCGACAAGCGGCGAGAGCCTTGCACCTATGCTAT**TGCGAGCTGAATGCCATGAC |
| **Chunk_16A_fw + DJ** | **GGCAGACACGTCAAGCATACGACTGACGTGCATTCTGTTGATACGTTAGACAGGTCAGCG**TGATGCGCGATGCTTATCAGG |
| **Chunk 20A_fw + DR** | **GCCACAGGCACAAGTTATCAGCATACCACTGCACATTGAGATCTTGCGAGGACCGTCGAT**CAGGTCAGGTTTATGGCCTTTC |
| **Chunk 20A_rv + AR** | **TGACGAGATTTGAGAAGTCCCCAATATCGACTCGTGATGTGCCATGCGTGCTGTCAGTAT**AGCGCATCAGGCAACTACG |

**Table S14 Diagnostic primers used to check insertion of E.coli chunks in SynCh1 (IMF6), SynCh2 (IMF2) and in the *can1* locus**

| Name | Sequence (5' - 3') |
|---|---|
| AE_dg_new_rv | TGCCGGACTATCGAAACGAG |
| dg_chunk1/2_ fw | AGAACTTGCTGGTGGAGATG |
| CAN1 PAGE rv | TGAAGGAGTTTCAAATGCTTCTAC |
| TPI1-forward | CATAGACGCAGTACTTACATTTAATAACAG |
| TPI1 Flank left RV | GAGAAGATGTTCTTATTCAAATTTCAACTG |
| FK186 | CTGCAGCGAGGAGCCGTAAT |
| c KanMX fw | GACATCATCTGCCCAGATGC |
| dg_15A_rv | GCATGATCCATCGCATCGTAG |
| dg_15A_fw | TCGATGCGATCCTCGAACC |
| dg_15B_rv | CATTTCAGCCAGCCCATTTCGTC |
| dg_15B_fw | AAAGACGACTGGGCGTCTG |
| dg_15C_rv | GCCCGCAAATACCATCCTG |
| dg_15C_fw | ACAAACCGTACCGGAAGG |
| dg_15D_rv | GAATGCGGCGGAATCATC |
| dg_15D_fw | CTCGCTAAGAAGCTGTTG |
| dg_16A_rv | TGGTCGCAGTAACCTTAGTG |
| dg_16A_fw | GCCGTGAGGAAACAGTTG |
| dg_16B_rv | AATGGTGATGCCCTATGG |
| dg_16B_fw | CCGCTTTGTAGTGTTCCC |
| dg_16C_rv | GCCTGGCGATGACTATTTCTC |
| dg_16C_fw | GACCTGGGCGATAGTTTC |
| dg_16D_rv | CTGTGCCTGCAAGATGAG |
| dg_16D_fw | GGAGAGGCTGGCTTCGTTATG |
| dg_17A_rv | CTTTCTGGCGCTGCCAATGAG |
| dg_17A_fw | CGGCAGTATTGGGATTGTAG |
| dg_17B_rv | CCAGGGTAGAAAGATGGGATTG |
| dg_17B_fw | TTGACCGATGCCGACCGGAATG |
| dg_19A_rv | CAAGCGCCAGCGAAGGTAACAC |
| dg_19A_fw | TTCCCGATGAGTAAGGAGGAC |
| dg_17D_rv | CTGGGCGCTGTACAATAAG |
| dg_17D_fw | CGGTGCACTGGCATCTTCATAATC |
| dg_20A_rv | CCCTGGCATAACCACGAATTTG |
| dg_20A_fw | AACGAGACGCCAGAGAAG |
| dg_20A_fw_new | GATCCCGGAAGAGTATTCG |
| FK074 | CGCTTTACTAGGGCTTTCTG |
| CAT2promDiagfw | GCCGGTCACAACCCTCTTTC |

#### Table S15 Primers used to repair *sga1* upon SinLoG removal

| Primer Name | Sequence 5' - 3' |
|---|---|
| **COUNTER SELECT oligo fw** | TTTTTCTCATCTCTTGGCTCTGGATCCGTTATCTGTTCTGTTACACAAGAAATCGTACATACTAGAGCAAGATTTCAAATAAGTAACAGCAGCCATACGTTGAAACTACGGCAAAGGATT |
| **COUNTER SELECT oligo rv** | AATCCTTTGCCGTAGTTTCAACGTATGGCTGCTGTTACTTATTTGAAATCTTGCTCTAGTATGTACGATTTCTTGTGTAACAGAACAGATAACGGATCCAGAGCCAAGAGATGAGAAAAA |
| **Diagnostic PCR primers:** | |
| **SGA1 outside fw** | CTTGGCTCTGGATCCGTTATCTG |
| **SGA1 outside rv** | TTGATGTAAATATCTAGGAAATACACTTG |

#### Table S16 Primers used for the *URA3* and *HIS3* expression cassette in the *X2* locus

Flanks for homologous recombination and SHRs are marked in bold.

| Primer Name | Sequence 5' - 3' |
|---|---|
| X2_URA3p_fw | **TCACAGAGGGATCCCGTTACCCATCTATGCTGAAGATTTATCATACTATTCCTCCGCTCG**TGAGTATTTTCAATAAATTTGTAGAGGACT |
| DT_ URA3t_rv | **ATACGAAATGAGGAGGCTCTACCGAAGCTCATCCATAATGCTCCATAATGCTGATGCAGG**TTGCTTTTGTTCCACTACTTTTTG |
| DT_HIS3p_fw | **CCTGCATCAGCATTATGGAGCATTATGGATGAGCTTCGGTAGAGCCTCCTCATTTCGTAT**TGCGGCATCAGAGCAGATTG |
| X2_HIS3t_rv | **GTCATAACTCAATTTGCCTATTTCTTACGGCTTCTCATAAAACGTCCCACACTATTCAGG**GCATCTGTGCGGTATTTCACAC |
| **Diagnostic PCR primers:** | |
| X-2 dg fw | TCCTCGGGCAGAGAAACTCG |
| X-2 dg rv | GTGAGCCTCTTACCTGTTTG |
| pH0 fw | GGGAATCTCGGTCGTAATG |
| HIS3_pROS_dg rv | TATACCTGTGTGGACGTTAA |

#### Table S17 Primers used for the fluorescent marker expression cassettes in the YPRcTau3 locus

Flanks for homologous recombination are marked in bold.

| Primer Name | Sequence 5' - 3' |
|---|---|
| **YPRcTau3_pCCW12_fw** | **ACAGTTTTGACAACTGGTTACTTCCCTAAGACTGTTTATATTAGGATTGTCAAGACACTC**AACGCACCCATGAACCACAC |
| **YPRcTau3_tENO1_rv** | **ATAATTATAATATCCTGGACACTTTACTTATCTAGCGTATGTTATTACTCGATAAGTGCT**GGCAGCATACATGGGTGACCAAA |
| **YPRcTau3_pTEF1_fwd** | **ACAGTTTTGACAACTGGTTACTTCCCTAAGACTGTTTATATTAGGATTGTCAAGACACTC**CCTTGCCAACAGGGAGTTC |
| **YPRcTau3_tTDH1_rev** | **ATAATTATAATATCCTGGACACTTTACTTATCTAGCGTATGTTATTACTCGATAAGTGCT**CGTTCAGGGTAATATATTTTAACC |
| **YPRcTau3_pTEF2_fw** | **ACAGTTTTGACAACTGGTTACTTCCCTAAGACTGTTTATATTAGGATTGTCAAGACACTC**GAACGTTGATAGGTCAAGATCAATG |
| **YPRcTau3_tSSA1_rv** | **ATAATTATAATATCCTGGACACTTTACTTATCTAGCGTATGTTATTACTCGATAAGTGCT**CGTCTCATTGGCAGCATAAA |
| **Diagnostic PCR primers:** | |
| **YPRCtau3 dg FW** | AATACGAGGCGAATGTCTAGG |
| **YPRCtau3 dg RV** | GCCTCCCCTAGCTGAACAAC |

#### Table S18 Primers used on the original purified *E.coli* fragments used for assembly of the SynCh to elucidate template origin.

Primers used to check if mutations observed with WGS are already present in template DNA. Template DNA is a mix of two *E.coli* strains: *E.coli* BL21 and *E.coli* XL1-Blue.

| Primer Name | Sequence 5' - 3' | Target fragment |
|---|---|---|
| Chunk 1_fw | TTGACGCCACCATCAAAGAG | Chunk 1B |
| Chunk_2D_fw | AGACTGATACCATCGCTACGAGATTTCGCCAATGCCGCTAGGACCATCTGAATCATGCGCGATGGCGCAAATTCTTCAATGGATCG | Chunk 2D |
| Chunk2_fw | TCCGTATGCGCGGTAAAG | Chunk 2C |
| Chunk_15C_rv | CGGGTCATTAGAGATAGTCTCTCAGGATTCAACTAGATGGTGATCTATTGTCTACGCGGCATGGCCCATATACACTTCGAGCAC | Chunk 15C |
| AU_FW | GATACGCATATCACTAGACACG | Chunk 1D |
| Chunk 6 short_fw | TAAGATGTGTGAACTGCGTCATACTC | Chunk 6A |

**Table S19 Plasmids**

| Plasmid | Relevant characteristics | Source |
|---|---|---|
| **pYTK002** | camR ConLS connector | Lee *et al*. (6) |
| **pYTK010** | camR *pCCW12* | Lee *et al*. (6) |
| **pYTK013** | camR *pTEF1* | Lee *et al*. (6) |
| **pYTK014** | camR *pTEF2* | Lee *et al*. (6) |
| **pYTK032** | camR *mTurquoise2* | Lee *et al*. (6) |
| **pYTK033** | camR *Venus* | Lee *et al*. (6) |
| **pYTK034** | camR *mRuby2* | Lee *et al*. (6) |
| **pYTK047** | camR GFP dropout | Lee *et al*. (6) |
| **pYTK051** | camR *tENO1* | Lee *et al*. (6) |
| **pYTK052** | camR *tSSA1* | Lee *et al*. (6) |
| **pYTK056** | camR *tTDH1* | Lee *et al*. (6) |
| **pYTK067** | camR ConR1 | Lee *et al*. (6) |
| **pYTK072** | camR ConRE | Lee *et al*. (6) |
| **pYTK074** | camR *URA3* | Lee *et al*. (6) |
| **pYTK081** | camR CEN6/ARS4 | Lee *et al*. (6) |
| **pYTK082** | camR 2 µm | Lee *et al*. (6) |
| **pYTK083** | camR ampR-ColE1 | Lee *et al*. (6) |
| **pUD538** | CEN6/ARS4, ampR, *URA3, sfGFP* | This study |
| **pUDC191** | CEN6/ARS4, ampR, *URA3, pCCW12-mRuby2-tENO1* | This study |
| **pUDC192** | CEN6/ARS4, ampR, *URA3, pTEF2-mTurquoise2-tSSA1* | This study |
| **pUDC193** | CEN6/ARS4, ampR, *URA3, pTEF1-Venus-tTDH1* | This study |
| **pGGkd017** | 2 µm, ampR *URA3, sfGFP* | This study |
| **pUDE767** | 2 µm, ampR, *URA3, pHXK2-ScHXK2-tHXK2* | This study |
| **pUDE768** | 2 µm, ampR, *URA3, pPGI1-ScPGI1-tPGI1* | This study |
| **pUDE769** | 2 µm, ampR, URA3, pPFK1-ScPFK1-tPFK1 | This study |
| **pUDE770** | 2 µm, ampR, *URA3, pPFK2-ScPFK2-tPFK2* | This study |
| **pUDE771** | 2 µm, ampR, *URA3, pFBA1-ScFBA1-tFBA1* | This study |
| **pUDE772** | 2 µm, ampR, *URA3, pTPI1-ScTPI1-tTPI1* | This study |
| **pUDE773** | 2 µm, ampR, *URA3, pTDH3-ScTDH3-tTDH3* | This study |
| **pUDE774** | 2 µm, ampR, *URA3, pPGK1-ScPGK1-tPGK1* | This study |
| **pUDE775** | 2 µm, ampR, URA3, *pGPM1-ScGPM1-tGPM1* | This study |
| **pUDE776** | 2 µm, ampR, *URA3, pENO2-ScENO2-tENO2* | This study |
| **pUDE777** | 2 µm, ampR, *URA3, pPYK1-ScPYK1-tPYK1* | This study |
| **pUDE778** | 2 µm, ampR, *URA3, pPDC1-ScPDC1-tPDC1* | This study |
| **pUDE779** | 2 µm, ampR, *URA3, pADH1-ScADH1-tADH1* | This study |
| **pUDE714** | 2 µm, ampR, *KanMX*, gRNA (cpf1)-*HIS4* | Swiat *et al*.(19) |
| **pLM092** | CEN6/ARS4, ampR, *HIS3*, 5'*URA3-ACT1*intron[Tess-I-SceI-Tess]-3'*URA3* | Mitchell *et al.*(20) |
| **pMEL10** | 2µm, ampR, *URA3*, gRNA-*CAN1*.Y | Mans *et al.* (7) |
| **pROS12** | 2 µm, ampR, *hphNT1*, gRNA-*CAN1*.Y gRNA-*ADE2*.Y | Mans *et al.* (7) |
| **pROS13** | 2 µm, ampR, *kanMX*, gRNA.*CAN1*.Y gRNA.*ADE2*.Y | Mans *et al.* (7) |
| **p426-SNR52p-gRNA.CAN1.Y-SUP4t** | 2 µm, ampR, *URA3*, gRNA.*CAN1*.Y | DiCarlo *et al.*(21) |
| **pUDR400** | 2 µm, ampR, *hphNT1*, gRNA.-*mTurquoise2* | This study |
| **pUDR514** | 2 µm, ampR, KanMX, gRNA-*YPRcTau3* gRNA-*YPRcTau3* | This study |
| **pUDR547** | 2µm ampR hphNT1 gRNA-*X2* gRNA-*X2* | This study |
| **pUDR557** | 2 µm, ampR, *hphNT1*, gRNA-*sgaI* gRNA-*sgaI* | This study |

#### Table S20 Template for PCR amplifications

| Fragment | Template |
|---|---|
| ***E. coli* chunks** | *E.coli* XL1-blue/*E.coli* BL21 genome |
| ***CEN6/ARS4*** | pLM092 |
| ***HIS3*** | pLM092 |
| **Telomerator** | pLM092 |
| ***mRuby2*** | pUDC191 |
| ***mTurquoise2*** | pUDC192 |
| ***Venus*** | pUDC193 |
| ***ARS417*** | Annealing of complementary primers |
| ***ARS1*** | Annealing of complementary primers |
| ***ARS418*** | *S. cerevisiae* CEN.PK113-7D genome |
| ***ARS1211*** | *S. cerevisiae* CEN.PK113-7D genome |
| ***KanMX*** | pUDE714 |
| ***FBA1*** | pUDE771 |
| ***TPI1*** | *S. cerevisiae* CEN.PK113-7D genome |
| ***PGK1*** | *S. cerevisiae* CEN.PK113-7D genome |
| ***ADH1*** | *S. cerevisiae* CEN.PK113-7D genome |
| ***PYK1*** | *S. cerevisiae* CEN.PK113-7D genome |
| ***TDH3*** | *S. cerevisiae* CEN.PK113-7D genome |
| ***ENO2*** | *S. cerevisiae* CEN.PK113-7D genome |
| ***HXK2*** | *S. cerevisiae* CEN.PK113-7D genome |
| ***PGI1*** | pUDE768 |
| ***PFK1*** | pUDE769 |
| ***PFK2*** | pUDE770 |
| ***GPM1*** | pUDE775 |
| ***PDC1*** | *S. cerevisiae* CEN.PK113-7D genome |

**Table S21 List of SynCh1 chromosome fragments (IMF1 and IMF6)**
* Size of the fragments does not include the SHR sequences.

| SHR 5’ | part | SHR 3’ | Size (bp)* | Primer Fw | Primer Rv |
|---|---|---|---|---|---|
| **AE** | Chunk 1A | AT | 2514 | Ecoli_ch1_fw | Chunk_1A_rv |
| **AT** | Chunk 1B | AS | 2424 | Chunk_1B_fw | Chunk_1B_rv |
| **AS** | Chunk 1C | AU | 2443 | Chunk_1C_fw | Chunk_1C_rv |
| **AU** | Chunk 1D | AF | 2596 | Chunk_1D_fw | Ecoli_ch1_rv |
| **AF** | Chunk 2A | AV | 2501 | Ecoli_ch2_fw | Chunk_2A_rv |
| **AV** | Chunk 2B | AW | 2506 | Chunk_2B_fw | Chunk_2B_rv |
| **AW** | Chunk 2C | AX | 2385 | Chunk_2C_fw | Chunk_2C_rv |
| **AX** | Chunk 2D | AG | 2491 | Chunk_2D_fw | Ecoli_ch2_rv |
| **AG** | Chunk 3A | AY | 2539 | Ecoli_ch3_fw | Chunk_3A_rv |
| **AY** | Chunk 3B | AZ | 2517 | Chunk_3B_fw | Chunk_3B_rv |
| **AZ** | Chunk 3C | BA | 2503 | Chunk_3C_fw | Chunk_3C_rv |
| **BA** | Chunk 3D | AH | 2406 | Chunk_3D_fw | Ecoli_ch3_rv |
| **AH** | *mTurquoise 2* | AI | 1683 | prTEF2_Turquoise2_tSSA1_fw | prTEF2_Turquoise2_tSSA1_rv |
| **AI** | *ARS417* | BU | 60 | ARS417_fw | ARS417_rv |
| **BU** | *HIS3* | AJ | 1250 | BU-His3_fw | terHIS3_rv |
| **AJ** | Chunk 4A | BC | 2526 | Ecoli_ch4_fw | Chunk_4A_rv |
| **BC** | Chunk 4B | BD | 2488 | Chunk_4B_fw | Chunk_4B_rv |
| **BD** | Chunk 4C | BE | 2470 | Chunk_4C_fw | Chunk_4C_rv |
| **BE** | Chunk 4D | AK | 2419 | Chunk_4D_fw | Ecoli_ch4_rv |
| **AK** | Chunk 9A | BF | 2405 | Ecoli_ch9_fw | Chunk_9.2A_rv |
| **BF** | Chunk 9B | BS | 2565 | Chunk_9.2B_fw | Chunk_9.2B_rv |
| **BS** | Chunk 9C | BT | 2487 | Chunk_9.2C_fw | Chunk_9.2C_rv |
| **BT** | Chunk 9D | AQ | 2502 | Chunk_9.2D_fw | Chunk_9.2D_rv |
| **AQ** | Chunk 5A | BG | 2485 | Ecoli_ch5_fw | Chunk_5A_rv |
| **BG** | Chunk 5B | BH | 2521 | Chunk_5B_fw | Chunk_5B_rv |
| **BH** | Chunk 5C | BI | 2513 | Chunk_5C_fw | Chunk_5C_rv |
| **BI** | Chunk 5D | AL | 2441 | Chunk_5D_fw | Ecoli_ch5_rv |
| **AL** | *Venus* | AM | 1666 | prTEF1_Venus_tENO2_fw | prTEF1_Venus_tTDH1_rv |
| **AM** | *Ars1* | AN | 56 | ARS1_fw | ARS1_rv |
| **AN** | Chunk 6A | BP | 2508 | Ecoli_ch6_fw | Chunk_6A_rv |
| **BP** | Chunk 6B | BQ | 2494 | Chunk_6B_fw | Chunk_6B_rv |
| **BQ** | Chunk 6C | BR | 2497 | Chunk_6C_fw | Chunk_6C_rv |
| **BR** | Chunk 6D | AO | 2414 | Chunk_6D_fw | Ecoli_ch6_rv |
| **AO 30bp** | Telomerator | AO 30bp | 1623 | Telomerator_pos7n_fw | Telomerator_pos7_rv |
| **AO** | Chunk 7A | BJ | 2472 | Ecoli_ch7_fw | Chunk_7A_rv |
| **BJ** | Chunk 7B | BK | 2435 | Chunk_7B_fw | Chunk_7B_rv |
| **BK** | Chunk 7C | BL | 2478 | Chunk_7C_fw | Chunk_7C_rv |
| **BL** | Chunk 7D | AP | 2526 | Chunk_7D_fw | Ecoli_ch7_rv |

| | | | | | |
|---|---|---|---|---|---|
| **AP** | Chunk 8A | BM | 2477 | Ecoli_ch8_fw | Chunk_8A_rv |
| **BM** | Chunk 8B | BN | 2567 | Chunk_8B_fw | Chunk_8B_rv |
| **BN** | Chunk 8C | BO | 2454 | Chunk_8C_fw | Chunk_8C_rv |
| **BO** | Chunk 8D | AC | 2452 | Chunk_8D_fw | Ecoli_ch8_rv |
| **AC** | *mRuby2* | AD | 1667 | prCCW12_mRuby_tENO1_fw | prCCW12_mRuby_tENO1_rv |
| **AD** | *CEN6/ARS4* | AE | 519 | CEN6_ARS4_fw | CEN6_ARS4_rv |

**Table S22 List of SynCh2 chromosome fragments (IMF2)**
* Size of the fragments does not include the SHR sequences.

| SHR 5’ | part | SHR 3’ | Size (bp)* | Primer Fw | Primer Rv |
|---|---|---|---|---|---|
| **AE** | Chunk 1A | AT | 2514 | Ecoli_ch1_fw | Chunk_1A_rv |
| **AT** | Chunk 1B | AS | 2422 | Chunk_1B_fw | Chunk_1B_rv |
| **AS** | Chunk 1C | AU | 2443 | Chunk_1C_fw | Chunk_1C_rv |
| **AU** | Chunk 1D | AH | 2622 | Chunk_1D_fw | Chunk_1-AH_rv |
| **AH** | *mTurquoise 2* | AI | 1683 | prTEF2_Turquoise2_tSSA1_fw | prTEF2_Turquoise2_tSSA1_rv |
| **AI** | *HIS3* | AJ | 1250 | prHIS3_fw | terHIS3_rv |
| **AJ** | Chunk 4A | BC | 2514 | Ecoli_ch4_fw | Chunk_4A_rv |
| **BC** | Chunk 4B | BD | 2529 | Chunk_4B_fw | Chunk_4B_rv |
| **BD** | Chunk 4C | BE | 2473 | Chunk_4C_fw | Chunk_4C_rv |
| **BE** | Chunk 4D | AL | 2419 | Chunk_4D_fw | Chunk_4-AL_rv |
| **AL** | *Venus* | AM | 1666 | prTEF1_Venus_tENO2_fw | prTEF1_Venus_tTDH1_rv |
| **AM** | *Ars1* | AN | 56 | ARS1_fw | ARS1_rv |
| **AN** | Chunk 6A | BP | 2508 | Ecoli_ch6_fw | Chunk_6A_rv |
| **BP** | Chunk 6B | BQ | 2494 | Chunk_6B_fw | Chunk_6B_rv |
| **BQ** | Chunk 6C | BR | 2497 | Chunk_6C_fw | Chunk_6C_rv |
| **BR** | Chunk 6D | AO | 2414 | Chunk_6D_fw | Ecoli_ch6_rv |
| **AO 30bp** | Telomerator | AO 30bp | 1683 | Telomerator_pos7n_fw | Telomerator_pos7_rv |
| **AO** | Chunk 7A | BJ | 2472 | Ecoli_ch7_fw | Chunk_7A_rv |
| **BJ** | Chunk 7B | BK | 2497 | Chunk_7B_fw | Chunk_7B_rv |
| **BK** | Chunk 7C | BL | 2472 | Chunk_7C_fw | Chunk_7C_rv |
| **BL** | Chunk 7D | AC | 2525 | Chunk_7D_fw | Chunk_7-AC_rv |
| **AC** | *mRuby2* | AD | 1667 | prCCW12_mRuby_tENO1_fw | prCCW12_mRuby_tENO1_rv |
| **AD** | *CEN6/ARS4* | AE | 519 | CEN6_ARS4_fw | CEN6_ARS4_rv |

**Table S23 List of parts for construction of Syn15 (IMF13, IMF17)**
* Size of the fragments does not include the SHR sequences.

| SHR 5’ | part | SHR 3’ | Size (bp)* | Primer Fw | Primer Rv |
|---|---|---|---|---|---|
| **AH** | ARS418 | G | 216 | ARS418 fw_SHR AH | ARS418_rv_tag G |
| **G** | KanMX | AA | 1358 | KanMX_fw + G | KanMX_rv + AA |
| **AA** | FBA1 ← | H | 2184 | FBA1 Rv + AA | FBA1 Fw + H |
| **H** | TPI1 → | P | 1851 | TPI FW + H | TPI Rv + P |
| **P** | PGK1 ← | Q | 2351 | PGK1 Rv + P | PGK1 Fw + Q |
| **Q** | ADH1 ← | N | 2151 | ADH1 Rv + Q | ADH1 Fw + N |
| **N** | PYK1 → | O | 2607 | PYK1 Fw + N | PYK1 Rv + O |
| **O** | TDH3 ← | A | 2099 | TDH3 Rv + O | TDH3 Fw + A |
| **A** | ENO2 → | B | 2418 | ENO2 Fw + A | ENO2 Rv + B |
| **B** | HXK2 → | C | 2565 | HXK2 Fw + B | HXK2 Rv + C |
| **C** | PGI1 ← | D | 2769 | PGI1 Rv + C | PGI1 Fw + D |
| **D** | PFK1 → | J | 4068 | PFK1 Fw + D | PFK1 Rv + J |
| **J** | PFK2 ← | L | 3984 | PFK2 Rv + J | PFK2 Fw + L |
| **L** | GPM1 ← | M | 1848 | GPM1 Rv + L | GPM1 Fw + M |
| **M** | PDC1 → | AR | 2796 | PDC1 Fw + M | PDC1 RV + AR |
| **AR** | ARS1211 | BU | 251 | ARS1211_fw +AR | ARS1211 Rv + BU |

**Table S24 List of parts for construction of Syn16 (IMF14 and IMF18)**
* Size of the fragments does not include the SHR sequences.

| SHR 5’ | part | SHR 3’ | Size (bp)* | Primer Fw | Primer Rv |
|---|---|---|---|---|---|
| **AH** | ARS418 | G | 216 | ARS418 fw_SHR AH | ARS418_rv_tag G |
| **G** | KanMX | AA | 1358 | KanMX_fw + G | KanMX_rv + AA |
| **AA** | FBA1 ← | H | 2184 | FBA1 Rv + AA | FBA1 Fw + H |
| **H** | TPI1 → | P | 1851 | TPI FW + H | TPI Rv + P |
| **P** | PGK1 ← | Q | 2351 | PGK1 Rv + P | PGK1 Fw + Q |
| **Q** | ADH1 ← | N | 2151 | ADH1 Rv + Q | ADH1 Fw + N |
| **N** | PYK1 → | O | 2607 | PYK1 Fw + N | PYK1 Rv + O |
| **O** | TDH3 ← | A | 2099 | TDH3 Rv + O | TDH3 Fw + A |
| **A** | ENO2 → | B | 2418 | ENO2 Fw + A | ENO2 Rv + B |
| **B** | HXK2 → | C | 2565 | HXK2 Fw + B | HXK2 Rv + C |
| **C** | PGI1 ← | D | 2769 | PGI1 Rv + C | PGI1 Fw + D |
| **D** | PFK1 → | J | 4068 | PFK1 Fw + D | PFK1 Rv + J |
| **J** | PFK2 ← | L | 3984 | PFK2 Rv + J | PFK2 Fw + L |
| **L** | GPM1 ← | M | 1848 | GPM1 Rv + L | GPM1 Fw + M |
| **M** | PDC1 → | AR | 2796 | PDC1 Fw + M | PDC1 RV + AR |
| **AR** | ARS1211 | AI | 251 | ARS1211_fw +AR | ARS1211 Rv + AI |

**Table S25 List of parts for construction of Syn19 (IMF11)**
* Size of the fragments does not include the SHR sequences.

| SHR 5' | part | SHR 3' | Size (bp)* | Primer Fw | Primer Rv |
|---|---|---|---|---|---|
| **AH** | ARS418 | G | 216 | ARS418 fw_SHR AH | ARS418_rv_tag G |
| **G** | *KanMX* | AA | 1358 | KanMX_fw + G | KanMX_rv + AA |
| **AA** | 15A | DF | 2515 | Chunk_15A_fw + AA | Chunk_15A_rv |
| **DF** | 15B | DH | 2504 | Chunk_15B_fw | Chunk_15B_rv |
| **DH** | 15C | DI | 2489 | Chunk_15C_fw | Chunk_15C_rv |
| **DI** | 15D | DJ | 2497 | Chunk_15D_fw | Chunk_15D_rv |
| **DJ** | 16A | DK | 2520 | Chunk_16A_fw + DJ | Chunk_16A_rv |
| **DK** | 16B | DL | 2470 | Chunk_16B_fw | Chunk_16B_rv |
| **DL** | 16C | DM | 2517 | Chunk_16C_fw | Chunk_16C_rv |
| **DM** | 16D | DN | 2520 | Chunk_16D_fw | Chunk_16D_rv |
| **DN** | 17A | DO | 2499 | Chunk 17A_fw | Chunk 17A_rv |
| **DO** | 17B | DP | 2498 | Chunk 17B_fw | Chunk 17B_rv |
| **DP** | 19A | DQ | 2509 | Chunk 19A_fw | Chunk 19A_rv |
| **DQ** | 17D | DR | 2543 | Chunk 17D_fw | Chunk 17D_rv |
| **DR** | 20A | AR | 3610 | Chunk 20A_fw + DR | Chunk 20A_rv + AR |
| **AR** | ARS1211 | BU | 251 | ARS1211_fw +AR | ARS1211 Rv + BU |

**Table S26 List of parts for construction of Syn20 (IMF12)**
* Size of the fragments does not include the SHR sequences.

| SHR 5’ | part | SHR 3’ | Size (bp)* | Primer Fw | Primer Rv |
|---|---|---|---|---|---|
| **AH** | ARS418 | G | 216 | ARS418 fw_SHR AH | ARS418_rv_tag G |
| **G** | *KanMX* | AA | 1358 | KanMX_fw + G | KanMX_rv + AA |
| **AA** | 15A | DF | 2515 | Chunk_15A_fw + AA | Chunk_15A_rv |
| **DF** | 15B | DH | 2504 | Chunk_15B_fw | Chunk_15B_rv |
| **DH** | 15C | DI | 2489 | Chunk_15C_fw | Chunk_15C_rv |
| **DI** | 15D | DJ | 2497 | Chunk_15D_fw | Chunk_15D_rv |
| **DJ** | 16A | DK | 2520 | Chunk_16A_fw + DJ | Chunk_16A_rv |
| **DK** | 16B | DL | 2470 | Chunk_16B_fw | Chunk_16B_rv |
| **DL** | 16C | DM | 2517 | Chunk_16C_fw | Chunk_16C_rv |
| **DM** | 16D | DN | 2520 | Chunk_16D_fw | Chunk_16D_rv |
| **DN** | 17A | DO | 2499 | Chunk 17A_fw | Chunk 17A_rv |
| **DO** | 17B | DP | 2498 | Chunk 17B_fw | Chunk 17B_rv |
| **DP** | 19A | DQ | 2509 | Chunk 19A_fw | Chunk 19A_rv |
| **DQ** | 17D | DR | 2543 | Chunk 17D_fw | Chunk 17D_rv |
| **DR** | 20A | AR | 3610 | Chunk 20A_fw + DR | Chunk 20A_rv + AR |
| **AR** | ARS1211 | AI | 251 | ARS1211_fw +AR | ARS1211 Rv + AI |

**Table S27 List of SynCh3.1 parts**
* Size of the fragments does not include the SHR sequences.

| SHR 5' | part | SHR 3' | Size (bp)* | Primer Fw | Primer Rv |
|---|---|---|---|---|---|
| **AE** | Chunk 1A | AT | 2514 | Ecoli_ch1_fw | Chunk_1A_rv |
| **AT** | Chunk 1B | AS | 2424 | Chunk_1B_fw | Chunk_1B_rv |
| **AS** | Chunk 1C | AU | 2443 | Chunk_1C_fw | Chunk_1C_rv |
| **AU** | Chunk 1D | AF | 2596 | Chunk_1D_fw | Ecoli_ch1_rv |
| **AF** | Chunk 2A | AV | 2501 | Ecoli_ch2_fw | Chunk_2A_rv |
| **AV** | Chunk 2B | AW | 2506 | Chunk_2B_fw | Chunk_2B_rv |
| **AW** | Chunk 2C | AX | 2385 | Chunk_2C_fw | Chunk_2C_rv |
| **AX** | Chunk 2D | AG | 2491 | Chunk_2D_fw | Ecoli_ch2_rv |
| **AG** | Chunk 3A | AY | 2539 | Ecoli_ch3_fw | Chunk_3A_rv |
| **AY** | Chunk 3B | AZ | 2517 | Chunk_3B_fw | Chunk_3B_rv |
| **AZ** | Chunk 3C | BA | 2503 | Chunk_3C_fw | Chunk_3C_rv |
| **BA** | Chunk 3D | AH | 2406 | Chunk_3D_fw | Ecoli_ch3_rv |
| **AH** | *mTurquoise 2* | AI | 1683 | prTEF2_Turquoise2_tSSA1_fw | prTEF2_Turquoise2_tSSA1_rv |
| **AI** | *ARS417* | BU | 60 | ARS417_fw | ARS417_rv |
| **BU** | *HIS3* | AJ | 1250 | BU-His3_fw | terHIS3_rv |
| **AJ** | Chunk 4A | BC | 2526 | Ecoli_ch4_fw | Chunk_4A_rv |
| **BC** | Chunk 4B | BD | 2488 | Chunk_4B_fw | Chunk_4B_rv |
| **BD** | Chunk 4C | BE | 2470 | Chunk_4C_fw | Chunk_4C_rv |
| **BE** | Chunk 4D | AK | 2419 | Chunk_4D_fw | Ecoli_ch4_rv |
| **AK** | Chunk 9A | BF | 2405 | Ecoli_ch9_fw | Chunk_9.2A_rv |
| **BF** | Chunk 9B | BS | 2565 | Chunk_9.2B_fw | Chunk_9.2B_rv |
| **BS** | Chunk 9C | BT | 2487 | Chunk_9.2C_fw | Chunk_9.2C_rv |
| **BT** | Chunk 9D | AQ | 2502 | Chunk_9.2D_fw | Chunk_9.2D_rv |
| **AQ** | Chunk 5A | BG | 2485 | Ecoli_ch5_fw | Chunk_5A_rv |
| **BG** | Chunk 5B | BH | 2521 | Chunk_5B_fw | Chunk_5B_rv |
| **BH** | Chunk 5C | BI | 2513 | Chunk_5C_fw | Chunk_5C_rv |
| **BI** | Chunk 5D | AL | 2441 | Chunk_5D_fw | Ecoli_ch5_rv |
| **AL** | *Venus* | AM | 1666 | prTEF1_Venus_tENO2_fw | prTEF1_Venus_tTDH1_rv |
| **AM** | *Ars1* | AN | 56 | ARS1_fw | ARS1_rv |
| **AN** | Chunk 6A | BP | 2508 | Ecoli_ch6_fw | Chunk_6A_rv |
| **BP** | Chunk 6B | BQ | 2494 | Chunk_6B_fw | Chunk_6B_rv |
| **BQ** | Chunk 6C | BR | 2497 | Chunk_6C_fw | Chunk_6C_rv |
| **BR** | Chunk 6D | AO | 2414 | Chunk_6D_fw | Ecoli_ch6_rv |
| **AO** | Chunk 7A | BJ | 2472 | Ecoli_ch7_fw | Chunk_7A_rv |
| **BJ** | Chunk 7B | BK | 2435 | Chunk_7B_fw | Chunk_7B_rv |
| **BK** | Chunk 7C | BL | 2478 | Chunk_7C_fw | Chunk_7C_rv |
| **BL** | Chunk 7D | AP | 2526 | Chunk_7D_fw | Ecoli_ch7_rv |
| **AP** | Chunk 8A | BM | 2477 | Ecoli_ch8_fw | Chunk_8A_rv |

| | | | | | |
|---|---|---|---|---|---|
| **BM** | Chunk 8B | BN | 2567 | Chunk_8B_fw | Chunk_8B_rv |
| **BN** | Chunk 8C | BO | 2454 | Chunk_8C_fw | Chunk_8C_rv |
| **BO** | Chunk 8D | AC | 2452 | Chunk_8D_fw | Ecoli_ch8_rv |
| **AC** | *mRuby2* | AD | 1667 | prCCW12_mRuby_tENO1_fw | prCCW12_mRuby_tENO1_rv |
| **AD** | *CEN6/ARS4* | AE | 519 | CEN6_ARS4_fw | CEN6_ARS4_rv |

#### Table S28 List of SynCh3.2 parts

* Size of the fragments does not include the SHR sequences.

| SHR 5’ | part | SHR 3’ | Size (bp)* | Primer Fw | Primer Rv |
|---|---|---|---|---|---|
| **AE** | 1AB | AS | 5116 | Ecoli_ch1_fw | Chunk_1B_rv |
| **AS** | 1CD | AF | 5245 | Chunk_1C_fw | Ecoli_ch1_rv |
| **AF** | 2AB | AW | 5187 | Ecoli_ch2_fw | Chunk_2B_rv |
| **AW** | 2CD | AG | 5056 | Chunk_2C_fw | Ecoli_ch2_rv |
| **AG** | 3AB | AZ | 5236 | Ecoli_ch3_fw | Chunk_3B_rv |
| **AZ** | 3CD | AH | 5086 | Chunk_3C_fw | Ecoli_ch3_rv |
| **AH** | *mTurquoise2 - HIS3* | AJ | 3113 | prTEF2_Turquoise2_tSSA1_fw | terHIS3_rv |
| **AJ** | 4AB | BD | 5223 | Ecoli_ch4_fw | Chunk_4B_rv |
| **BD** | 4CD | AK | 5072 | Chunk_4C_fw | Ecoli_ch4_rv |
| **AK** | 9AB | BS | 5150 | Ecoli_ch9_fw | Chunk_9.2B_rv |
| **BS** | 9CD | AQ | 5169 | Chunk_9.2C_fw | Chunk_9.2D_rv |
| **AQ** | 5AB | BH | 5186 | Ecoli_ch5_fw | Chunk_5B_rv |
| **BH** | 5CD | AL | 5134 | Chunk_5C_fw | Ecoli_ch5_rv |
| **AL** | *Venus - ARS1* | AN | 1902 | prTEF1_Venus_tENO2_fw | ARS1_rv |
| **AN** | 6AB | BQ | 5182 | Ecoli_ch6_fw | Chunk_6B_rv |
| **BQ** | 6CD | AO | 5091 | Chunk_6C_fw | Ecoli_ch6_rv |
| **AO** | 7AB | BK | 5149 | Ecoli_ch7_fw | Chunk_7B_rv |
| **BK** | 7CD | AP | 5177 | Chunk_7C_fw | Ecoli_ch7_rv |
| **AP** | 8AB | BN | 5224 | Ecoli_ch8_fw | Chunk_8B_rv |
| **BN** | 8CD | AC | 5086 | Chunk_8C_fw | Ecoli_ch8_rv |
| **AC** | *mRuby2* | AD | 1787 | prCCW12_mRuby_tENO1_fw | prCCW12_mRuby_tENO1_rv |
| **AD** | *CEN6/ARS4* | AE | 639 | CEN6_ARS4_fw | CEN6_ARS4_rv |

#### Table S29 List of SynCh3.3 parts

* Size of the fragments does not include the SHR sequences.

| SHR 5’ | part | SHR 3’ | Size (bp)* | Primer Fw | Primer Rv |
|---|---|---|---|---|---|
| **AE** | 1AD | AF | 10301 | Ecoli_ch1_fw | Ecoli_ch1_rv |
| **AF** | 2AD | AG | 10183 | Ecoli_ch2_fw | Ecoli_ch2_rv |
| **AG** | 3AD | AH | 10265 | Ecoli_ch3_fw | Ecoli_ch3_rv |
| **AH** | *mTurquoise2* - 4B | BD | 8396 | prTEF2_Turquoise2_tSSA1_fw | Chunk_4B_rv |
| **BD** | 4C - 9B | BS | 10159 | Chunk_4C_fw | Chunk_9.2B_rv |
| **BS** | 9C - 5B | BH | 10295 | Chunk_9.2C_fw | Chunk_5B_rv |
| **BH** | 5C - *ARS1* | AN | 6976 | Chunk_5C_fw | ARS1_rv |
| **AN** | 6AD | AO | 10213 | Ecoli_ch6_fw | Ecoli_ch6_rv |
| **AO** | 7AD | AP | 10211 | Ecoli_ch7_fw | Ecoli_ch7_rv |
| **AP** | 8A – *mRuby2* | AD | 11977 | Ecoli_ch8_fw | prCCW12_mRuby_tENO1_rv |
| **AD** | *CEN6/ARS4* | AE | 639 | CEN6_ARS4_fw | CEN6_ARS4_rv |

**Table S30 List of SynCh4.1 parts**
* Size of the fragments does not include the SHR sequences.

| SHR 5’ | part | SHR 3’ | Size (bp)* | Primer Fw | Primer Rv |
|---|---|---|---|---|---|
| **AE** | Chunk 1A | AT | 2514 | Ecoli_ch1_fw | Chunk_1A_rv |
| **AT** | Chunk 1B | AS | 2422 | Chunk_1B_fw | Chunk_1B_rv |
| **AS** | Chunk 1C | AU | 2443 | Chunk_1C_fw | Chunk_1C_rv |
| **AU** | Chunk 1D | AH | 2622 | Chunk_1D_fw | Chunk_1-AH_rv |
| **AH** | *mTurquoise 2* | AI | 1683 | prTEF2_Turquoise2_tSSA1_fw | prTEF2_Turquoise2_tSSA1_rv |
| **AI** | *HIS3* | AJ | 1250 | prHIS3_fw | terHIS3_rv |
| **AJ** | Chunk 4A | BC | 2514 | Ecoli_ch4_fw | Chunk_4A_rv |
| **BC** | Chunk 4B | BD | 2529 | Chunk_4B_fw | Chunk_4B_rv |
| **BD** | Chunk 4C | BE | 2473 | Chunk_4C_fw | Chunk_4C_rv |
| **BE** | Chunk 4D | AL | 2419 | Chunk_4D_fw | Chunk_4-AL_rv |
| **AL** | *Venus* | AM | 1666 | prTEF1_Venus_tENO2_fw | prTEF1_Venus_tTDH1_rv |
| **AM** | *Ars1* | AN | 56 | ARS1_fw | ARS1_rv |
| **AN** | Chunk 6A | BP | 2508 | Ecoli_ch6_fw | Chunk_6A_rv |
| **BP** | Chunk 6B | BQ | 2494 | Chunk_6B_fw | Chunk_6B_rv |
| **BQ** | Chunk 6C | BR | 2497 | Chunk_6C_fw | Chunk_6C_rv |
| **BR** | Chunk 6D | AO | 2414 | Chunk_6D_fw | Ecoli_ch6_rv |
| **AO** | Chunk 7A | BJ | 2472 | Ecoli_ch7_fw | Chunk_7A_rv |
| **BJ** | Chunk 7B | BK | 2497 | Chunk_7B_fw | Chunk_7B_rv |
| **BK** | Chunk 7C | BL | 2472 | Chunk_7C_fw | Chunk_7C_rv |
| **BL** | Chunk 7D | AC | 2525 | Chunk_7D_fw | Chunk_7-AC_rv |
| **AC** | *mRuby2* | AD | 1667 | prCCW12_mRuby_tENO1_fw | prCCW12_mRuby_tENO1_rv |
| **AD** | *CEN6/ARS4* | AE | 519 | CEN6_ARS4_fw | CEN6_ARS4_rv |

**Table S31 List of SynCh4.2 parts**
* Size of the fragments does not include the SHR sequences.

| SHR Fw | part | SHR Rv | Size (bp)* | Primer Fw | Primer Rv |
|---|---|---|---|---|---|
| **AE** | 1AB | AS | 5116 | Ecoli_ch1_fw | Chunk_1B_rv |
| **AS** | 1CD | AH | 5245 | Chunk_1C_fw | Chunk_1-AH_rv |
| **AH** | *mTurquoise2 - HIS3* | AJ | 3113 | prTEF2_Turquoise2_tSSA1_fw | terHIS3_rv |
| **AJ** | 4AB | BD | 5223 | Ecoli_ch4_fw | Chunk_4B_rv |
| **BD** | 4CD | AL | 5072 | Chunk_4C_fw | Chunk_4-AL_rv |
| **AL** | *Venus - ARS1* | AN | 1902 | prTEF1_Venus_tENO2_fw | ARS1_rv |
| **AN** | 6AB | BQ | 5182 | Ecoli_ch6_fw | Chunk_6B_rv |
| **BQ** | 6CD | AO | 5091 | Chunk_6C_fw | Ecoli_ch6_rv |
| **AO** | 7AB | BK | 5149 | Ecoli_ch7_fw | Chunk_7B_rv |
| **BK** | 7C – *mRuby2* | AD | 6904 | Chunk_7C_fw | prCCW12_mRuby_tENO1_rv |
| **AD** | *CEN6/ARS4* | AE | 639 | CEN6_ARS4_fw | CEN6_ARS4_rv |

#### Table S32 List of SynCh4.3 parts

* Size of the fragments does not include the SHR sequences.

| SHR 5' | part | SHR 3' | Size (bp)* | Primer Fw | Primer Rv |
|---|---|---|---|---|---|
| **AE** | 1AD | AH | 10301 | Ecoli_ch1_fw | Chunk_1-AH_rv |
| **AH** | *mTurquoise2* - 4B | BD | 8276 | prTEF2_Turquoise2_tSSA1_fw | Chunk_4B_rv |
| **BD** | 4C - ARS1 | AN | 6914 | Chunk_4C_fw | ARS1_rv |
| **AN** | 6AD | AO | 10213 | Ecoli_ch6_fw | Ecoli_ch6_rv |
| **AO** | 7A – *mRuby2* | AD | 11993 | Ecoli_ch7_fw | prCCW12_mRuby_tENO1_rv |
| **AD** | *CEN6/ARS4* | AE | 639 | CEN6_ARS4_fw | CEN6_ARS4_rv |

**Table S33 List of Syn12 chromosome parts (IMF23, IMF24, IMF25, IMF26)**
* Size of the fragments does not include the SHR sequences.

| SHR 5' | part | SHR 3' | Size (bp)* | Primer Fw | Primer Rv |
|---|---|---|---|---|---|
| **AO** | Chunk 7A | BJ | 2472 | Ecoli_ch7_fw | Chunk_7A_rv |
| **BJ** | Chunk 7B | BK | 2435 | Chunk_7B_fw | Chunk_7B_rv |
| **BK** | Chunk 7C | BL | 2478 | Chunk_7C_fw | Chunk_7C_rv |
| **BL** | Chunk 7D | AP | 2526 | Chunk_7D_fw | Ecoli_ch7_rv |
| **AP** | Chunk 8A | BM | 2477 | Ecoli_ch8_fw | Chunk_8A_rv |
| **BM** | Chunk 8B | BN | 2567 | Chunk_8B_fw | Chunk_8B_rv |
| **BN** | Chunk 8C | BO | 2454 | Chunk_8C_fw | Chunk_8C_rv |
| **BO** | Chunk 8D | AC | 2452 | Chunk_8D_fw | Ecoli_ch8_rv |
| **AC** | *mRuby2* | AD | 1667 | prCCW12_mRuby_tENO1_fw | prCCW12_mRuby_tENO1_rv |
| **AD** | *ARS1* | AN | 56 | ARS1-AD_fw | ARS1_rv |
| **AN** | Chunk 18A | BP | 2525 | Chunk_18A_fw | Chunk_18A_rv |
| **BP** | Chunk 18B | BQ | 2510 | Chunk_18B_fw | Chunk_18B_rv |
| **BQ** | Chunk 19C | BR | 2495 | Chunk 19C_fw | Chunk 19C_rv |
| **BR** | Chunk 19D | DE | 2496 | Chunk 19D_fw | Chunk 19D_rv |
| **DE** | Chunk 15A | DF | 2515 | Chunk_15A_fw | Chunk_15A_rv |
| **DF** | Chunk 15B | DH | 2504 | Chunk_15B_fw | Chunk_15B_rv |
| **DH** | Chunk 15C | DI | 2489 | Chunk_15C_fw | Chunk_15C_rv |
| **DI** | Chunk 15D | DJ | 2497 | Chunk_15D_fw | Chunk_15D_rv |
| **DJ** | *CEN6/ARS4* | AE | 519 | CEN6_ARS4_DJ_fw | CEN6_ARS4_rv |
| **AE** | Chunk 16A | DK | 2520 | Chunk_16A_fw | Chunk_16A_rv |
| **DK** | Chunk 16B | DL | 2470 | Chunk_16B_fw | Chunk_16B_rv |
| **DL** | Chunk 16C | DM | 2517 | Chunk_16C_fw | Chunk_16C_rv |
| **DM** | Chunk 16D | DN | 2520 | Chunk_16D_fw | Chunk_16D_rv |
| **DN** | Chunk 17A | DO | 2499 | Chunk 17A_fw | Chunk 17A_rv |
| **DO** | Chunk 17B | DP | 2498 | Chunk 17B_fw | Chunk 17B_rv |
| **DP** | Chunk 19A | DQ | 2509 | Chunk 19A_fw | Chunk 19A_rv |
| **DQ** | Chunk 17D | DR | 2543 | Chunk 17D_fw | Chunk 17D_rv |
| **DR** | *ARS417* | BU | 60 | ARS417_DR_fw | ARS417_rv |
| **BU** | *HIS3* | AJ | 1250 | BU-His3_fw | terHIS3_rv |
| **AJ** | Chunk 4A | BC | 2526 | Ecoli_ch4_fw | Chunk_4A_rv |
| **BC** | Chunk 4B | BD | 2488 | Chunk_4B_fw | Chunk_4B_rv |
| **BD** | Chunk 4C | BE | 2470 | Chunk_4C_fw | Chunk_4C_rv |
| **BE** | Chunk 4D | AK | 2419 | Chunk_4D_fw | Ecoli_ch4_rv |
| **AK** | Chunk 9.2A | BF | 2405 | Ecoli_ch9_fw | Chunk_9.2A_rv |
| **BF** | Chunk 9.2B | BS | 2565 | Chunk_9.2B_fw | Chunk_9.2B_rv |
| **BS** | Chunk 9.2C | BT | 2487 | Chunk_9.2C_fw | Chunk_9.2C_rv |
| **BT** | Chunk 9.2D | AQ | 2502 | Chunk_9.2D_fw | Chunk_9.2D_rv |
| **AQ** | Chunk 5A | BG | 2485 | Ecoli_ch5_fw | Chunk_5A_rv |

| | | | | | |
|---|---|---|---|---|---|
| **BG** | Chunk 5B | BH | 2521 | Chunk_5B_fw | Chunk_5B_rv |
| **BH** | Chunk 5C | BI | 2513 | Chunk_5C_fw | Chunk_5C_rv |
| **BI** | Chunk 5D | AL | 2441 | Chunk_5D_fw | Ecoli_ch5_rv |
| **AL** | *mTurquoise2* | DS | 1683 | Turquoise_AL_rv | Turquoise_DS_fw |
| **DS** | Telomerator | AO | 807 | Telomerator_r_fw | Telomerator_l_rv |

#### Table S34 Assembly of Syn12 synthetic chromosome.

Colonies were checked by FACS on mRuby2 and mTurquoise2 fluorescence (Suppl. Fig. S6). Correct size of ~100 Kb was checked by CHEF (Suppl. Fig. S9). Correct assembly was checked by Illumina WGS.

| | ***mRuby2*** | ***mTurquoise2*** | **CHEF** | **Correct by sequencing** |
|---|---|---|---|---|
| **Syn12.1** | YES | YES | Correct | YES |
| **Syn12.2** | NO | YES | - | - |
| **Syn12.3** | YES | YES | Correct | YES |
| **Syn12.4** | YES | YES | Correct | YES |
| **Syn12.5** | YES | NO | - | - |
| **Syn12.6** | NO | YES | - | - |
| **Syn12.7** | YES | YES | Correct | NO: Chunk 4D and 5C missing |
| **Syn12.8** | YES | YES | Correct | YES |
| **Syn12.9** | YES | YES | Smaller | - |
| **Syn12.10** | YES | YES | Smaller | - |
| **Syn12.11** | YES | YES | Not visible | - |

#### Figure S1 Microscope imaging of SynChs

Sample images captured with fluorescence microscopy (40x). IMF6, IMF1 and IMF2 contained a putative correct assembly of SynCh1 (for IMF6 and IMF1) and SynCh2 respectively, based on fluorescence for the red, green and blue channels. mRuby2 control: IMC111, Venus control: IMC113, mTurquoise2 control: IMC112, Negative control: CEN.PK113-7D. PC: Phase Contrast For visibility red channel images were adjusted by 60% in brightness. The red bar in the bottom right of the fluorescence microscopy pictures (40x) represents 10 µm.

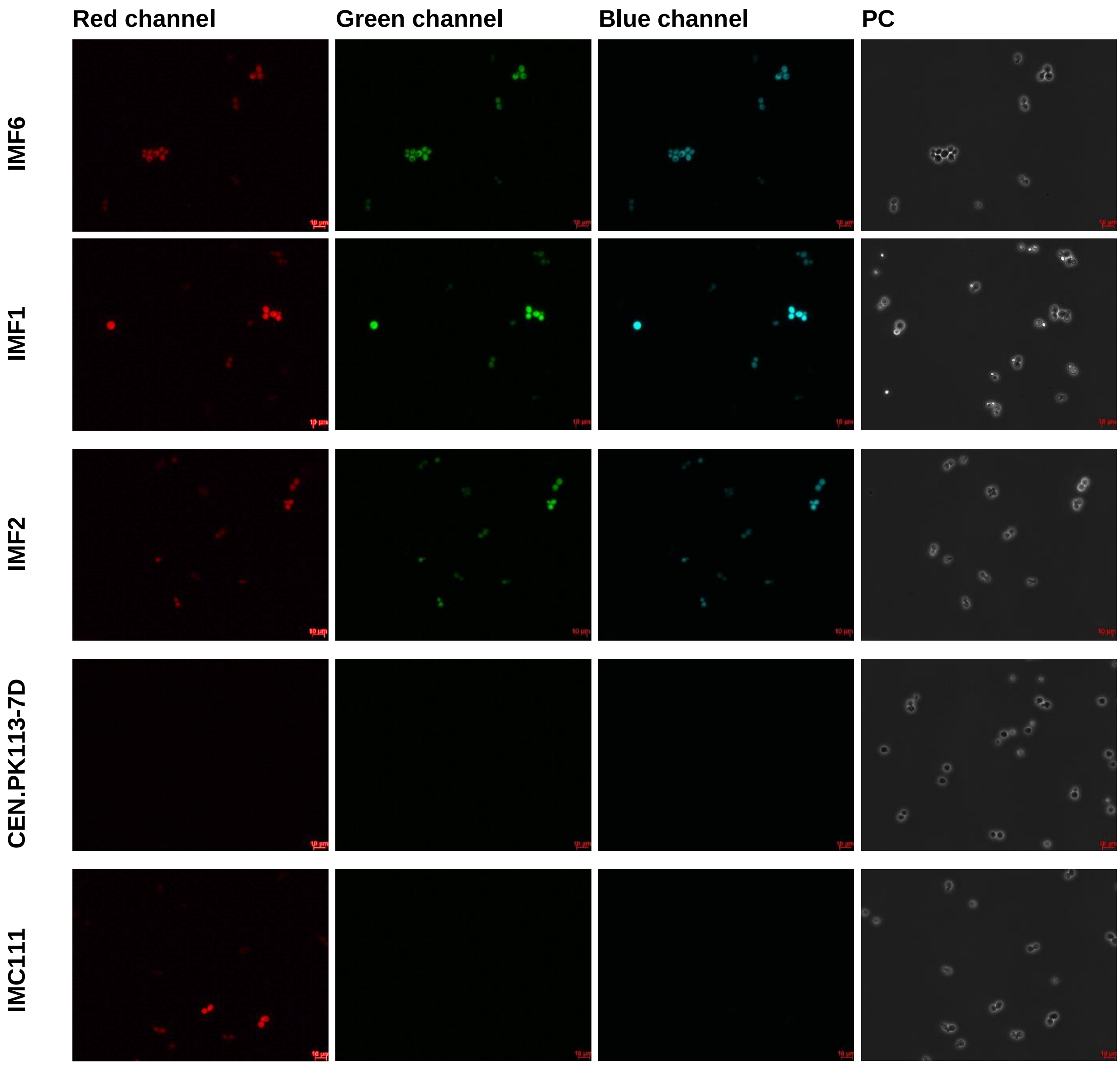

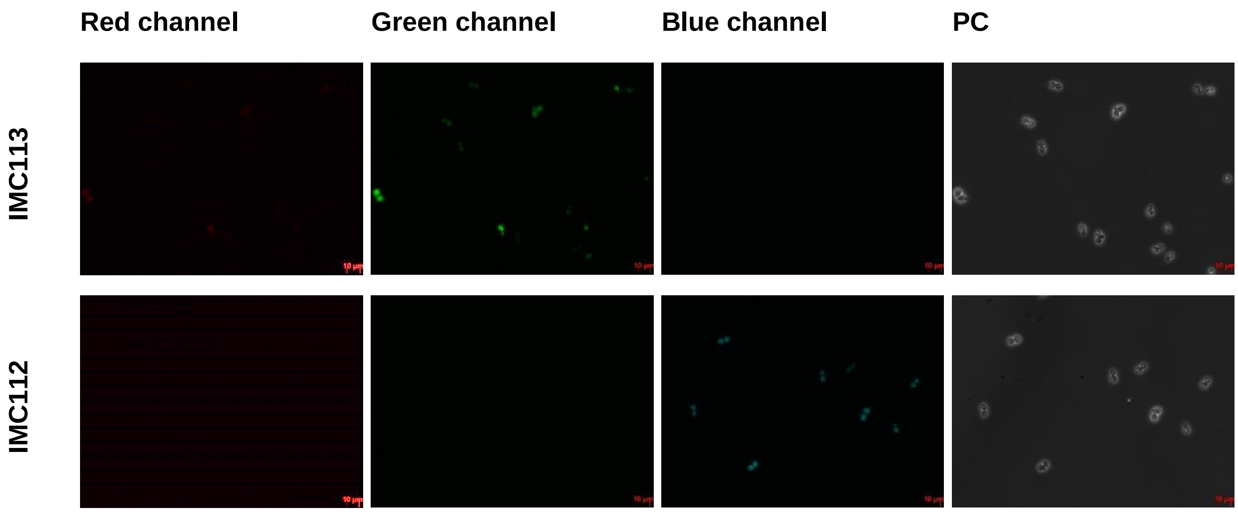
Red channel
Green channel
Blue channel
PC
IMC113
IMC112

#### Figure S2 Strategy to test correct assembly of a SynCh by diagnostic PCR.

The IMF1, IMF2 and IMF6 SynCh were initially checked by diagnostic PCR. Correct assembly of adjacent fragments was checked in three possible ways. Most often amplification of a 5 Kb fragment confirming two adjacent *E.coli* chunks (green). Otherwise an amplification of 10 Kb fragment spanning 4 *E.coli* chunks (blue). Lastly, an amplification of a short PCR fragment spanning over an SHR (red).

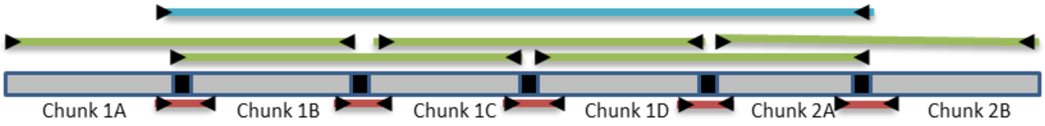

#### Figure S3 Sequencing: coverage and magnolya plots of SynChs

WGS data from strains carrying SynCh. Both the coverage plot (left) and the Magnolya plot (right) for each SynCh are represented. The y-axis of the coverage plots represent the read coverage (a.u.). For IMF1, a whole segment in the middle of the chromosome from the middle of *mTurquoise2* to the middle of *Venus* is duplicated. All other strains harbour all fragments of the SynChs in the correct order. For parts in SynCh the coverage is 0 or 0.5 (ex. IMF18 and IMF17). This is because CEN.PK113-7D was used as reference strain to align to the sequencing reads to. This reference contains promoters/genes/terminators which are absent in the actual strain (IMX1338) in which the SynChs were constructed. Therefore reads will (also) map to this native location instead of to the SynCh. All these sites were verified to be present in the particular SynCh. From the Magnolya plots it can be observed that the SynCh in IMF2 is present on average in 0.6 copies/cell, for IMF12 and IMF6 on average in 1.5 copies/cell, for IMF11 on average in 1.7 copies/cell and for IMF1 on average in 2.0 copies/cell. The SynChs: Syn12.1 (IMF24), Syn12.3 (IMF25), Syn12.4 (IMF26) and Syn12.8 (IMF23) are on average in 1 copy/cell, as well as the SynCh in IMF18 and IMF17.

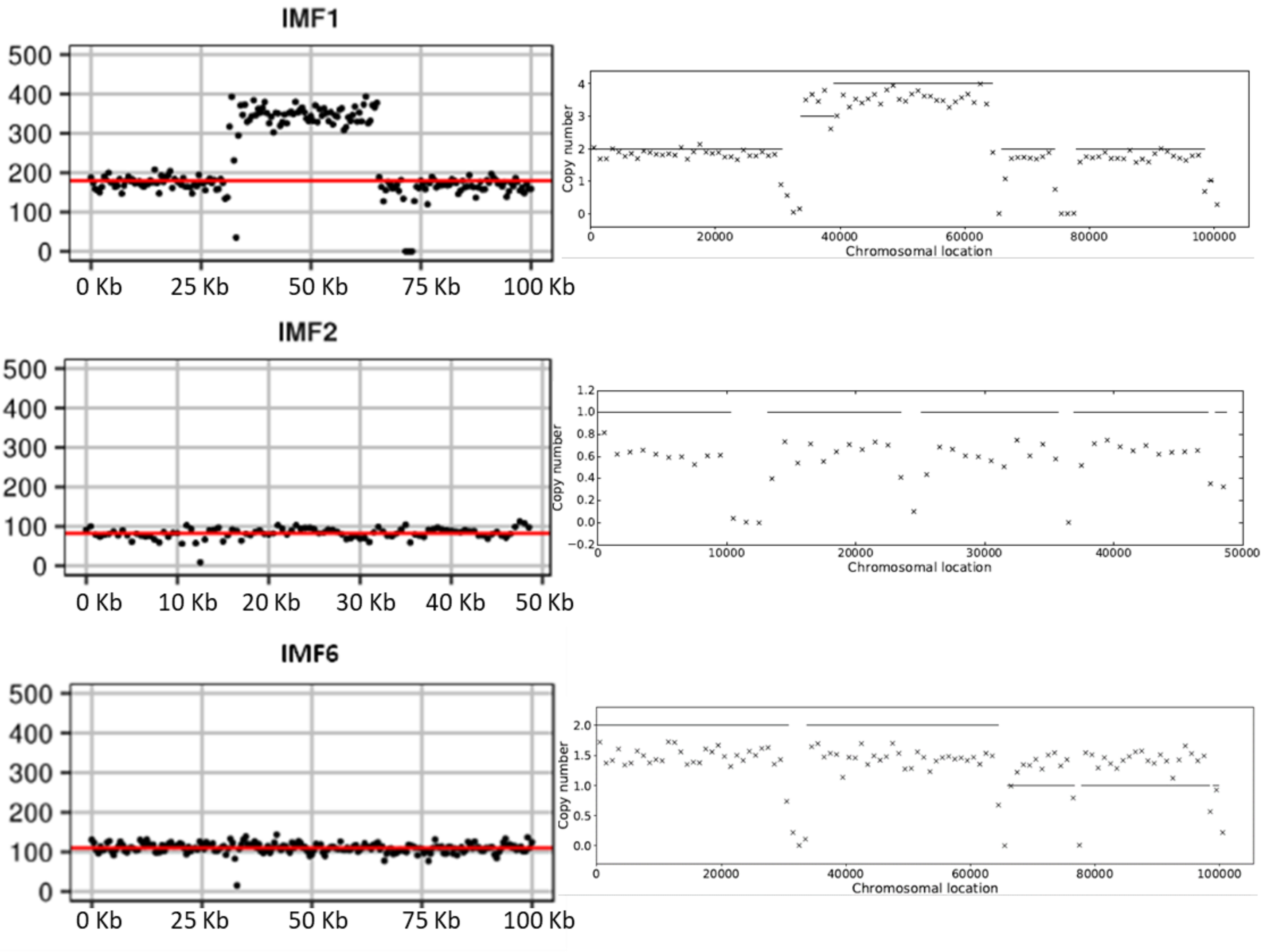

#### IMF11

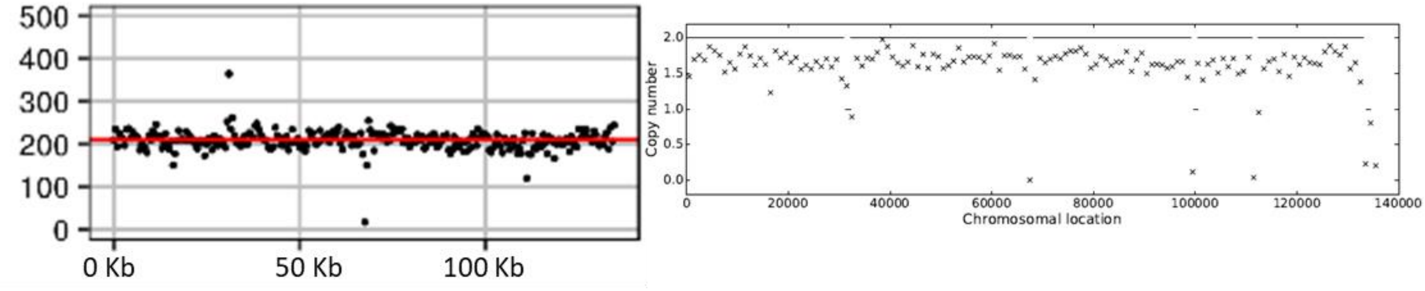


#### IMF12

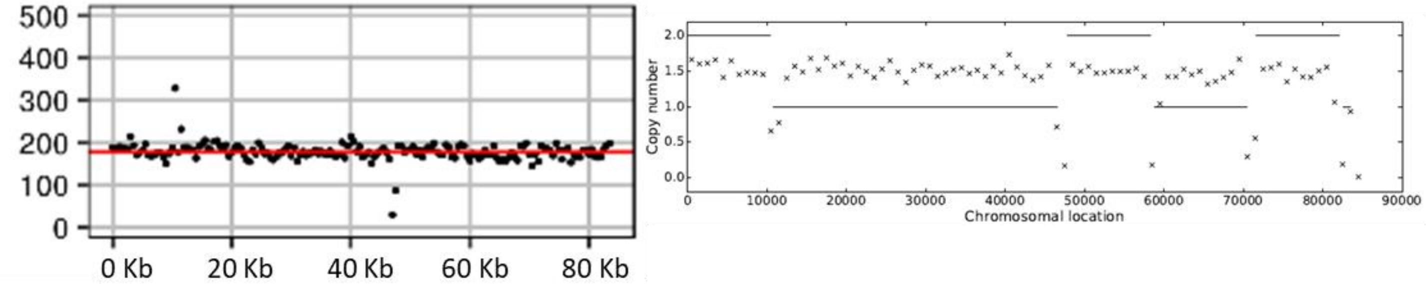


#### IMF17

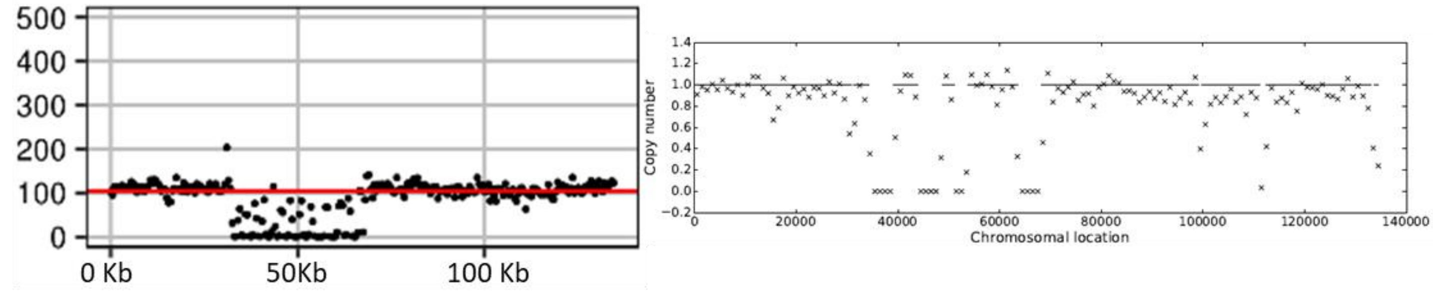


#### IMF18

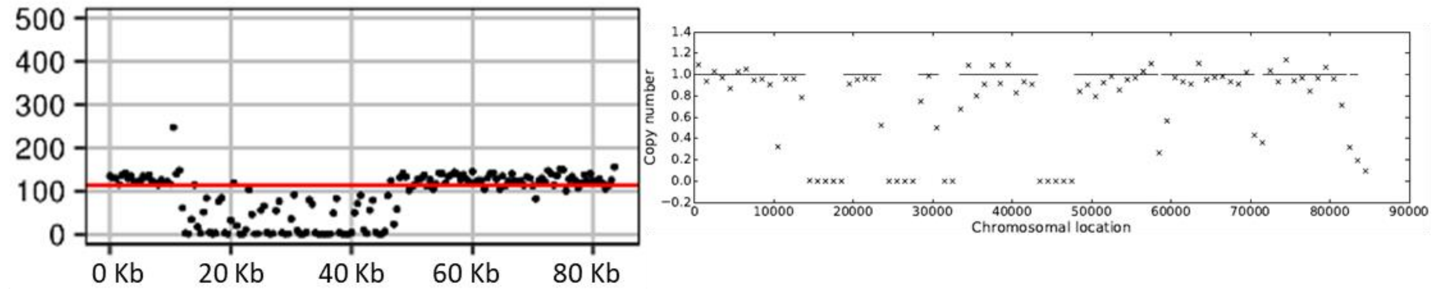

Syn12.1

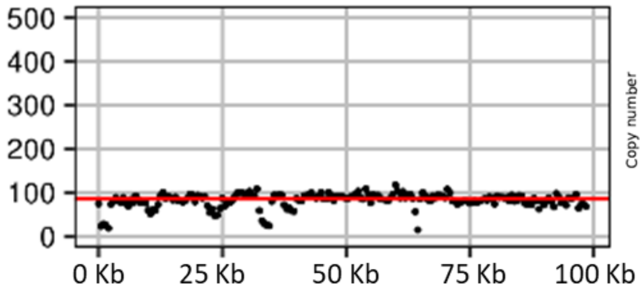


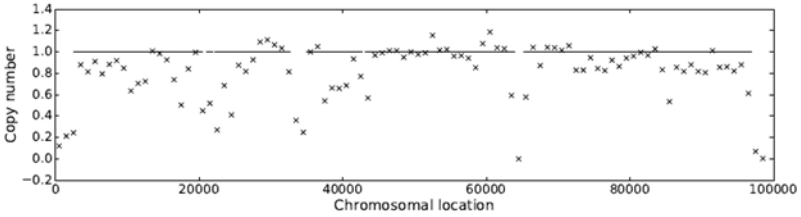


Syn12.3

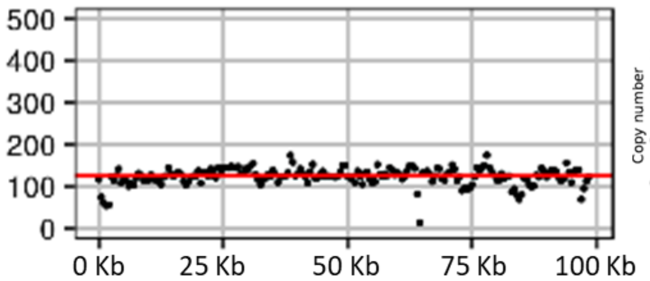


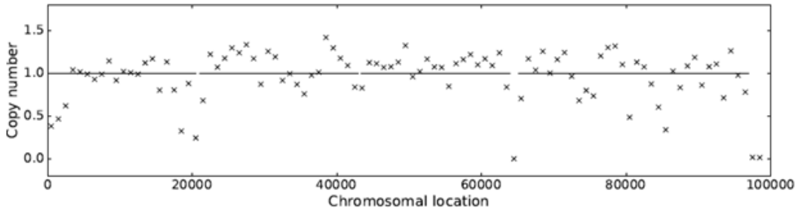


Syn12.4

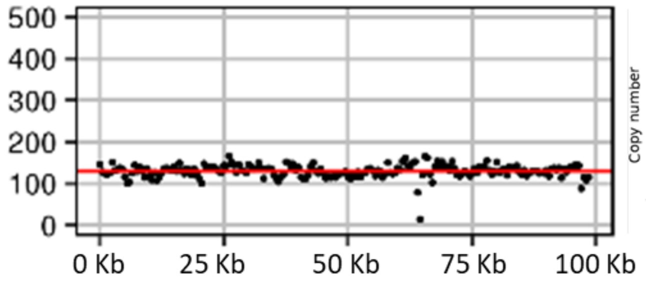


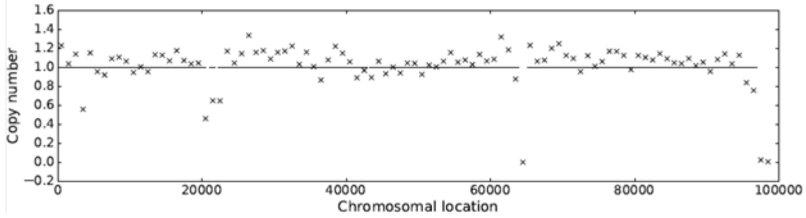


Syn12.8

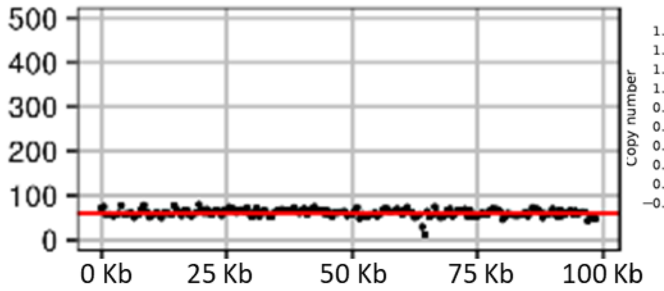


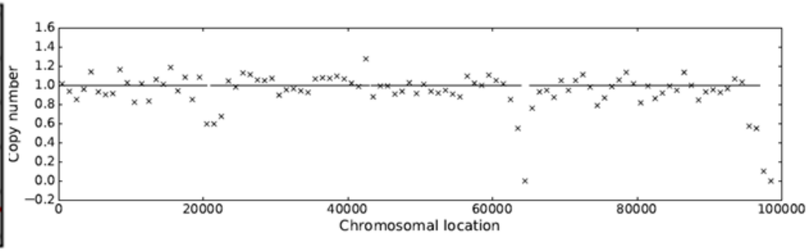

#### Figure S4 CHEF analysis of IMF1 and IMF2

CHEF analysis of IMF1 and IMF2. 1)CEN.PK113-5D (- control), 2)IMX1338 (- control), 3)IMS0480 (+ control), 4)in plug I-SceI digested IMX212, 5)in plug I-SceI digested (1h) IMF1, 6) in plug I-SceI digested (O/N) IMF1, 7)In plug NotI digested IMF1, 8) In plug I-SceI digested IMF2, 9) Lambda PFG ladder. SynCh1 in IMF1, shows it is approximately 135 Kb, which is 35 Kb larger than the *in silico* design. SynCh2 in IMF2 has the size of 50 Kb as *in silico* designed.

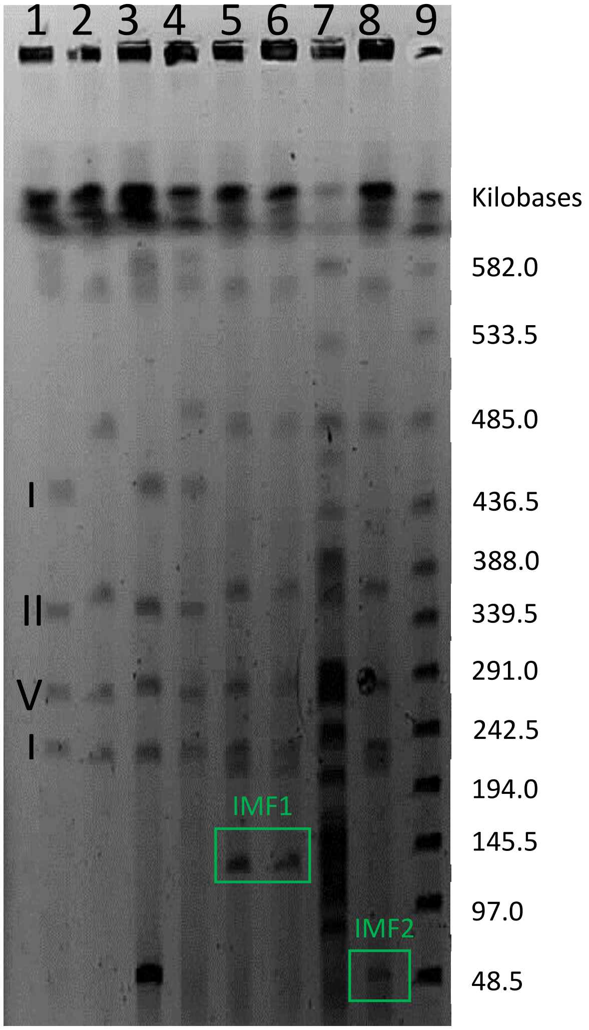

**Figure S5 CHEF IMF1, IMF2 and IMF6**

1)CEN.PK113-5D (- control), 2)IMX1338 (- control), 3)IMS0480 (+ control), 4)in plug I-SceI digested IMF1, 5) in plug I-SceI digested IMF2, 6) in plug I-SceI digested IMF1, 7) undigested transformant 3 (IMF6), 8) in plug I-SceI digested transformant 3 (IMF6), 9) undigested transformant 4, 10) in plug I-SceI digested transformant 4., 11) Lambda PFG ladder. SynCh1 in IMF1, shows it is approximately 135 Kb, which is 35 Kb larger than the in silico design. SynCh2 in IMF2 has the size of 50 Kb as *in silico* designed. SynCh1 in transformant 3 and 4 both show a correct size of approximately 100 Kb.

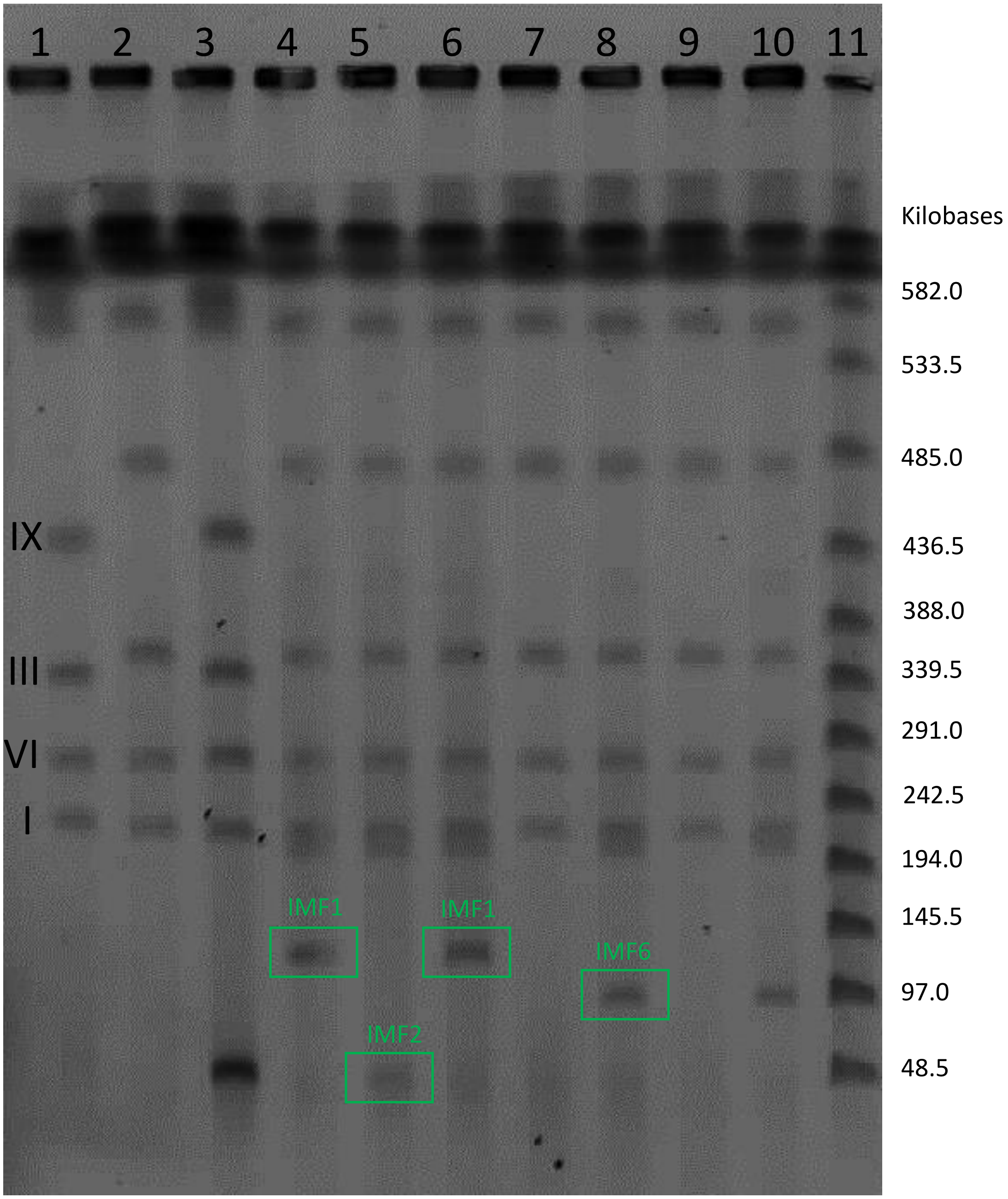

#### Figure S6 Syn12 assembly

The assembly of 43 fragments into a 100 Kb Syn12 chromosome. Chunks represent 2.5 Kb non-coding *E.coli* DNA. *HIS3*, *URA3* (in telomerator), *mRuby2* and *mTurquoise2* were used as markers. CEN6/ARS4 was included for segregation of the SynCh and *ARS1* and *ARS417* were included to ensure correct replication.

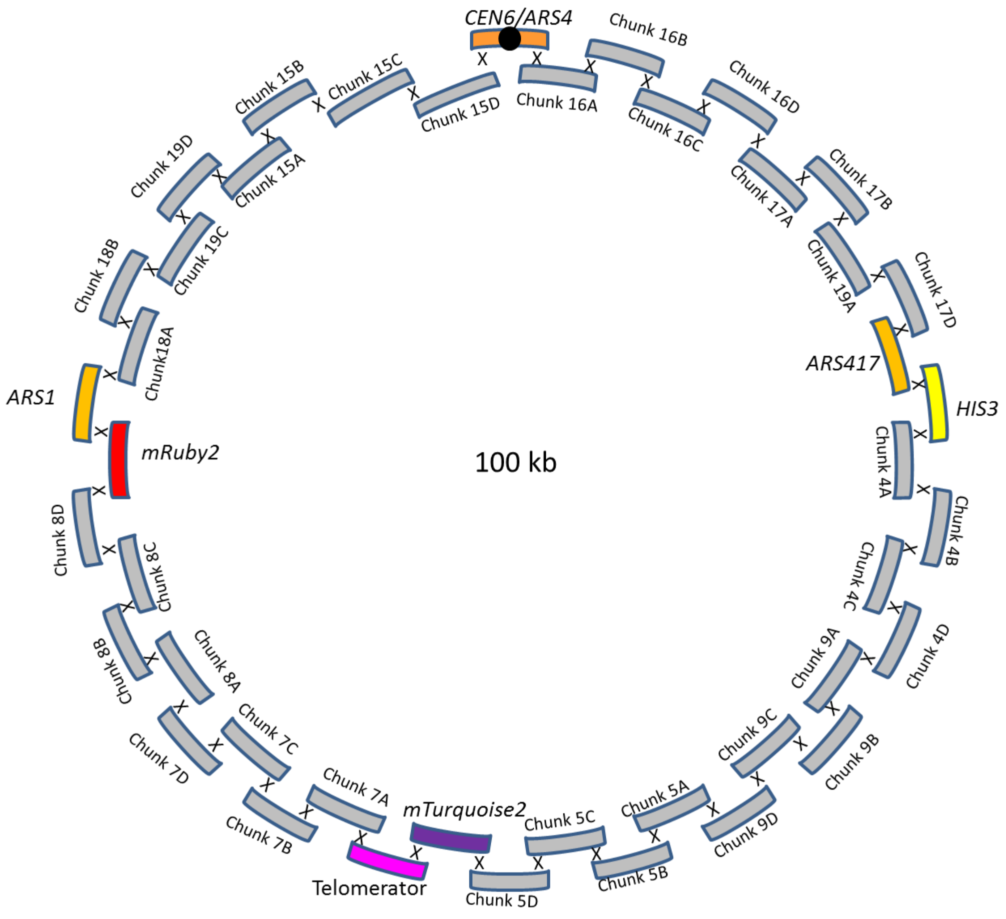

#### Figure S7 FACS imaging of Syn12 transformants

FACS imaging of SynChs. The fluorescence is plotted on the y-axis and the FSC-A on the x-axis. Negative control: CEN.PK113-7D. Positive controls: IMC112(mRuby2), IMC111 (mTurquoise2). Gates for fluorescence of the two different fluorescent proteins were drawn based on the IMC111 and IMC112 controls. Double fluorescence: Syn12.1, Syn12.3, Syn12.4, Syn12.7, Syn12.8, Syn12.9, Syn12.10 and Syn12.11. mRuby2 fluorescence only: Syn12.5. mTurquoise2 fluorescence only: Syn12.2 and Syn12.6. Approximately 10000 events are shown.

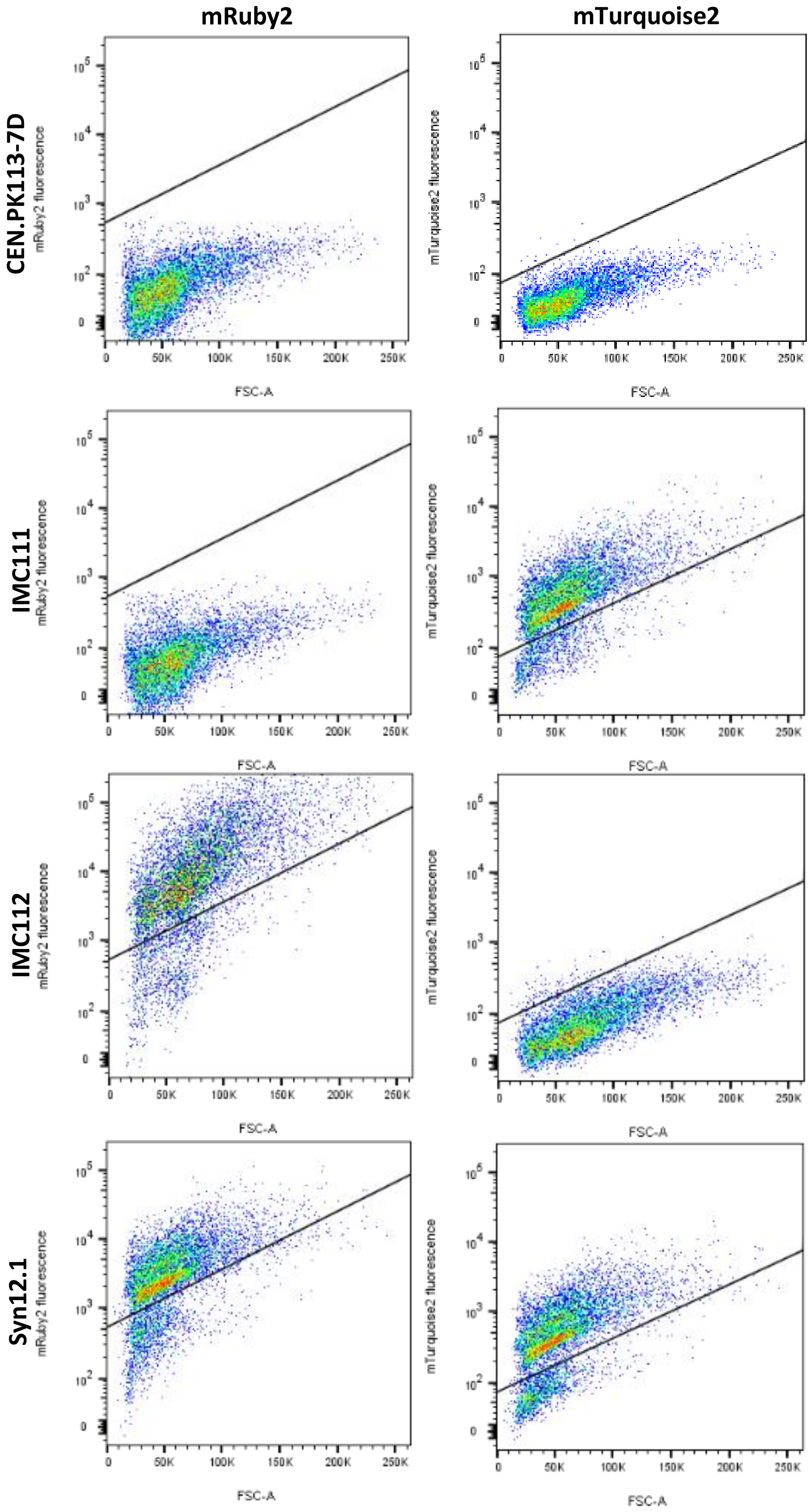

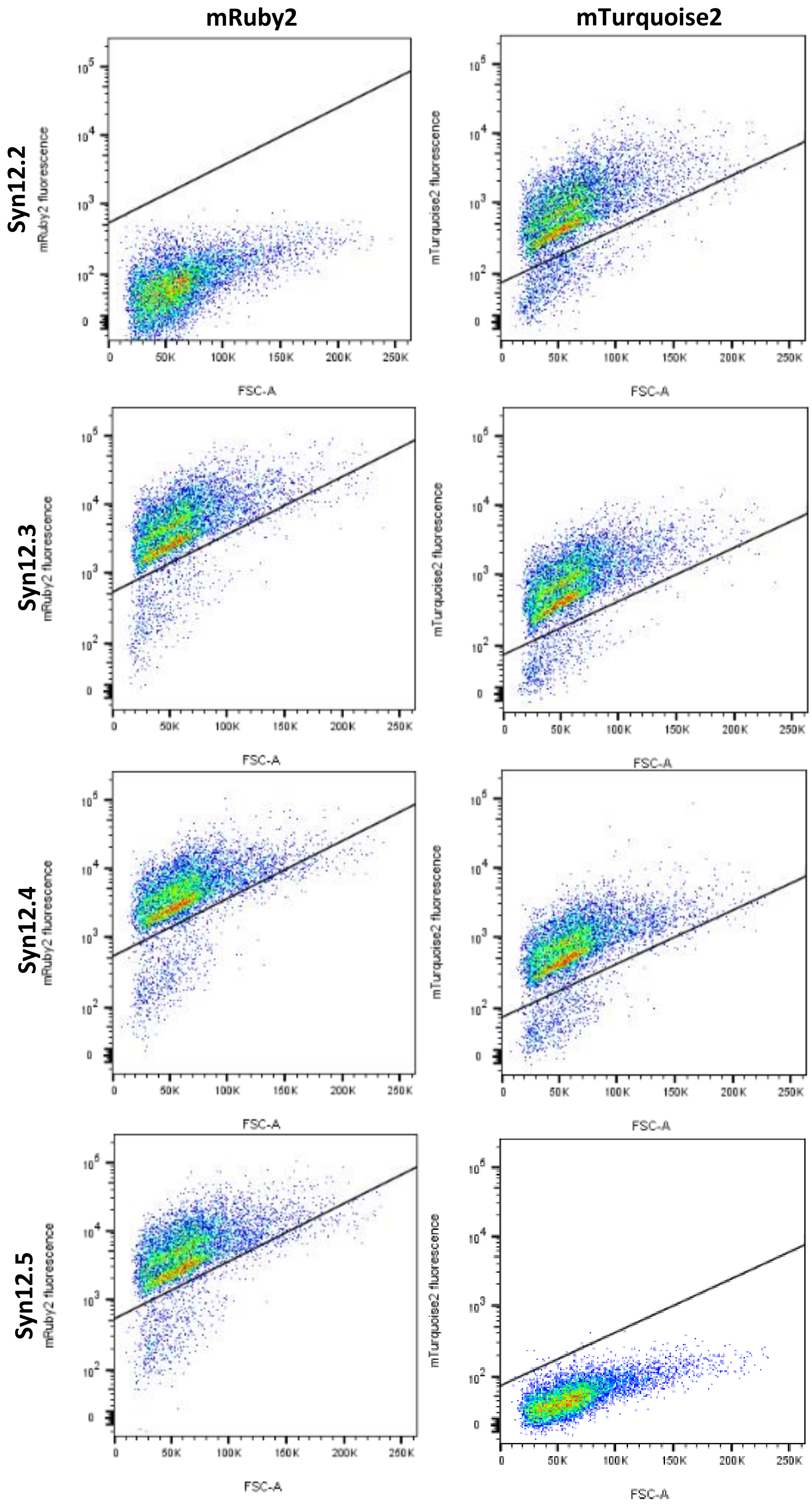
mRuby2
mTurquoise2
Syn12.2
Syn12.3
Syn12.4
Syn12.5
mRuby2 fluorescence
mTurquoise2 fluorescence
FSC-A
0
50K
100K
150K
200K
250K
$10^2$
$10^3$
$10^4$
$10^5$

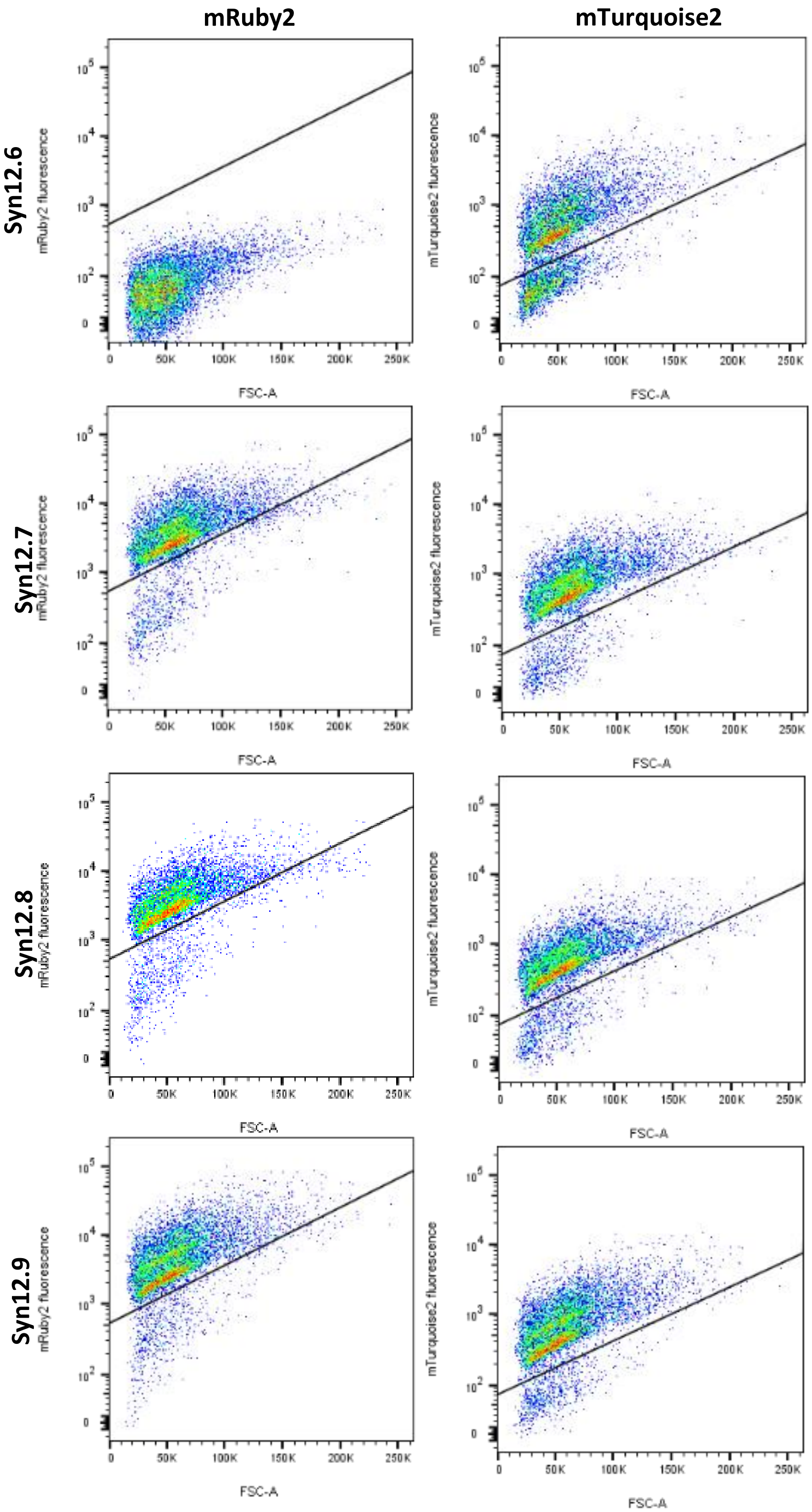
mRuby2
mTurquoise2
Syn12.6
Syn12.7
Syn12.8
Syn12.9
mRuby2 fluorescence
mTurquoise2 fluorescence
FSC-A
0
50K
100K
150K
200K
250K
$10^2$
$10^3$
$10^4$
$10^5$

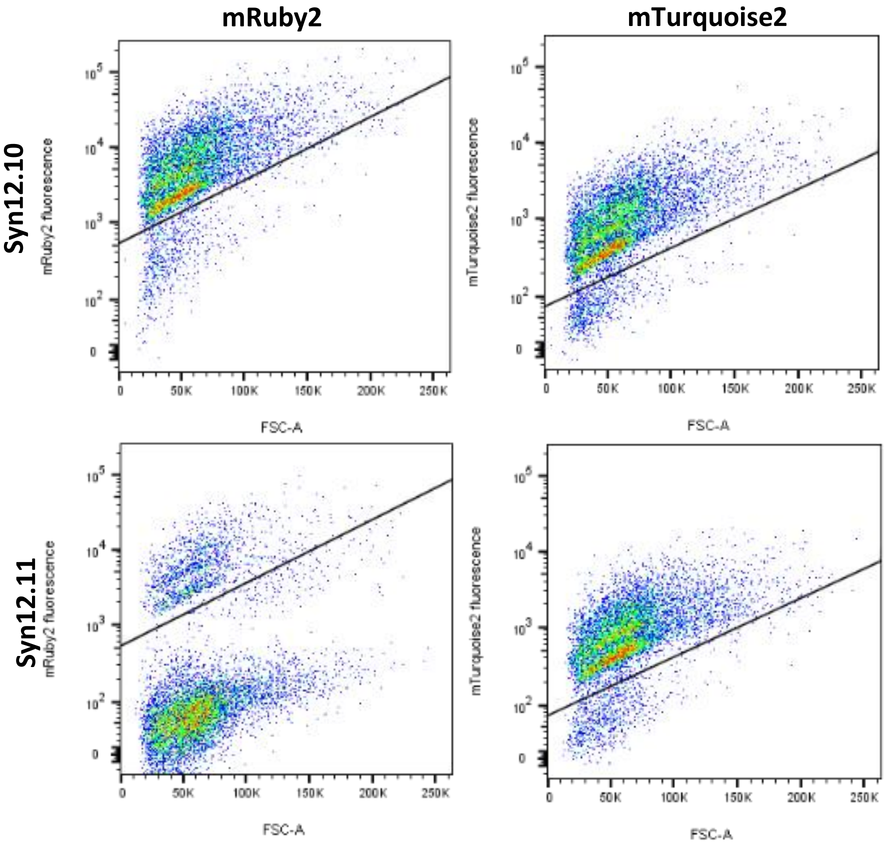
mRuby2
mTurquoise2
Syn12.10
Syn12.11
mRuby2 fluorescence
mTurquoise2 fluorescence
FSC-A

#### Figure S8 CHEF analysis of Syn12 transformants.

CHEF analysis of Syn12 transformants. 1)CEN.PK113-7D (- control), 2)IMX1338 (- control), 3) in plug I-SceI digested IMF6 (+ control) 4)in plug I-SceI digested Syn12.1 (IMF24), 5) in plug I-SceI digested Syn12.3 (IMF25), 6) in plug I-SceI digested Syn12.4 (IMF26), 7) in plug I-SceI digested Syn12.7, 8) in plug I-SceI digested Syn12.8 (IMF23), 9) in plug I-SceI digested Syn12.9, 10) in plug I-SceI digested Syn12.10, 11) in plug I-SceI digested Syn12.11, 12) Lambda PFG ladder. Transformants 1, 3, 4, 7 and 8 appear to have a SynCh of the correct size of 100 Kb. Transformants 9 and 10 have a smaller SynCh than 100 Kb. There seems to no band for a SynCh for transformant 11

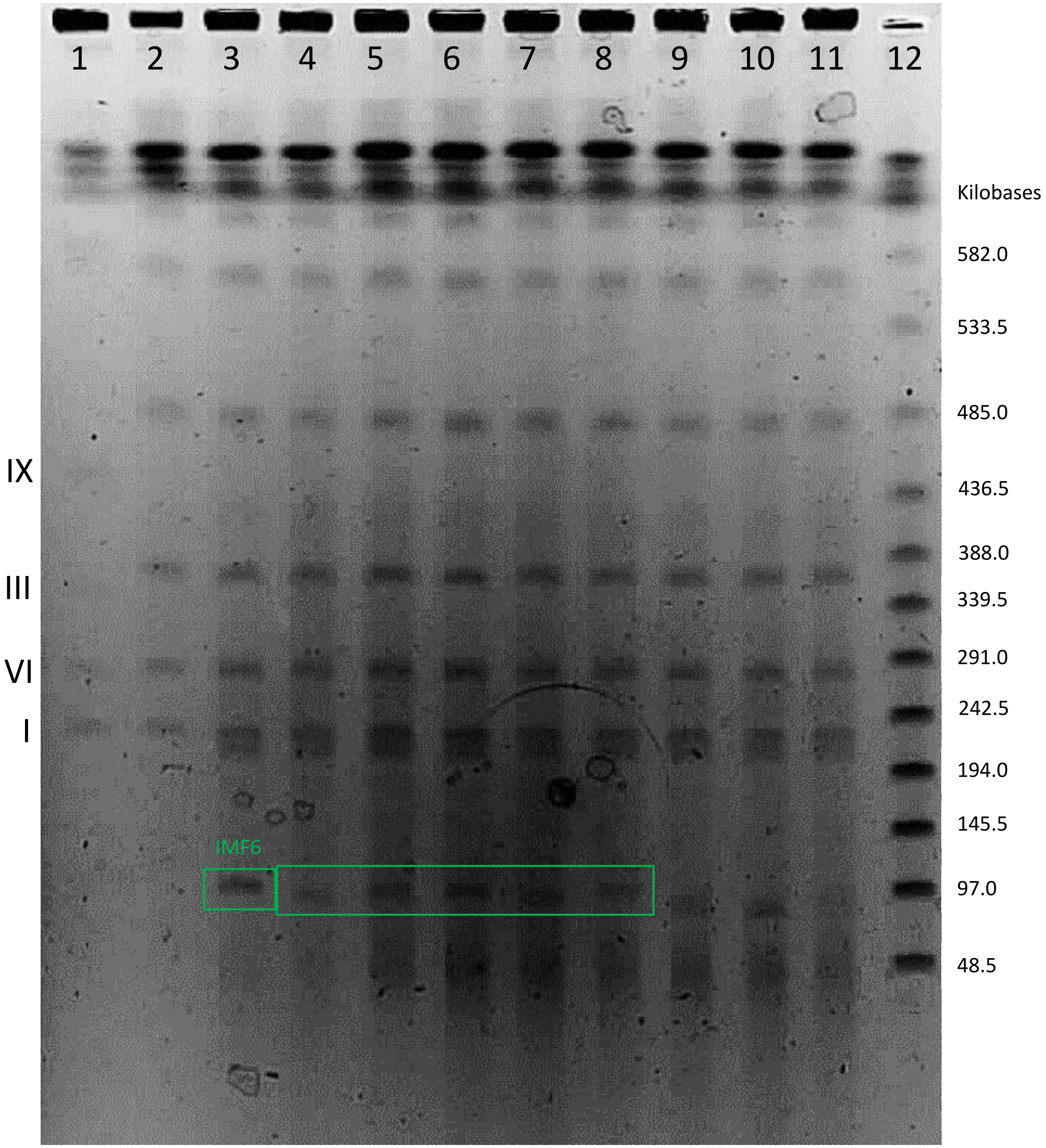
1
2
3
4
5
6
7
8
9
10
11
12
Kilobases
582.0
533.5
485.0
436.5
388.0
339.5
291.0
242.5
194.0
145.5
97.0
48.5
IX
III
VI
I
IMF6

#### Figure S9 FACS imaging of SynChs

FACS imaging of SynChs. The fluorescence is plotted on the y-axis and the FSC-A on the x-axis. The Negative control: CEN.PK113-7D. Postive controls: IMX2224 (mRuby2), IMX2225 (Venus), IMX2226 (mTurquoise2). The Venus protein showed bleed through in the mTurquoise2 channel, this was compensated based on the compensation matrix in FlowJo V10 with IMX2226 as postive signal and IMX2225 as negative signal. Gates for fluorescence of the three different fluorescent proteins were drawn based on the IMX2224, IMX2225 and IMX2226 controls. IMF2 (50 Kb, empty) showed all three fluorescence. IMF6 (100 Kb, empty) showed all three fluorescence. IMF11 (135 Kb, empty) showed mRuby2 fluorescence and Venus fluorescence but no mTurquoise2 fluorescence. IMF12 (85 Kb, empty) showed mRuby2 fluorescence and Venus fluorescence but no mTurquoise2 fluorescence. IMF17 (135 Kb, glycolysis) showed mRuby2 fluorescence and Venus fluorescence but no mTurquoise2 fluorescence. IMF18 (85 Kb, glycolysis) showed mRuby2 fluorescence and Venus fluorescence but no mTurquoise2 fluorescence. Approximatly 100000 events are shown.

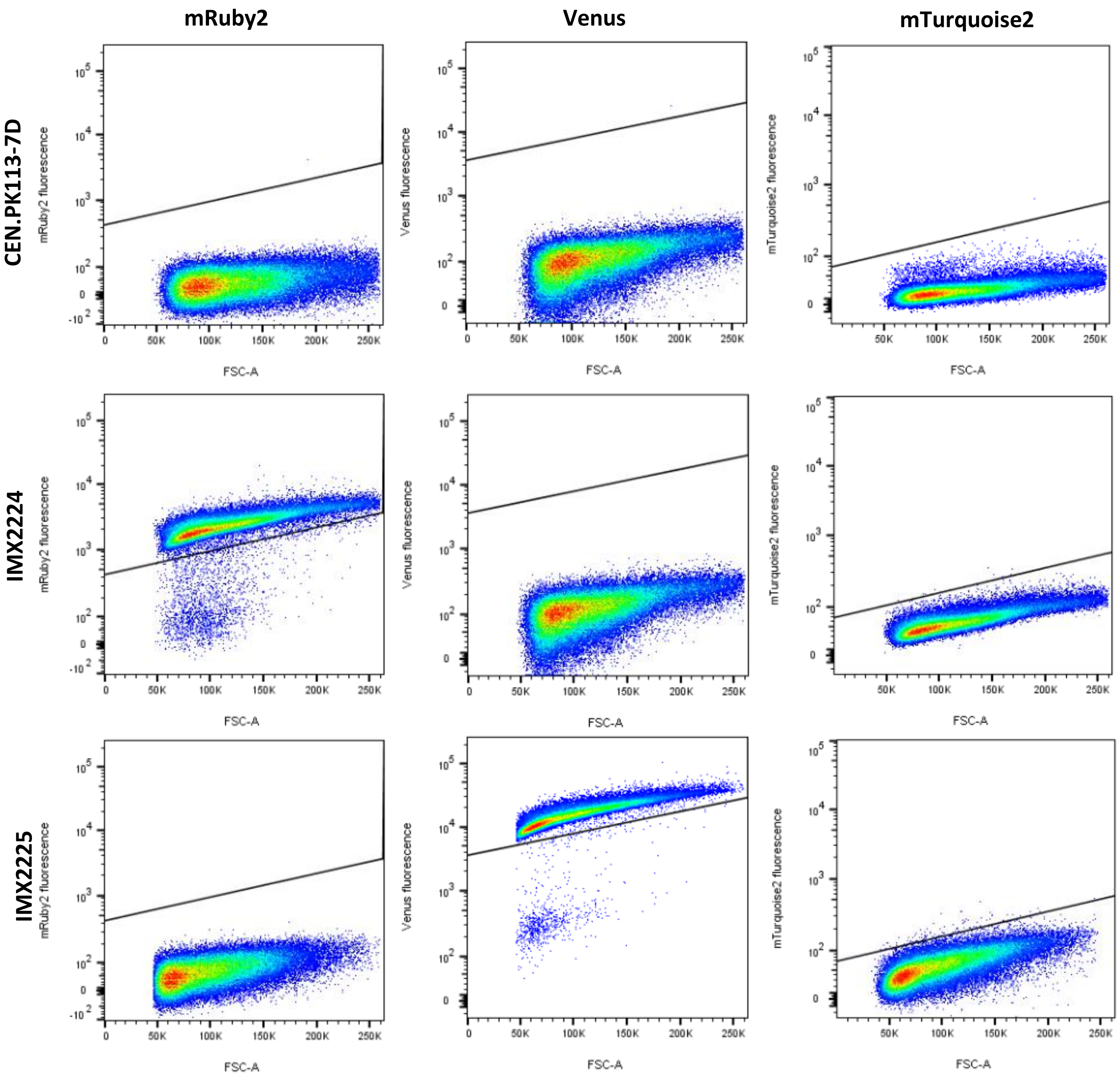

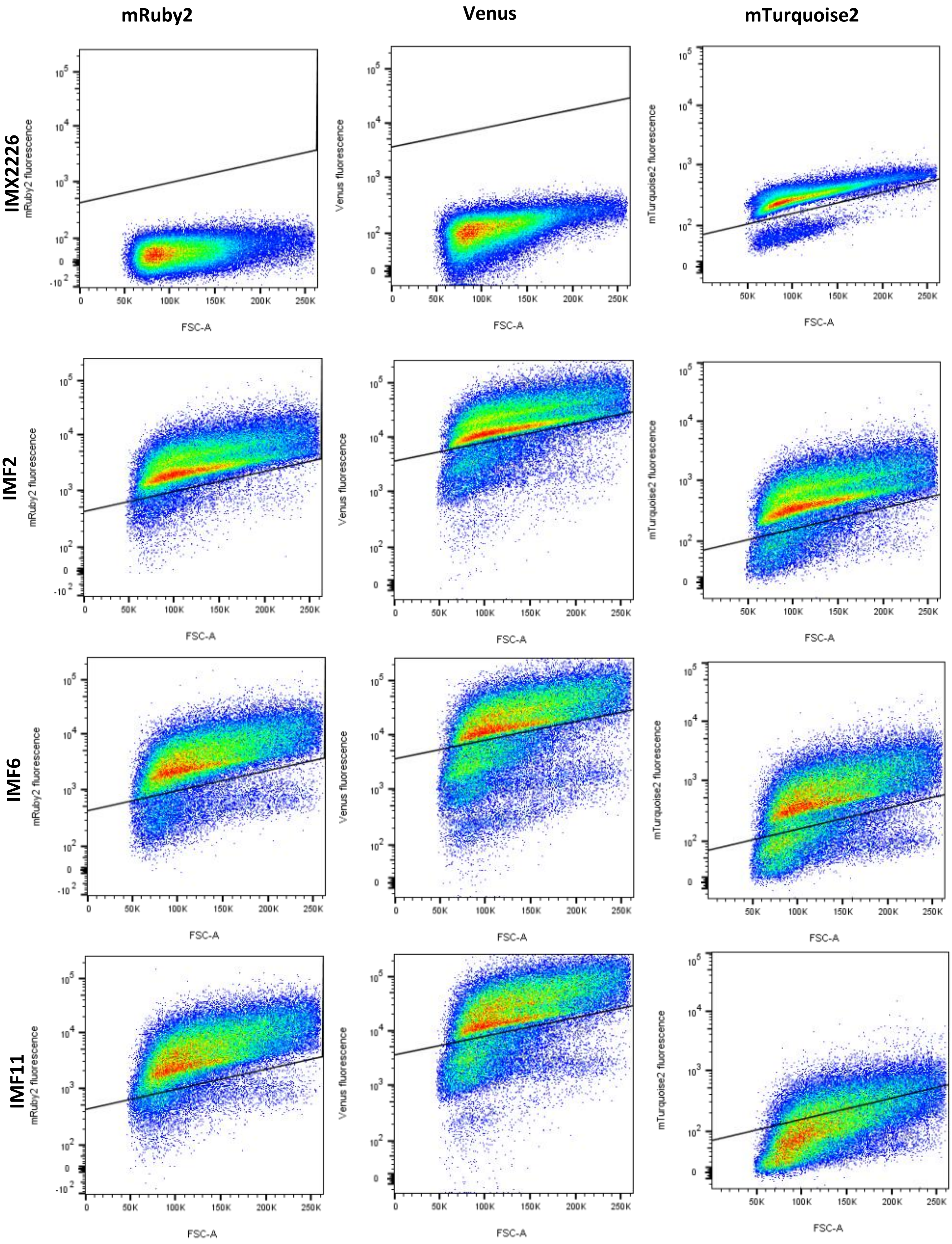
mRuby2
Venus
mTurquoise2
IMX2226
IMF2
IMF6
IMF11
mRuby2 fluorescence
Venus fluorescence
mTurquoise2 fluorescence
FSC-A

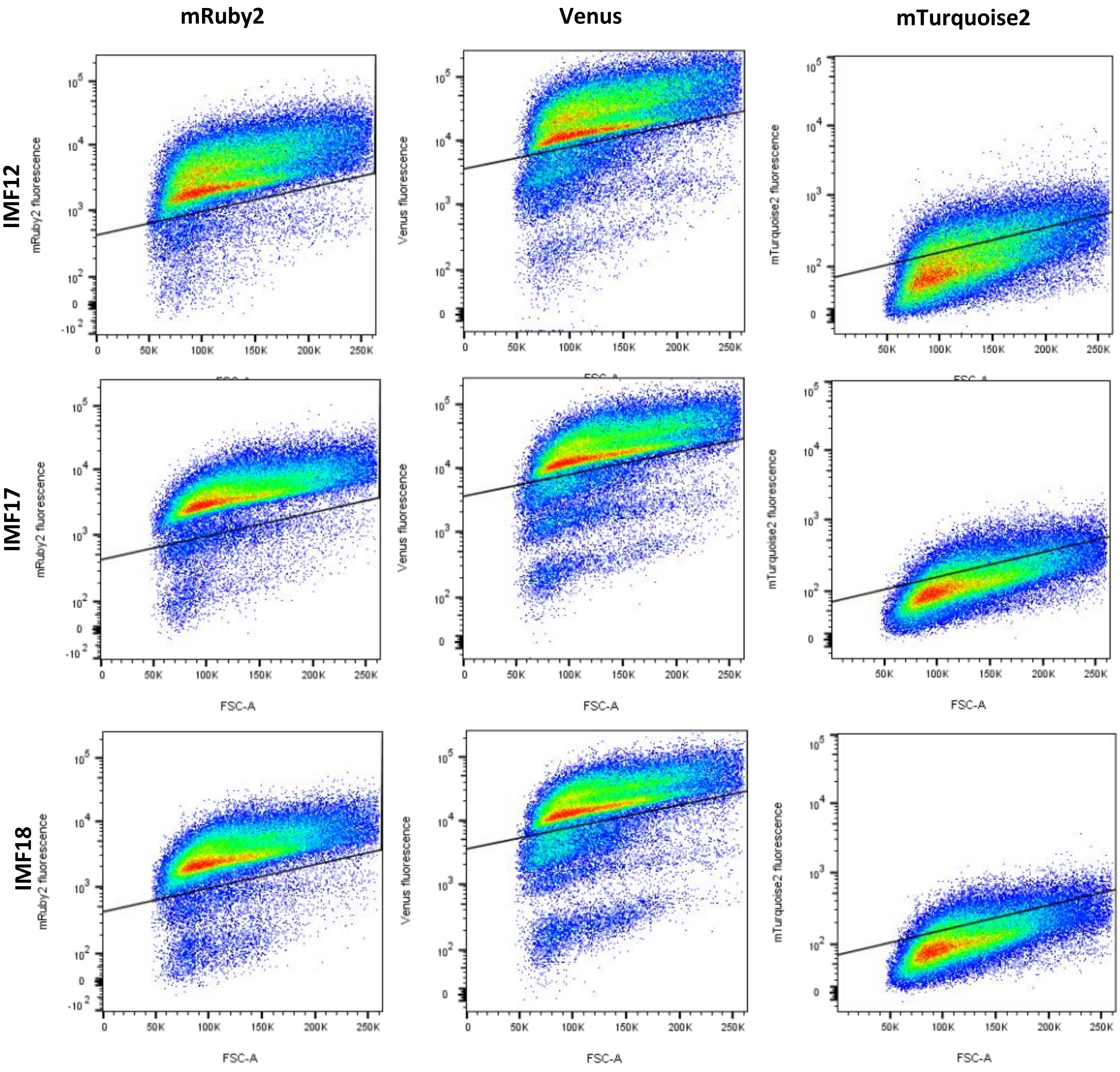
mRuby2
Venus
mTurquoise2
IMF12
IMF17
IMF18
mRuby2 fluorescence
Venus fluorescence
mTurquoise2 fluorescence
FSC-A

**Figure S10 CHEF analysis of IMF2, IMF12, IMF18, IMF6, IMF11, IMF17**

1) Lambda PFG ladder, 2)CEN.PK113-7D (- control), 3)IMX1338 (- control), 4) in plug I-SceI digested IMF2 (50 Kb SynCh), 5)in plug I-SceI digested IMF12 (85 Kb SynCh), 6) in plug I-SceI digested IMF18 (85 Kb SynCh containing SinLoG), 7) in plug I-SceI digested IMF6 (100 Kb SynCh), 8) in plug I-SceI digested IMF11 (135 Kb SynCh), 9) in plug I-SceI digested IMF17 (135 Kb SynCh containing SinLoG), 10)Lambda PFG ladder.

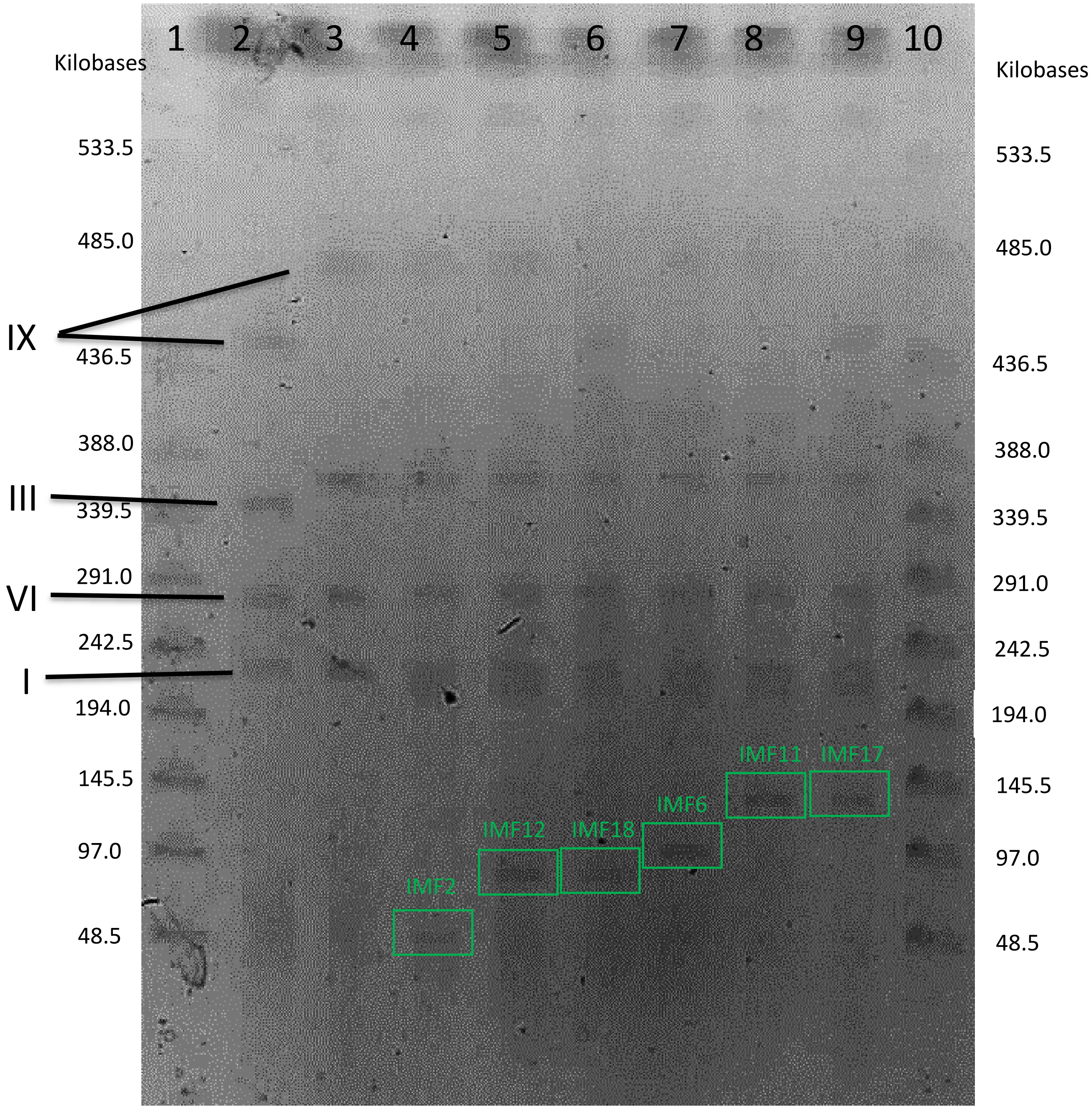

**Figure S11 SynCh stability upon transfer.**

The stability of the SynCh calculated by:

$$\%\ \boldsymbol{of\ cells\ with\ SynCh}: \frac{\boldsymbol{number\ of\ colonies\ on\ SMD}}{\boldsymbol{number\ of\ colonies\ of\ YPD}} \times \mathbf{100\%}$$

IMF18 day 1: only in single measurements
IMF11 day 1: no measurements
IMF17 day 1: no measurements
IMF23: only measurements on day 1 and 4

For CEN.PK113-7D, IMF18 (50 Kb, glycolysis), IMF17 (50 Kb, glycolysis) the percentage is approximatly 100% for all measurments. This is due to the fact that SMD and YPD medium are both selective media with respect to the (synthetic) chromosome(s) that they carry. For IMF23 (100 Kb empty, improved design) and IMC153 (6.5 Kb plasmid) the percentage of cells with the SynCh/plasmid is high on all days meaning it is very stable. For IMF2 (50 Kb, empty), IMF12 (85 Kb, empty), IMF6 (100 Kb, empty) and IMF11 (135 Kb, empty) the percentage of cells with SynCh was lower on all days tested (around 80%), revealing reduced stability of the respective SynChs.

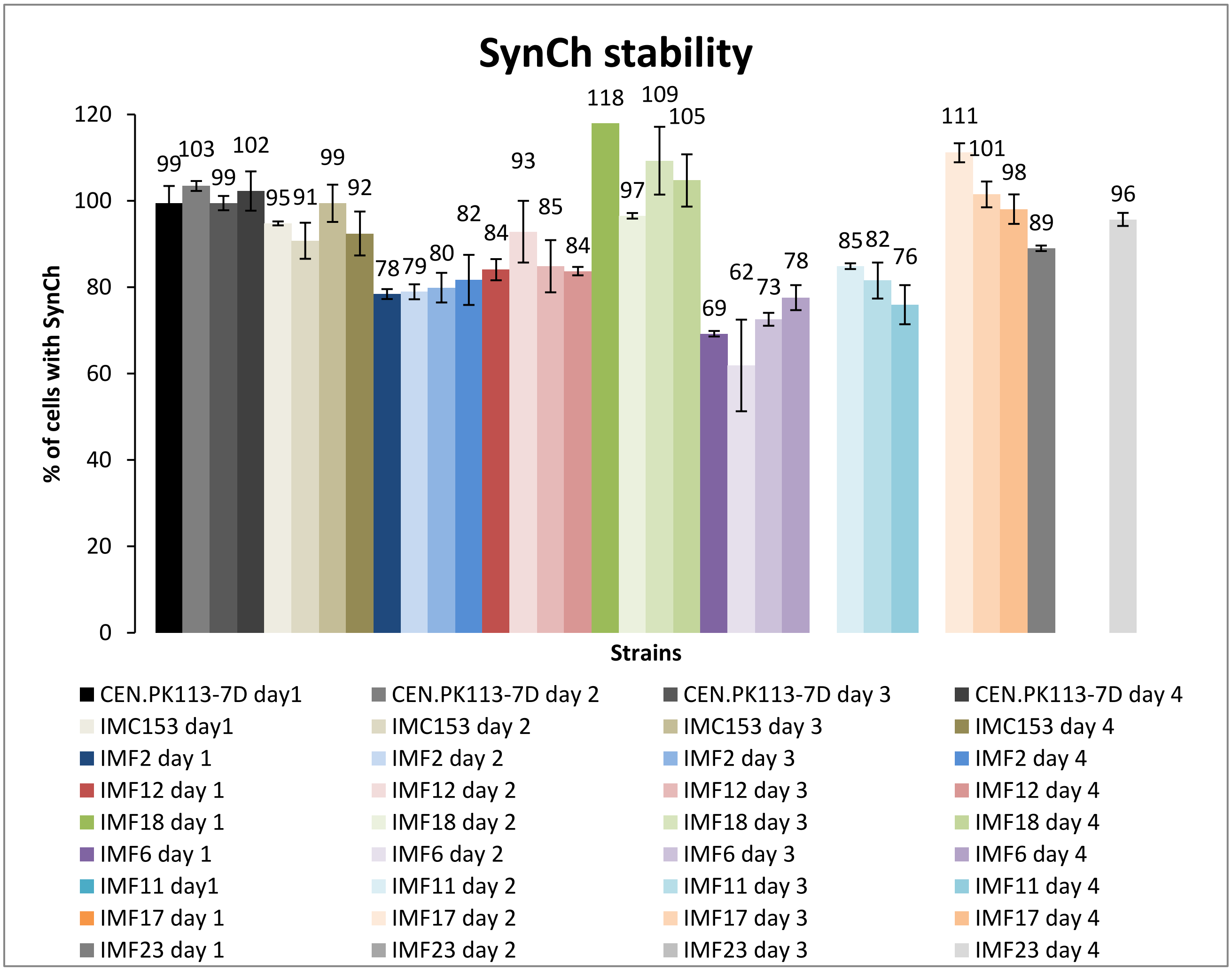

**Figure S12 Amplification strategy of *E.coli* fragments (Chunks) of various lengths.** Schematic representation of 5 Kb and 10 Kb chunk amplification from a 2.5 Kb chunk assembled chromosome (strain IMF1). This was used for the assembly of SynCh3.2, SynCh3.3, SynCh4.2 and SynCh4.3 (Suppl. Table S28-S29 and S31-S32)

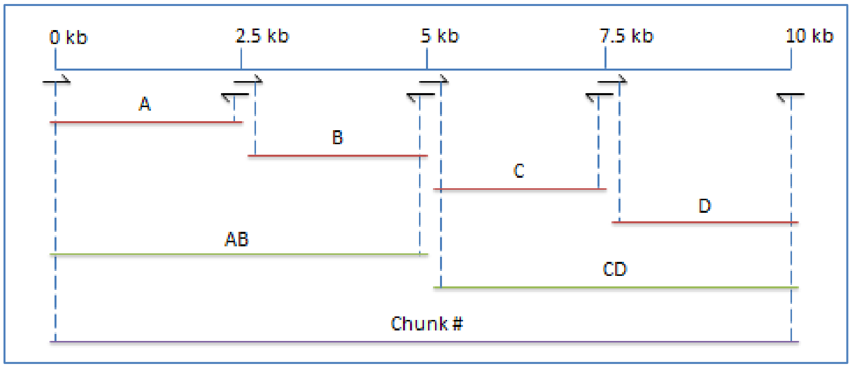

#### Figure S13 Confirmation mixed population of *E.coli* for template isolation *of E.coli chunks* in SynCh.

It was discovered that the culture of *E.coli*, from which genomic DNA was isolated which was used for amplification of *E.coli* chunks for the SynCh, was a mixture of two *E.coli* strains, probably by contamination of the culture. Based on whole genome sequencing data of all SynCh strains and *E.coli* strains, we determined that the two *E.coli* strains were *E.coli* BL21 (used as *in silico* template for the SynCh designs) and *E.coli* XL1-blue. To verify that indeed the base pair variation observed from the E.coli BL21 used as in silico template originated from the *E.coli* XL1-blue genome and did not occur during assembly and propagation of the SynChs, we sanger sequenced part of the PCR fragments which were used in the SynCh transformation. It was verified in all cases that the nucleotide deviation from the *in silico* design (based on *E.coli* BL21) was already present in the template DNA used for transformation and originated from *E.coli* XL1-blue DNA. As an example chunk 1D is represented here, where some SynCh carried the DNA of E.coli XL1 blue and others the sequence of E.coli BL21. Similar results were obtained for all other PCR fragments verified (1B, 1C 2C, 2D, 6A, 8C, 15C)

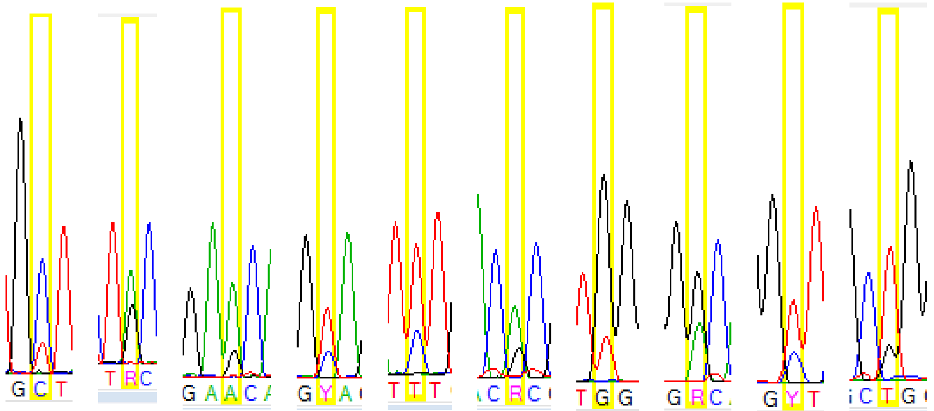

| | | | | | | | | | | |
|---|---|---|---|---|---|---|---|---|---|---|
| BL21 | T | G | G | C | C | G | T | A | C | G |
| XL1-Blue | C | A | A | T | T | A | G | G | T | T |

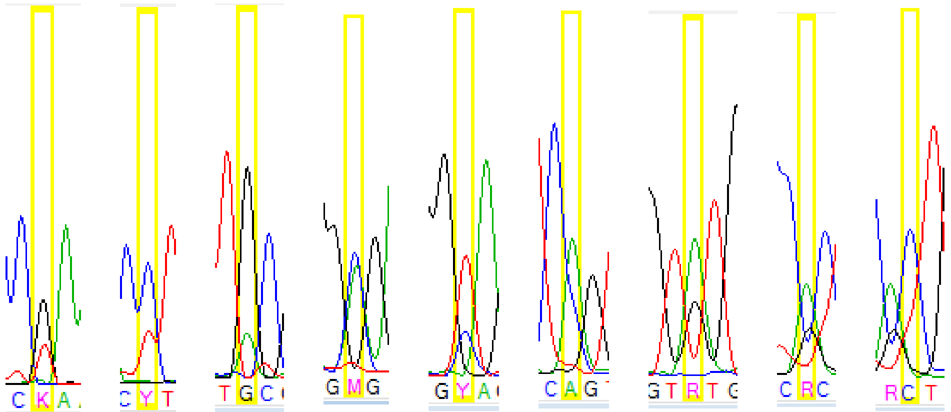

| | | | | | | | | | |
|---|---|---|---|---|---|---|---|---|---|
| BL21 | T | T | A | A | C | C | G | G | T |
| XL1-Blue | G | C | G | C | T | A | A | A | C |

*Represented are sanger sequencing data of parts of PCR fragment of chunk 1D used for transformation of SynCh. In the WGS of these SynCh, nucleotide variations were observed. It was verified that these originated from the template DNA which was a mix of E.coli BL21 DNA and E.coli XL1-blue DNA. Here is confirmed that this mixed DNA was already present in the PCR fragment used for transformation (A=green, G=black, C=blue and T=red)*

#### References


1. Entian, K.-D. and Kötter, P. (2007) 25 Yeast Genetic Strain and Plasmid Collections. In Stansfield, I. and Stark, M. J. R. (eds.), *Methods in Microbiology*. Academic Press, Vol. 36, pp. 629-666.
2. Verduyn, C., Postma, E., Scheffers, W.A. and Van Dijken, J.P. (1992) Effect of benzoic acid on metabolic fluxes in yeasts: A continuous-culture study on the regulation of respiration and alcoholic fermentation. *Yeast*, **8**, 501-517.
3. Gietz, R.D. and Woods, R.A. (2002) Transformation of yeast by lithium acetate/single-stranded carrier DNA/polyethylene glycol method. *Methods Enzymol.*, **350**, 87-96.
4. Looke, M., Kristjuhan, K. and Kristjuhan, A. (2011) Extraction of genomic DNA from yeasts for PCR-based applications. *BioTechniques*, **50**, 325-328.
5. Inoue, H., Nojima, H. and Okayama, H. (1990) High efficiency transformation of *Escherichia coli* with plasmids. *Gene*, **96**, 23-28.
6. Lee, M.E., DeLoache, W.C., Cervantes, B. and Dueber, J.E. (2015) A Highly Characterized Yeast Toolkit for Modular, Multipart Assembly. *ACS Synth. Biol.*, **4**, 975-986.
7. Mans, R., van Rossum, H.M., Wijsman, M., Backx, A., Kuijpers, N.G., van den Broek, M., Daran-Lapujade, P., Pronk, J.T., van Maris, A.J. and Daran, J.M. (2015) CRISPR/Cas9: a molecular Swiss army knife for simultaneous introduction of multiple genetic modifications in *Saccharomyces cerevisiae*. *FEMS Yeast Res.*, **15**.
8. Kuijpers, N.G., Solis-Escalante, D., Luttik, M.A., Bisschops, M.M., Boonekamp, F.J., van den Broek, M., Pronk, J.T., Daran, J.-M. and Daran-Lapujade, P. (2016) Pathway swapping: Toward modular engineering of essential cellular processes. *Proc. Natl. Acad. Sci.*, 15060-15065.
9. Kuijpers, N.G., Solis-Escalante, D., Bosman, L., van den Broek, M., Pronk, J.T., Daran, J.-M. and Daran-Lapujade, P. (2013) A versatile, efficient strategy for assembly of multi-fragment expression vectors in *Saccharomyces cerevisiae* using 60 bp synthetic recombination sequences. *Microb. Cell Fact.*, **12**, 47.
10. Salazar, A.N., Gorter de Vries, A.R., van den Broek, M., Wijsman, M., de la Torre Cortes, P., Brickwedde, A., Brouwers, N., Daran, J.G. and Abeel, T. (2017) Nanopore sequencing enables near-complete de novo assembly of *Saccharomyces cerevisiae* reference strain CEN.PK113-7D. *FEMS Yeast Res.*, **17**.
11. Li, H. and Durbin, R. (2009) Fast and accurate short read alignment with Burrows-Wheeler transform. *Bioinformatics*, **25**, 1754-1760.
12. Thorvaldsdottir, H., Robinson, J.T. and Mesirov, J.P. (2013) Integrative Genomics Viewer (IGV): high-performance genomics data visualization and exploration. *Brief. Bioinform.*, **14**, 178-192.
13. Nijkamp, J.F., van den Broek, M.A., Geertman, J.M., Reinders, M.J., Daran, J.M. and de Ridder, D. (2012) De novo detection of copy number variation by co-assembly. *Bioinformatics*, **28**, 3195-3202.
14. Walker, B.J., Abeel, T., Shea, T., Priest, M., Abouelliel, A., Sakthikumar, S., Cuomo, C.A., Zeng, Q., Wortman, J., Young, S.K. *et al.* (2014) Pilon: an integrated tool for comprehensive microbial variant detection and genome assembly improvement. *PloS one*, **9**, e112963.
15. Postma, E., Verduyn, C., Scheffers, W.A. and Van Dijken, J.P. (1989) Enzymic analysis of the crabtree effect in glucose-limited chemostat cultures of *Saccharomyces cerevisiae*. *Appl. Environ. Microbiol.*, **55**, 468-477.
16. Jansen, M.L., Diderich, J.A., Mashego, M., Hassane, A., de Winde, J.H., Daran-Lapujade, P. and Pronk, J.T. (2005) Prolonged selection in aerobic, glucose-limited chemostat cultures of *Saccharomyces cerevisiae* causes a partial loss of glycolytic capacity. *Microbiology*, **151**, 1657-1669.

17. Cruz, L.A., Hebly, M., Duong, G.H., Wahl, S.A., Pronk, J.T., Heijnen, J.J., Daran-Lapujade, P. and van Gulik, W.M. (2012) Similar temperature dependencies of glycolytic enzymes: an evolutionary adaptation to temperature dynamics? *BMC Syst. Biol.*, **6**, 151.
18. Lowry, O.H., Rosebrough, N.J., Farr, A.L. and Randall, R.J. (1951) Protein measurement with the Folin phenol reagent. *J. Biol. Chem.*, **193**, 265-275.
19. Swiat, M.A., Dashko, S., den Ridder, M., Wijsman, M., van der Oost, J., Daran, J.-M. and Daran-Lapujade, P. (2017) FnCpf1: a novel and efficient genome editing tool for *Saccharomyces cerevisiae*. *Nucleic Acids Res.*, **45**, 12585-12598.
20. Mitchell, L.A. and Boeke, J.D. (2014) Circular permutation of a synthetic eukaryotic chromosome with the telomerator. *Proc. Natl. Acad. Sci.*, **111**, 17003-17010.
21. DiCarlo, J.E., Norville, J.E., Mali, P., Rios, X., Aach, J. and Church, G.M. (2013) Genome engineering in *Saccharomyces cerevisiae* using CRISPR-Cas systems. *Nucleic Acids Res.*, **41**, 4336-4343.
